# Linkage disequilibrium under polysomic inheritance

**DOI:** 10.1101/2020.12.07.415414

**Authors:** Kang Huang, Derek W. Dunn, Wenkai Li, Dan Wang, Baoguo Li

## Abstract

The influence of genetic drift on linkage disequilibrium in finite populations has been extensively studied in diploids. However, to date the effects of ploidy on LD has not been extensively studied. We here extend the linkage disequilibrium measure *D* and Burrow’s Δ statistic to include polysomic inheritance, as well as their corresponding squared correlation coefficients *r*^2^ and 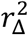, where the former is for phased genotypes and the latter for unphased genotypes. Weir & Hill’s double non-identity framework is also extended to include polysomic inheritance, and the expressions of double non-identity coefficients are derived under five mating systems. On this basis, the approximated expectations of estimated *r*^2^ and 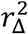 at equilibrium state, *d*^2^ and *δ*^2^, are derived under five mating systems. We assess the behaviors of the estimated *r*^2^ and 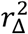 and the influence of the recombination rate on *d*^2^ or *δ*^2^, simulate the application of estimating effective population size, and evaluate the statistical performance of the method of estimating.

## Introduction

Linkage disequilibrium (LD) is the non-random association of alleles at different loci in a given population (Slatkin 2008), and can be influenced by many factors, such as selection, mutation, recombination, genetic drift, and the mating system (Nei 1987). Linkage disequilibrium can be measured by several parameters, such as the correlation coefficient r, Lewontin’s (1964) *D*’, Hill’s (1975) *Q*, Maruyama’s (1982) *D*\*, Ohta’s (1980) *F*\*, and Brown *et al.’s* (1980) *χ*. The most frequently used measure of LD is the squared correlation coefficient *r*^2^ (Hill and Weir 1994), which is the weighted sum of the squared correlation coefficient between alleles at two loci.

The influence of genetic drift on linkage disequilibrium in finite populations has been extensively studied in diploids (Hill and Robertson 1968; Ohta and Kimura 1969; Sved and Feldman 1973; Hill 1974; Weir 1979; Weir and Cockerham 1979; Weir and Hill 1980). In general, this previous work has shown that the squared correlation coefficient *r*^2^ (or 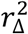 for unphased genotypes) will converge to a constant after several generations of random mating. This constant is determined by the recombination rate, effective population size and the mating system. Based on these factors, LD has been incorporated into two major applications: (i) gene mapping (Hästbacka *et al.* 1992; Hill and Weir 1994; Devlin and Risch 1995; Jorde 1995; Hosking *et al.* 2002) and (ii) the estimation of effective population size (Hill 1981; Hayes *et al.* 2003; England *et al.* 2006; Sved *et al.* 2013; Waples *et al.* 2014).

Many plant species are polyploid, with anywhere between 30-80% of angiosperm species being at least partially polyploid (Burow *et al.* 2001), with evidence for paleo-polyploidy in most plant lineages (Otto 2007). Polyploidy is also a major factor in the evolution of both wild and cultivated plants, and plays a key role in plant breeding (Udall and Wendel 2006; Sattler *et al.* 2016). However, to date the effects of ploidy on LD has not been extensively studied.

There are at least three typical features in polysomic inheritances: (i) the chromosomes are randomly paired and exchange their chromatid segments during meiosis, in which the recombination rate *c* is 1 — 1/*ν* if the corresponding loci are located on different chromosomes (*ν* is the ploidy level), or ≤ 0.5 if the corresponding loci are located on the same chromosome; (ii) the decay coefficient of heterozygosity (i.e. the ratio of single non-identity coefficients in the next and the current generations) is 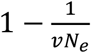 in polyploids (*N_e_* is the effective population size); (iii) the multivalents may be formed during meiosis (Rieger *et al.* 1968), resulting in a particular phenomenon in polysomic inheritance, termed the double-reduction (Butruille and Boiteux 2000), in which a gamete may inherit a single gene copy twice.

In this paper, we extend both the linkage disequilibrium measure *D* and Burrow’s Δ statistic to account for polysomic inheritance, and calculate their corresponding squared correlation coefficients *r^2^* and 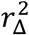. We also extend Weir & Hill’s (1980) double non-identity framework to account for polysomic inheritance, and derive the expressions of these double non-identity coefficients under five mating systems. On this basis, we are able to derive 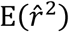 and 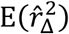 at equilibrium state, and these two expectations are approximated by *d*^2^ and *δ*^2^, respectively. Both approximates are closely related to the mating system together with the effective population size *N_e_* and the recombination rate *c*. We study the behavior of the squared correlation coefficient estimators 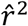 and 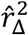 during genetic drift, investigate the influence of recombination rate *c* on *d*^2^ or *δ*^2^, simulate the application for estimating effective population size *N_e_*, and evaluate the statistical performance of estimating 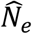. We discuss the relationship between *r^2^* and *c* (or between 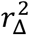 and *c*), and that between *r*^2^ and *ν* (or between 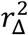 and *ν*). We enable the estimation of Burrow’s Δ, the testing of linkage disequilibrium based on Burrow’s Δ, and the estimation of effective population size in our software package Polygene V1.3 (Huang *et al.* 2020), which is freely available via http://github.com/huangkang1987/polygene.

## Theory and modelling

### LD measurements

We denote *A* and *B* for two alleles each from a unique locus. The generalized LD measurement *D* between *A* and *B* is defined as the difference between the observed and the expected frequencies of the haplotype *AB.* We model five specific variants of *D:* (i) 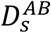 (for the same haplotype), (ii) 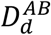 (for two different haplotypes within the same individual), (iii) 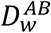 (for the within-individual component), (iv) 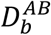 (for the between-individual component) and (v) *D_AB_* (for the usual LD measurement). These measurements can be defined by symbols as follows:

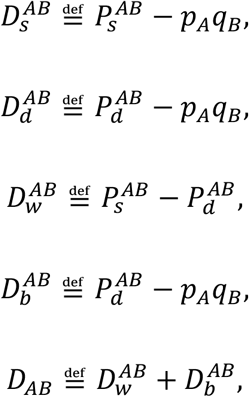

where 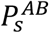 is the probability that the alleles in the same haplotype are *A* and *B*, 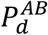 is the probability that alleles in different haplotypes within the same individual are *A* and *B,* and *p_A_* and *q_B_* are respectively the frequencies of *A* and *B,* respectively.

According to these definitions, the following expressions hold:

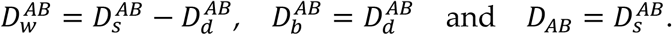

The usual LD measurement *D_AB_* is the covariance between *A* and *B* in the same haplotype, i.e. 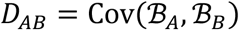, where 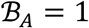 if the first allele copy in the target allele pair is *A*, otherwise 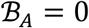, and the meaning of 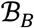 is analogous.

The values of *D_AB_* may be negative, and its range is influenced by the frequencies of *A* and *B*. It is therefore more intuitive to use Pearson’s correlation coefficient *r_AB_* to measure LD, which is the quotient of *D_AB_* divided by its theoretical maximum 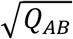. In other words, 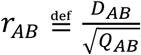, whose range is [-1,1], where *Q_AB_* = *p_A_p_X_q_B_q_X_* (*X* represents any allele distinct from both *A* and *B*, and thus *p_x_* = 1 — *p_A_* and *q_x_* = 1 — *q_B_*). Note that 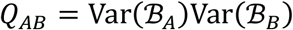. We obtain

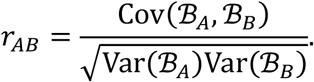

The values of *r_AB_* may also be negative. However, the squared correlation coefficient 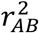 ensures that its values are non-negative, ranging from 0 to 1. We will adopt the average value of 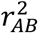 across all allele pairs to evaluate the LD between two loci for the situation of phased genotypes.

The above LD measurements are applicable for phased genotypes although unphased genotypes are more common. For unphased genotypes, Burrows’s Δ statistic (Cockerham and Weir 1977) can be used, and we will extend this to account for polysomic inheritance. By using 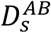 and 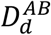, Burrows’s *Δ* statistic between *A* and *B* can be defined as 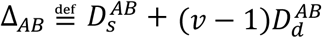. Moreover, for two-locus unphased genotypes, Burrow’s Δ statistic can be expanded to:

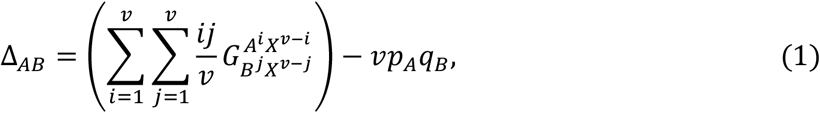

where *X* is an arbitrary allele distinct from both *A* and *B*, with each 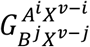 denoting a two-locus unphased genotypic frequency whose superscript (or subscript) is an unphased genotype containing exactly *i* copies of *A* (or *j* copies of *B*). In Appendix A, we use triploids to illustrate how Δ_*AB*_ is expanded. Substituting the observed values of *p_A_, p_X_, q_B_, q_X_* and 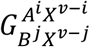 into Equation (1), Δ_*AB*_ can be estimated.

Similarly, the Pearson’s correlation coefficient *r*_Δ*AB*_ is the standardization of Δ_*AB*_. Where 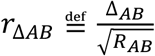, and 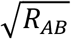 is the theoretical maximum of Δ_*AB*_. Because 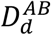 is determined by both LD and inbreeding, 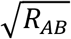 is defined as the value of Δ_*AB*_ under both complete LD and the current degree of inbreeding, denoted by Δ_*AB,complete*_. In this case, if *p_A_ = q_B_*, the corresponding 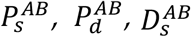 and 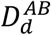 can be calculated by the following formulas:

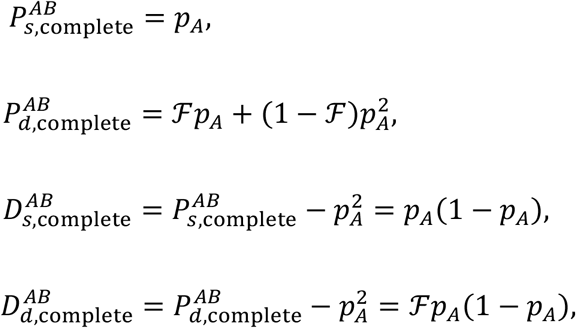

where 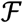 is the inbreeding coefficient. Therefore

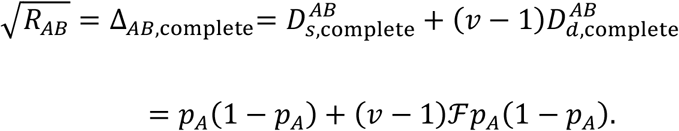

Let *P_AA_* be the probability of sampling two copies of *A* within the same individual without replacement. Then 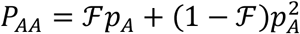, and so 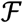 can be calculated by

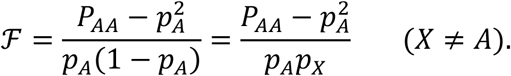

Substituting the expression of 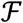 into that of 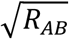, a simplified expression of 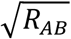 for the case of *p_A_* = *q_B_* is obtained, which is

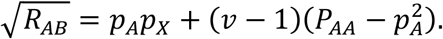

Next, if *p_A_* ≠ *q_B_*, we let 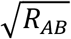 be the following geometric mean:

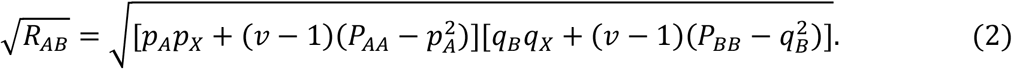

Likewise, *r*_Δ*AB*_ may be negative, but the squared correlation coefficient 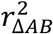 ranges from 0 to 1, which can also be used to evaluate the LD between two loci for unphased genotypes.

In the following text, for simplicity, we will use *D_w_, D_b_, D*, Δ, *Q, R, r* and *r*_Δ_ to replace 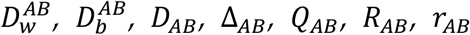 and *r*_Δ*AB*_ in turn. Due to genetic drift, *D*^2^ and *Q* (or Δ^2^ and *R*) converge to zero after an infinite number of generations. However, the ratio *r*^2^ of *D*^2^ to *Q* (or the ratio 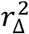 of Δ^2^ to *R*) converges to a constant, whose value is determined by the mating system together with the recombination rate *c* and the effective population size *N_e_* (Weir and Hill 1980). In the following sections, we extend Weir and Hill’s (1980) double non-identity framework, to obtain the expectations of *r*^2^ and 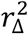.

### Double non-identity coefficients

The double non-identity coefficients can be used to derive the moments of various LD measurements and to approximate 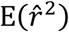 and 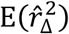 at equilibrium state. The term *identity* means identical-by-descent (IBD).

The single non-identity coefficient is defined as the probability that the two alleles of an allele pair are not IBD. There are two configurations for two such alleles: (i) they originate from the same individual, or (ii) they originate from different individuals. We denote the single non-identity coefficient by *P* for (i), or by Π for (ii). Then, *P* and Π can be described by symbols as follows:

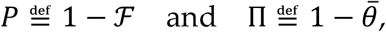

where 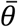 is the average kinship coefficient between all individuals in a population, i.e. the probability that two alleles (each randomly sampled from a separate individual) are IBD.

The double non-identity coefficient is defined as the probability that neither of two allele pairs are IBD. There are multiple configurations for these two allele pairs. Based on Weir & Hill (1980), we establish 3 digenic, 6 trigenic and 13 quadgenic two-locus allele configurations for different polysomic inheritances, including the following four novel allele configurations (9^th^, 15^th^, 21^st^ and 22^nd^) that have more than two haplotypes within individuals. These allele configurations along with the notations of the corresponding frequencies, double non-identity coefficients, and expectations are presented in Table 1, where the first nine allele configurations do not have corresponding double non-identity coefficients because they share the same alleles.

**Table 1.**
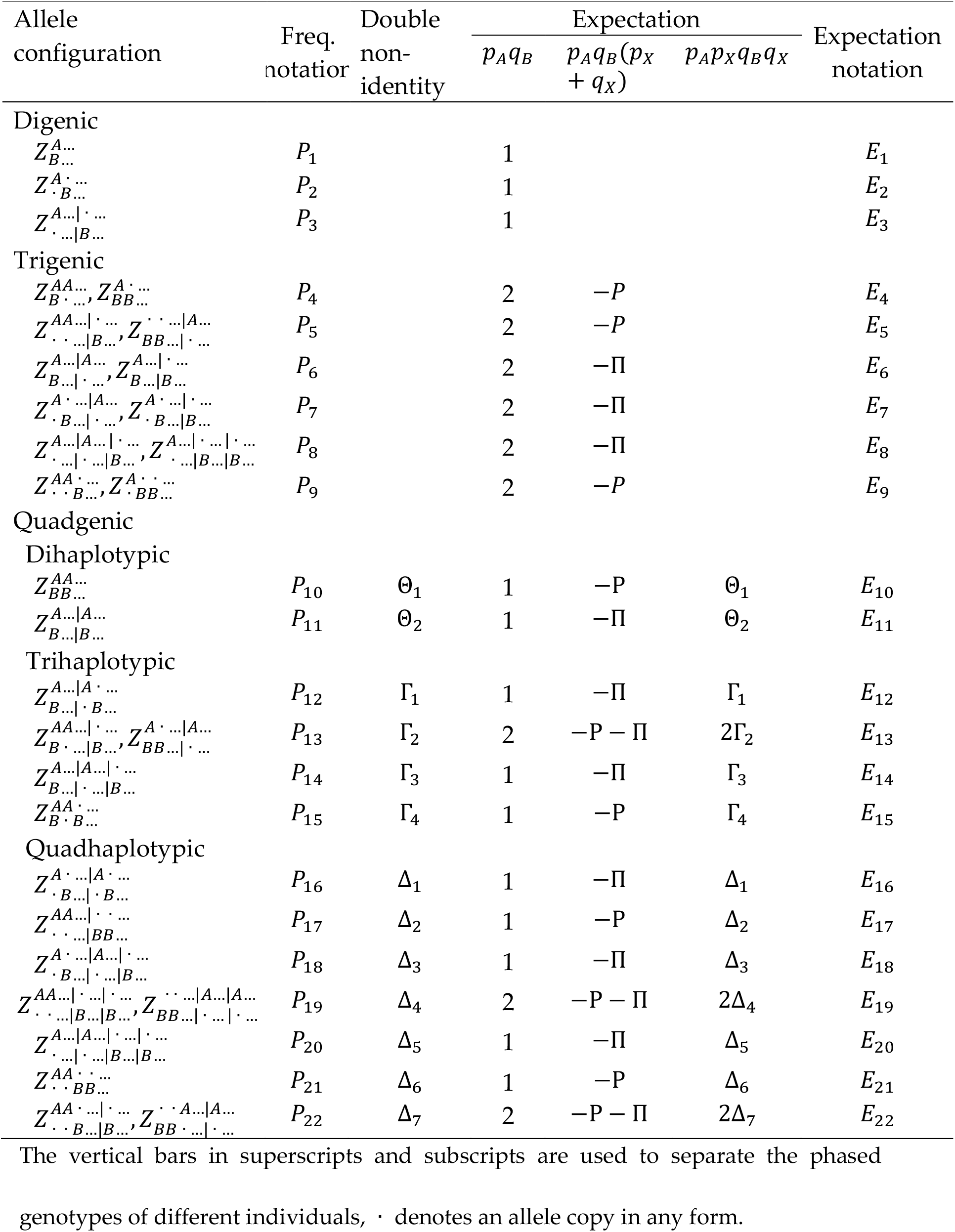
Allele configurations and their expected frequencies

For example, the 10^th^ allele configuration 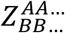 in Table 1 means that these two allele pairs are from two haplotypes within the same individual, the first *A* and first *B* are in one haplotype, and the second *A* and second *B* are in another haplotype. Moreover, the corresponding frequency, double nonidentity coefficient and the expectation of frequency are denoted by *P*_10_, and *E*_10_, respectively.

The expectation *E_i_* of each frequency *P_i_* in Table 1 is derived by assuming no initial LD, which is a linear combination of *p_A_q_B_, p,_A_q_B_*(*p_X_* + *q_X_*) and *p_A_p_X_q_B_q_X_*, whose combination coefficients are listed in the three cells before *E_i_* in Table 1. For example, the combination coefficients of *E*_18_ are 1, —Π and Δ_3_. The allele pair *AA* or *BB* in the 18^th^ allele configuration 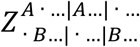 consists of the alleles from different individuals, then the single non-identity coefficient of each allele pair is Π and the double non-identity coefficient is Δ_3_. Hence the identity states of these two allele pairs can be described by the following three aspects: (i) both pairs are non-IBD at the probability Δ_3_, (ii) only one pair is IBD at the probability Π — Δ_3_ or (iii) both pairs are IBD at the probability 1 — 2Π + Δ_3_. Therefore, the expectation *E*_18_ is the following linear combination with 1, —Π and Δ_3_ as the combination coefficients:

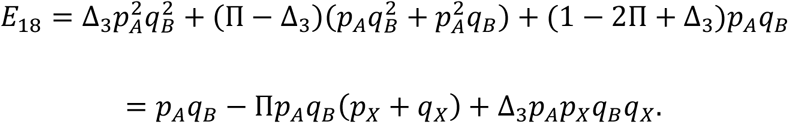

Various moments of LD measurements can be derived by using the expectations *E*_1_ to *E*_22_, with the derivations presented in Appendix B. Because these expectations are functions of both the single and the double nonidentity coefficients, by using Table 1, each moment in Appendix B can be converted to a function of double non-identity coefficients (two single nonidentity coefficients *P* and Π are accommodated into double non-identity coefficients), whose essential factors are listed in Table 2.

**Table 2.** Essential factors of moment expressions

| | $E(\bar{D}_w^2)$ | $E(\bar{D}_b^2)$ | $E(\bar{D}_w \bar{D}_b)$ | $E(\bar{D}^2)$ | $E(\bar{\Delta}^2)$ | $E(\bar{Q})$ | $E(\bar{R})$ | Divisor |
| --- | --- | --- | --- | --- | --- | --- | --- | --- |
| $\Theta_1$ | $n^2 v^2 \times$<br>$(2 + v v_2)$ | $\lambda_7$ | $-n v \times$<br>$(v v_1 + \lambda_1)$ | $v_1^2 \lambda_5$ | $2 v^2 n_1^2 v_1^2$ | $2 v_1^2$ | $2 v^2 n_1^2 v_1^2$ | $n^3 v^3 v_1 / Q$ |
| $\Theta_2$ | $n^2 v^2$ | 2 | $-n v$ | $\lambda_5$ | $v^2 \lambda_2$ | 2 | $2 v^2$ | $n^3 v^2 / n_1 Q$ |
| $\Gamma_1$ | $-2 n^2 v^2$ | $-2 \lambda_1$ | $n v (1 + \lambda_4)$ | $-2 v_1 \lambda_3$ | $2 v^2 v_1 \lambda_2$ | $4 v_1$ | $4 v^2 v_1$ | $n^3 v^2 / n_1 Q$ |
| $\Gamma_2$ | 0 | $-2 \lambda_1$ | $-n v v_2$ | $-2 v_1 \lambda_3$ | $-4 v^2 n_1 v_1$ | $4 v_1$ | $-4 v^2 n_1 v_1$ | $n^3 v^2 / 2 n_1 Q$ |
| $\Gamma_3$ | 0 | 4 | $-n v$ | $-2 \lambda_3$ | $-2 v^2 n_2$ | 4 | $4 v^2$ | $n^3 v / n_1 n_2 Q$ |
| $\Gamma_4$ | $-2 n^2 v^2 v_2$ | $2 \lambda_1 \lambda_4$ | $\lambda_6$ | $-2 v_1^2 \lambda_3$ | $4 v^2 n_1^2 v_1^2$ | $4 v_1^2$ | $4 v^2 n_1^2 v_1^2$ | $n^3 v^3 v_1 / v_2 Q$ |
| $\Delta_1$ | $n^2 v^2$ | $\lambda_7$ | $-n v \lambda_4$ | $2 v_1^2$ | $v^2 v_1^2 \lambda_2$ | $2 v_1^2$ | $2 v^2 v_1^2$ | $n^3 v^2 / n_1 Q$ |
| $\Delta_2$ | 0 | 1 | 0 | 1 | $v^2$ | 1 | $v^2 n_1^2$ | $n^3 v^2 / n_1 v_1^2 Q$ |
| $\Delta_3$ | 0 | $-2 \lambda_1$ | $n v$ | $4 v_1$ | $-2 v^2 n_2 v_1$ | $4 v_1$ | $4 v^2 v_1$ | $n^3 v / n_1 n_2 Q$ |
| $\Delta_4$ | 0 | $v_1$ | 0 | $v_1$ | $v^2 v_1$ | $v_1$ | $-v^2 n_1 v_1$ | $n^3 v / 2 n_1 n_2 Q$ |
| $\Delta_5$ | 0 | 1 | 0 | 1 | $v^2$ | 1 | $v^2$ | $n^3 / n_1 n_2 n_3 Q$ |
| $\Delta_6$ | $n^2 v^2$ | $\lambda_4^2$ | $-n v \lambda_4$ | $v_1^2$ | $v^2 n_1^2 v_1^2$ | $v_1^2$ | $v^2 n_1^2 v_1^2$ | $n^3 v^3 v_1 / v_2 v_3 Q$ |
| $\Delta_7$ | 0 | $-2 \lambda_4$ | $n v$ | $2 v_1$ | $-2 v^2 n_1 v_1$ | $2 v_1$ | $-2 v^2 n_1 v_1$ | $n^3 v^2 / 2 n_1 v_2 Q$ |

For brevity, we denote $n - i$ by $n_i$ and $v - i$ by $v_i$ , and let $\lambda_1 = n_2 v + 2$ , $\lambda_2 =$ $n n_2 + 2$ , $\lambda_3 = n v - 2$ , $\lambda_4 = v n_1 + 1$ , $\lambda_5 = 2 + n v \lambda_3$ , $\lambda_6 = n v [2 + (\lambda_3 - \lambda_4)(1 + \lambda_4)]$ , $\lambda_7 = 2 + 2 n_2 v + \lambda_2 v^2$ .

The expressions of various moments can now be expressed uniformly by matrices. Let **M** be the row vector consisting of the 7 moments in the header row of Table 2, and let **Φ** be the column vector consisting of the 13 double non-identity coefficients in the header column of Table 2. Denote A as a 13 × 7 matrix, whose *i*^th^ column consists of the *i*^th^ column divided by the last column of Table 2. Then

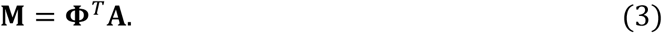

We call **M** the *moment vector,* and **Φ** the *double non-identity vector.* Moreover, A can be decomposed as the following combination:

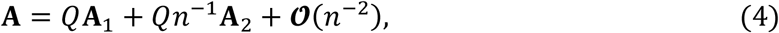

if we let

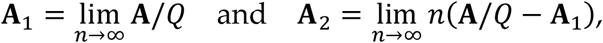

where the elements in the combination matrices **A**_1_ and **A**_2_ in the principal part of **A** are listed in Tables S1 and S2, respectively.

Finally, if the correlation between 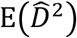 and 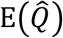 is not considered, the ratio *d*^2^ of 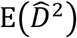 to 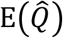 can be used as an approximation of the moment 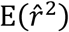. Similarly, the ratio *δ*^2^ of 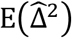 to 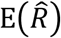 can also be used as that of 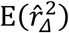 if the correlation between 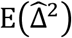 and 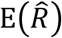 is not considered. These facts can be denoted by symbols as follows:

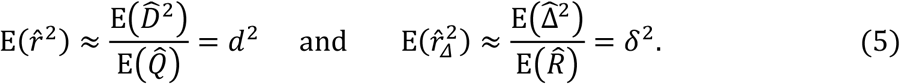

### Transition matrix of double non-identity coefficients

The transition matrix of double non-identity coefficients can be used to describe the behavior of double non-identity coefficients due to genetic drift.

Let **Φ** be the double non-identity vector in the current generation, and let **Φ**′ be that in the next generation. If **Φ** is given, then **Φ**′ can be calculated from **Φ**. We use a diploid Wright-Fisher population with *N* individuals to demonstrate how to calculate **Φ**′. For this population, we have Θ_1_ = Θ_2_, Γ_1_, = Γ_2_ = Γ_3_ and Δ_1_, = Δ_2_ = Δ_3_ = Δ_4_ = Δ_5_, but Γ_4_, Δ_6_ and Δ_7_ are undefined. Therefore, both **Φ** and *Φ*’ contain 10 elements, and there are only three distinct elements in **Φ** or *Φ’*. Because the individuals are formed by the random unison of gametes, Θ_1_ in **Φ** can be updated to 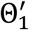 in **Φ**’, in the sense that

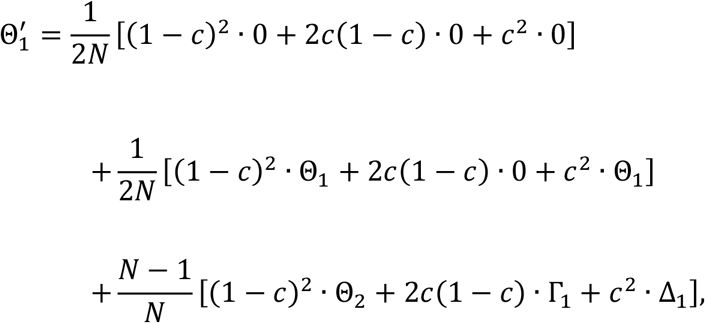

where *c* is the recombination rate between the two given loci. Similarly, the other two elements 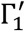, and 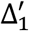, in **Φ**’ can also be calculated. It can be shown that each element in **Φ**’ can be expressed as a linear combination in the form of *a*Θ_1_, + *b*Γ_1_ + *c*Δ_1_.

In general, each updated double non-identity coefficient can also be expressed as a linear combination of the current double non-identity coefficients. If **Ω** denotes the matrix consisting of those combination coefficients, then **Φ**’ can be expressed as **Φ**’ = **ΩΦ**. We call **Ω** the *transition matrix* from **Φ** to **Φ**’.

Let **Φ**_0_ be the double non-identity vector in the founder generation and let **Φ**_*t*_ be that in the *t*^th^ generation. This gives *t*_th_ = **Ω**^*t*^**Φ**_0_. If a population is allowed to reproduce for several generations, the vector sequence is:

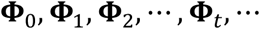

and will reach a steady state as *t* increases. In other words, this sequence will converge to a constant vector, denoted by **Φ**_∞_. This limit vector **Φ**_∞_ is independent to the initial vector **Φ**_0_ if **Φ**_0_ ≠ ***0***.

To simplify the model for polysomic inheritance, we established a virtual mating system, named the *haplotype sampling* (HS) *mating system.* In this mating system, it is assumed that each individual is reproduced by randomly sampling *v* haplotypes with replacement from the previous generation. The genes in an offspring therefore come from a maximum of *v* parents. Because the haplotypes within (or among) individuals are randomly sampled, there is no difference among dihaplotypic, trihaplotypic and quadhaplotypic double non-identity coefficients, symbolically Θ_1_ = Θ_2_, Γ_1_ = Γ_2_ = Θ_3_ = Θ_4_ and Δ_1_ = Δ_2_ =…. = Δ_7_. Therefore, for this mating system, the transition matrix **Ω** can be simplified as a 3 × 3 matrix, whose transposition **Ω**^*T*^ is listed in Table S3 and derivation is given in Appendix C. The transposition of a simplified transition matrix (still denoted by **Ω**) is presented in Table 3, and its element expressions can be easily exported from this table. In the following discussion, we will also use Table 3 to investigate the behavior of LD in polyploids.

**Table 3.** Simplified Ω^*T*^ for HS mating system

| 1 | $N^{-1}$ | 1 | $N^{-1}$ | $N^{-2}$ | 1 | $N^{-1}$ | $N^{-2}$ | $N^{-3}$ |
| --- | --- | --- | --- | --- | --- | --- | --- | --- |
| $(c - 1)^2$ | $\frac{c^2}{v - 1} - \frac{1 + 2c(c - 1)}{v}$ | 0 | $\frac{1 - c}{v}$ | $\frac{2c - 1}{v^2}$ | 0 | 0 | $\frac{2}{v^2}$ | $-\frac{2}{v^3}$ |
| $2c(1 - c)$ | $\frac{4c(2c - 1)}{v} - \frac{2c^2}{v - 1}$ | $1 - c$ | $\frac{6c - 3}{v}$ | $\frac{2 - 8c}{v^2}$ | 0 | $\frac{4}{v}$ | $-\frac{12}{v^2}$ | $\frac{8}{v^3}$ |
| $c^2$ | $\frac{2c^2(3 - 2v)}{v(v - 1)}$ | $c$ | $-\frac{5c}{v}$ | $\frac{6c}{v^2}$ | 1 | $-\frac{6}{v}$ | $\frac{11}{v^2}$ | $-\frac{6}{v^3}$ |

Each element of $\Omega^T$ is a combination of 1, $N^{-1}$ , $N^{-2}$ and $N^{-3}$ with the combination coefficients in the corresponding cell. The combination coefficients are zero for unrepresented terms.

It is noteworthy that the sum of the combination coefficients of 1 in each column in Table 3 is exactly one, but the sum of the elements in each column of **Ω**^*T*^ is less than one. This indicates that the transition (i.e. a generation of random mating) will gradually reduce the double-nonidentity coefficients, and their values will be eventually converge to zero, i.e. **Ω**^∞^ = ***0***. This illustrates the loss of heterozygosity and the fixation of alleles.

Although **Φ**_∞_ will eventually converge to zero, the ratio of the moments 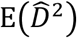 to 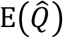, and of the moments 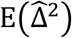 to 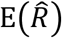 will converge to some constants. This can be considered as the double non-identity vector **Φ** reaches a relatively stable state so the direction of **Φ** is constant during reproduction, symbolically **Φ**’ = *ν***Φ**. The direction of **Φ** (say *ω*) and the scale factor *v* can be solved by performing eigen-value decomposition for *Ω,* i.e. solving *Ωω* = *ν**ω***, which means *ν* is an eigenvalue of **Ω**, and ***ω*** is an eigenvector of **Ω** belonging to *ν*. It is also noteworthy that there are multiple eigenvalues, with the highest eigenvalue be of our interest. Therefore, *d*^2^ and *δ*^2^ can be calculated from the *moment vector* calculated from Equation (3) by substituting **Ω** with ***ω***, i.e. **M**_*ω*_ = ***ω***^*T*^**A**. We denote the elements in **M**_*ω*_ as E_*ω*_(·), e.g. 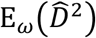, then the exact *d*^2^ and *δ*^2^ are as follows:

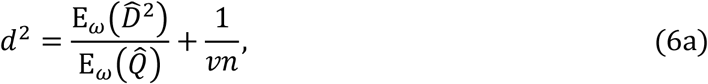

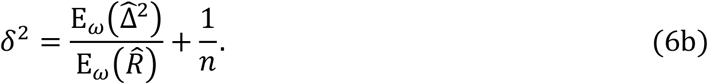

The additional terms 1/*vn* and 1/*n* are used to include any error resulting from finite sample size (see Appendix D for details)

### Approximations of *d*^2^ and *δ*^2^

To simplify the calculation and to facilitate the discussion of the properties of *d*^2^ and *δ*^2^, Weir & Hill (1980) adopted a matrix decomposition technique to approximate *ν* and *ω* for disomic inheritance and also to approximate *d*^2^ and *δ*^2^. Here, we will follow this approach to derive the approximate expressions of *d*^2^ and *δ*^2^ for the HS mating system and four additional mating systems.

Let **Ω** be the simplified transition matrix for the HS mating system, as detailed in Table 3. If *N* is large enough, the values of the terms with *N*^-2^ and *N*^-3^ in Table 3 will be small, then **Ω** can be decomposed to:

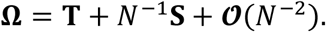

For the matrices **T** and **S** in the principal part of **Ω**, if we denote *c_i_* = *c* — *i* and *ν_i_* = *ν* — *i* (*i* = 1,2), we will obtain from Table 3:

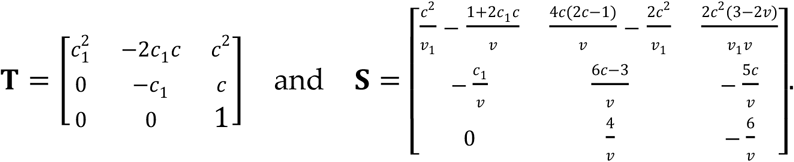

Similarly, *ν* and ***ω*** can be decomposed to

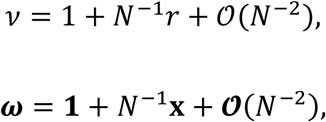

where **1** = [1,1,1]^*T*^ and **x** = [*x*_1_, *x*_2_, *x*_3_]^*T*^. According to ***Ωω*** = ***νω***, we obtain a matrix equation as follows:

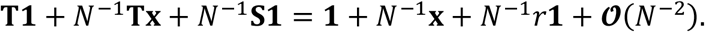

Because **T1** = **1**, if the term 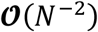 is omitted, we obtain

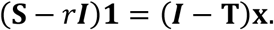

This matrix equation is an approximation of the previous equation, which is a linear equation set with 3 equations and 4 unknowns, the solutions of which are as follows:

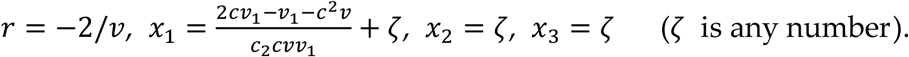

If we let *ζ* = 0, we obtain a special solution: *r* = — 2/*v* and 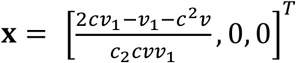. Replacing this solution into the expressions of *ν* and ***ω***, it follows that:

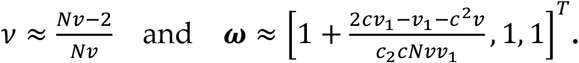

Now, by substituting ***ω*** in Equation (3) with the approximation of ***ω***, it can be calculated that

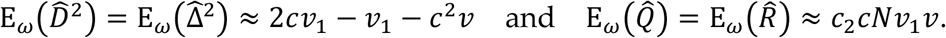

Therefore, the approximated *d*^2^ and *δ*^2^ are as follows:

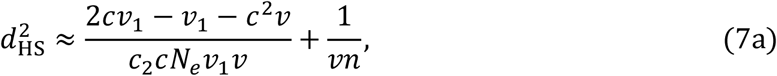

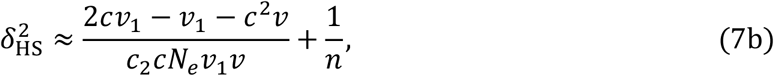

where *N_e_* and *N* are equivalent under the HS mating system. The results from Equations (7a) and (7b) accord with those of Ohta and Kimura (1969) and Weir & Hill (1980) for the monoecious selfing mating system in diploids.

The transition of single non-identity coefficients satisfies the relations: 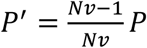 and 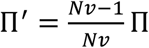. Moreover, if two loci are located at two extreme far locations on the same chromosome, and the thirteen double non-identity, coefficients are all equal to *P*^2^ and 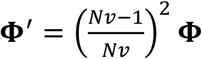, and thus also the corresponding eigenvalue 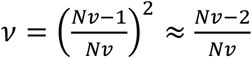. By comparing with the previous conclusion of 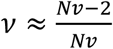 by substituting *ζ* = 0, we see that *r* = —2/*ν* is a good approximation to the rate of loss of heterozygosity at the pairs of independent loci.

We now follow the approach of Weir & Hill (1980) to establish four additional mating systems. Two are monecious mating systems: (i) selfing being allowed (termed MS), and (ii) selfing being excluded (termed ME). In both of these mating systems, the effective population size *N_e_* is the same as the population size *N*. The other two mating systems we use are both dioecious systems: (i) dioecious with random pairing (termed DR), and dioecious with lifetime pairing (termed DH). In DR, each offspring is produced from a new pairing. In DH, each individual remains in a single reproductive unit for its entire lifetime. Moreover, in both DR and the DH, there are *M* males and *F* females in the population for each generation and *F* = *fM*, the effective population size is calculated by 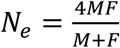.

The transition matrix **Ω** for each of the four additional mating systems is a 13 × 13 matrix, whose element expressions are derived in Appendices E to G. The results are not shown, but the matrices **T** and **S** in the principal part of **Ω** for all five mating systems (HS, MS, ME/DR and DH) are listed in Appendix H.

The equality **T1 = 1** still holds for all five mating systems. Therefore, the same approximations of *d*^2^ and *δ*^2^ can be used to derive the next matrix equation if *N_e_* is large enough:

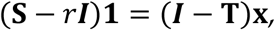

where **x** = [*x*_1_, *x*_2_, –, *x*_13_]^*T*^. From Appendix H, we show that the elements in the 11^th^ row of ***I*** — **T** are all zero, and the 11^th^ element in the vector (**S** — *r***I**)**1** is 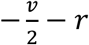 for all five mating systems. Therefore, the 11^th^ equation in this matrix equation is 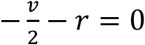, and thus *r* = —2/*ν*. For the other 13 unknowns *x*_1_, *x*_2_, –, *x*_13_, the 13 differences

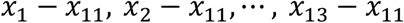

can be solved for each mating system. The appropriate expressions are listed in Table 4.

**Table 4.** Expressions of *x*_1_ — *x*_11_,*x*_2_ – *x*_11_, …, *x*_13_ – *x*_11_

|  | HS | MS | ME/ DR | DH |
| --- | --- | --- | --- | --- |
| $x_1 - x_{11}$ | $\frac{2c + c_1^2 v - 1}{c_2 c v_1 v}$ | * | * | * |
| $x_2 - x_{11}$ | $\frac{2c + c_1^2 v - 1}{c_2 c v_1 v}$ | * | * | * |
| $x_3 - x_{11}$ | 0 | $\frac{4c v_1 v_2}{v^2(c v_2 + v)(3v - 4)}$ | $\frac{4c v_1 v_2}{v^2(c v_2 + v)(3v - 4)}$ | $\frac{2c v_2(2f v_1 + 3v - 2)}{(1 + f)v^2(c v_2 + v)(3v - 4)}$ |
| $x_4 - x_{11}$ | 0 | $\frac{v_2}{v^2}$ | $\frac{2v_1}{v^2}$ | $\frac{2v_1}{v^2}$ |
| $x_5 - x_{11}$ | 0 | 0 | 0 | 0 |
| $x_6 - x_{11}$ | 0 | * | * | * |
| $x_7 - x_{11}$ | 0 | $\frac{v_2^2}{v_1 v^2(3v - 4)}$ | $\frac{v_2^2}{v_1 v^2(3v - 4)}$ | $\frac{f v_2^2 + 3v^2 - 6v + 4}{(1 + f)v_1 v^2(3v - 4)}$ |
| $x_8 - x_{11}$ | 0 | $\frac{2v_2}{v^2}$ | $\frac{4v_1}{v^2}$ | $\frac{4v_1}{v^2}$ |
| $x_9 - x_{11}$ | 0 | 0 | 0 | 0 |
| $x_{10} - x_{11}$ | 0 | $\frac{v_2}{v^2}$ | $\frac{2v_1}{v^2}$ | $\frac{2v_1}{v^2}$ |
| $x_{11} - x_{11}$ | 0 | 0 | 0 | 0 |
| $x_{12} - x_{11}$ | 0 | * | * | * |
| $x_{13} - x_{11}$ | 0 | $\frac{v_2}{v^2}$ | $\frac{2v_1}{v^2}$ | $\frac{2v_1}{v^2}$ |

The expressions represented by \* are too long and are placed in the supplementary files.

Let *a_i_*, = *x_i_* — *x*_11_ then *x_i_* = *a_i_*, + *x*_11_, *i* = 1,2, —, 13. The solutions of the matrix equation for each mating system can be expressed as

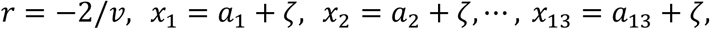

where *ζ* is any number. Now, if we let *ζ* be 0, we obtain a special solution as follows:

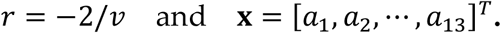

In addition, 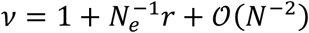 and 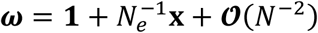, it follows

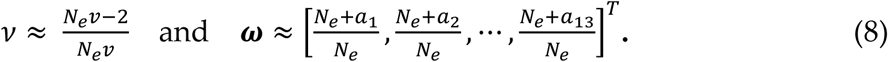

Similarly, by substituting the approximation of **ω** into Equation (3), we can calculate the approximations of various moments, and then annotate the approximate expressions of *d*^2^ and *δ*^2^ for four mating systems (MS, ME, DR and DH) as follows:

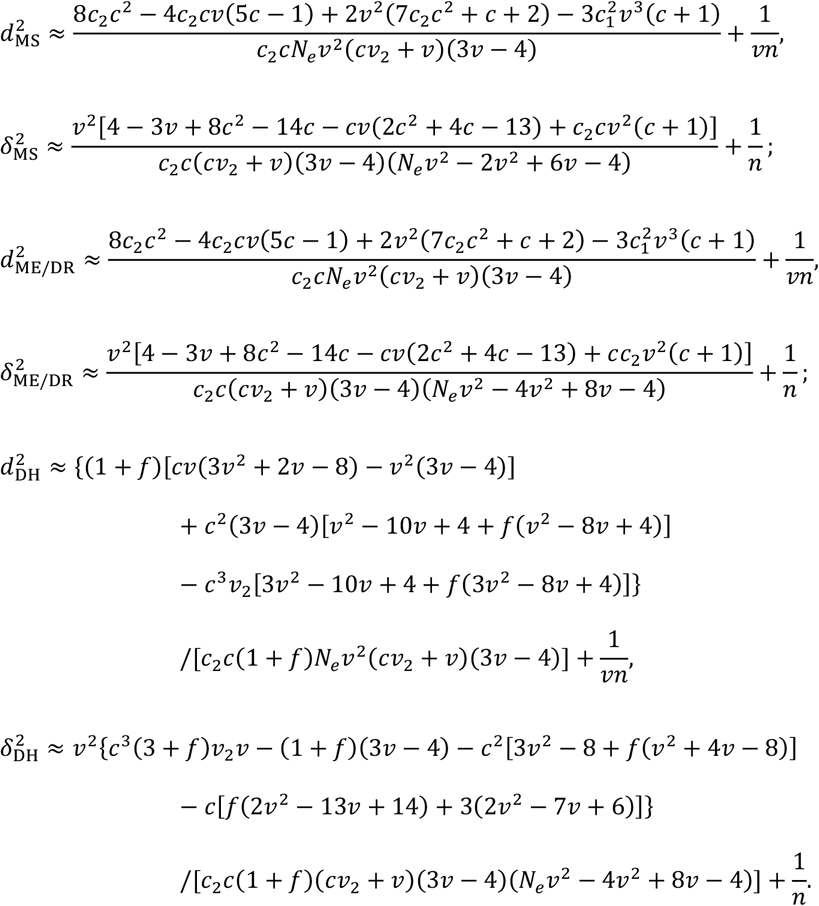

The approximate expressions of *d*^2^ and *δ*^2^ for each of the five mating systems for inheritances ranging from disomic to decasomic (with even ploidy levels) are presented in Tables S4 and S5. The corresponding values between unlinked loci located on either the same chromosome (*c* = 0.5) or different chromosomes (*c* = 1 — 1/*ν*) are presented in Table 5.

**Table 5.**
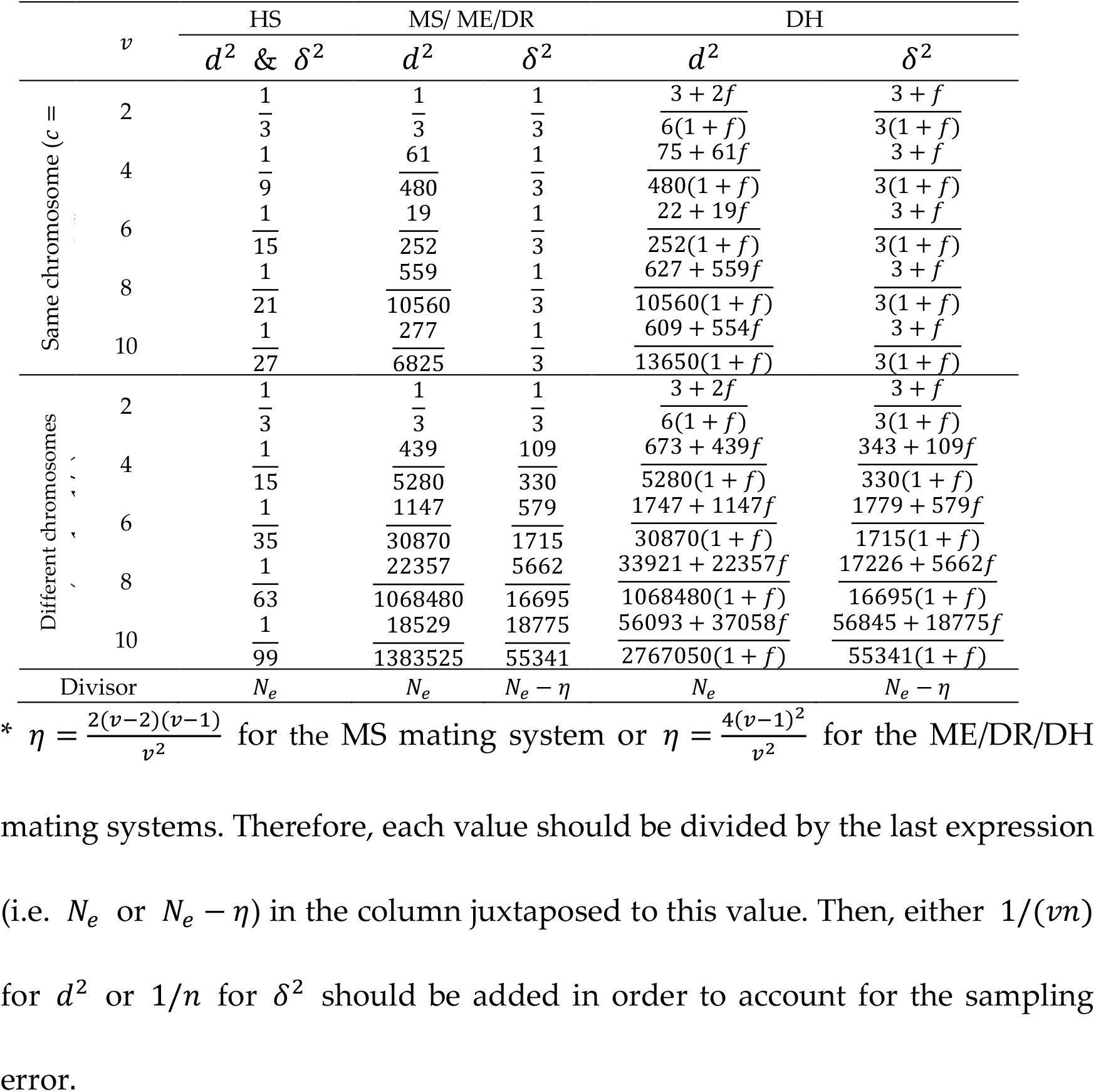
Approximations of *d*^2^ and *δ*^2^

### Simulations and evaluations

#### Behaviors of 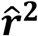 and 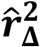

In this section, we discuss the behaviors of the squared correlation coefficient estimators 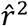 and 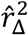 during reproduction and provide the exact and the approximate values of *d*^2^ or *δ*^2^ for reference.

Because the approximations of 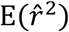 and 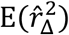 in Equation (5) are affected by the correlation between 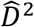 and 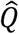 (or between 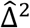 and 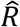), to eliminate such effects, we consider all values of 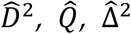 and 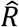, and let 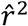 and 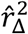 be the following ratios:

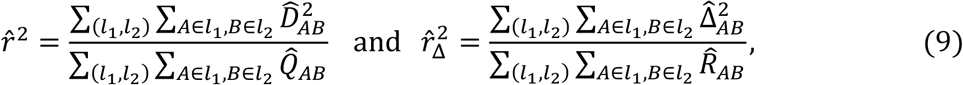

where (*l*_1_, *l*_2_) is taken from all diallelic locus pairs, the symbol *A* ∈ *l*_1_ (or *B* ∈ *l*_2_) represents *A* (or *B*) is taken from all alleles at the first (or the second) locus in (*l*_1_, *l*_2_).

We adopt a Monte-Carlo method to simulate the behavior of 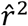 and 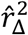. During simulation, a population with the MS mating system is generated, which contains 40 or 80 individuals with a ploidy level of either 2 or 4. Next, the individuals generated are genotyped at 200 linked diallelic locus pairs, with a recombination rate 0.1 for each locus pair. The population is then allowed to reproduce for 250 generations. For each generation, by using the data of genotypes of all individuals under various situations, 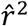 and 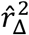 are calculated by Equation (9), and the exact and the approximate *d*^2^ and *δ*^2^ are also calculated by Equations (6) and (7), respectively. This process is performed 300,000 times in total. The results are shown in Figure 1.

**Figure 1.**
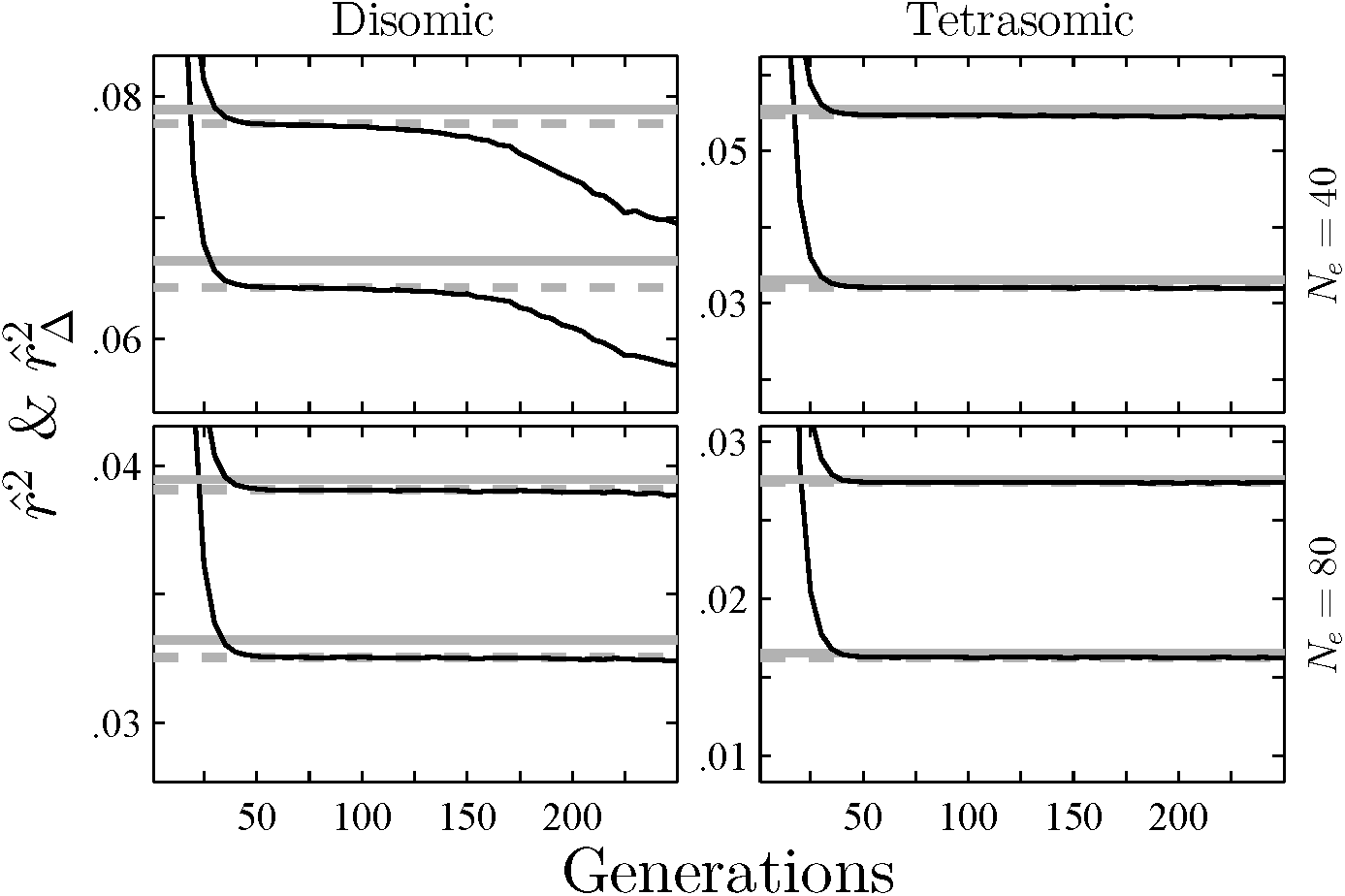
The behaviors of 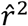 and 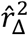 during reproduction for the MS mating system (set *N_e_* = 40 or 80, *v* = 2 or 4, *L* = 200 and *c* = 0.1). Each of the two columns shows the results of a different ploidy level, and each of the two rows shows the results of a different effective population size. Solid gray lines denote approximate *d*^2^ or *δ*^2^, dotted gray lines denote exact *d*^2^ or *δ*^2^, and solid lines denote 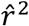 or 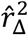 where the lines representing *δ*^2^ (or 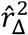) are above those representing *d*^2^ (or 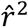) for each situation.

Figure 1 shows that the approximate *d*^2^ or *δ*^2^ are both slightly higher than their exact value, and both the exact and the approximate *d*^2^ or *δ*^2^ decrease as *N_e_* or *v* increases. The values of 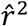 and 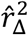 are both initially 1, and reduce respectively to exact *d*^2^ and *δ*^2^ after about 40 generations. Henceforth, 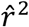 and 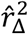 achieve a relatively stable state, and remain around the exact values of *d*^2^ and *δ*^2^ for several generations. In particular, if the ploidy level is four, these values will both converge to the exact *d*^2^ and *δ*^2^ as the number of generations increases.

Due to genetic drift, some loci become fixed and are excluded from the simulation, causing that the number *L* of locus pairs used for genotyping to decline. The correlation between the numerator and the denominator in each of both formulas in Equation (9) therefore increases, such that 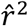 and 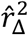 correspondingly decrease. The duration of a stable state depends on three factors: (i) ploidy level *v*, (ii) effective population size *N_e_* and (iii) the number *L* of locus pairs. As the value of each of these factors increases, the longer the duration of the stable state of both 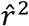 and 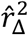.

We also simulate the behaviors of 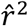 and 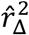 during reproduction for five mating systems (including cases with *f* being set to either 2 or 5 for the DR and the DH mating systems). The simulation process is as follows. First, a population for each of the five mating systems is generated, which contains 40 individuals with a ploidy level of either 2, 4, 6 or 8. Next, these 40 individuals are genotyped as described for the previous simulation. Then, the population is allowed to reproduce for 50 generations. For each generation, by using data of the genotypes of all individuals under various situations, 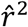 and 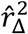 are calculated. The exact and approximate *d*^2^ and *δ*^2^ values are also calculated. The process is repeated 30,000 times. The results are shown in Figure S1, and are similar to those shown in Figure 1. However, the approximate values *d*^2^ and *δ*^2^ deviate more from their exact values for some mating systems.

Finally, we also simulate the behaviors of 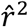 and 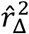 for the MS mating system under different recombination rates (set *N_e_* = 80, *v* = 2 or 4, *L* = 200 and *c* = 0.001, 0.002, 0.004, 0.01, 0.02, 0.04, 1 or 2). The simulation process is similar to the previous method and is performed 20,000 times. The population is allowed to reproduce for 100 generations. For each generation, 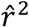 and 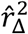 are calculated, with the results shown in Figure S2. This shows that the convergent rates for 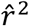 or 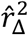 among different ploidy levels differ little as the number of generations increases, but are strongly affected by the recombination rate: the higher the recombination rate, the faster the rate of convergence.

#### Recombination rate

To investigate the influence of the recombination rate *c* on *d*^2^ and *δ*^2^, the exact and the approximate *d*^2^ and *δ*^2^ are calculated for each mating system under different recombination rates (set *N_e_* = 100, *n* = 100, *v* = 2, 4, 6 or 8, *f* = 1 for DR and *f* = 2 or 5 for DH). The recombination rate *c* ranges from 0 to 1. The results for the MS mating system are shown in Figure 2, and the results for all mating systems (including MS) are uniformly shown in Figure S3.

**Figure 2.**
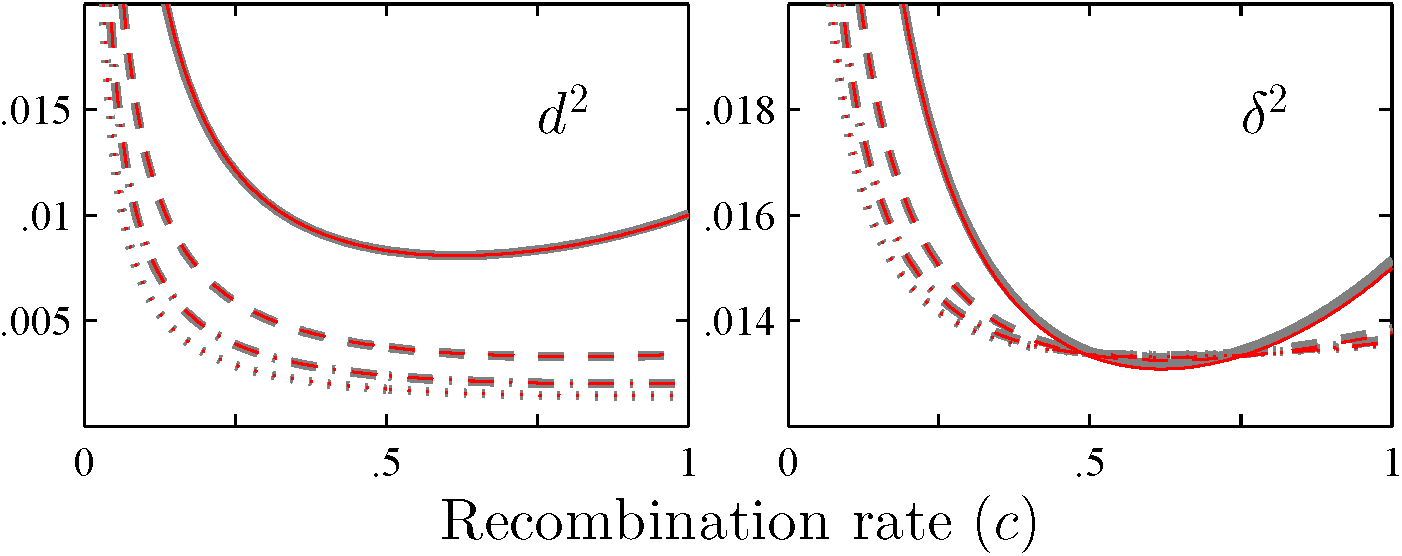
The relationship between *d*^2^ (or *δ*^2^) and the recombination rate *c* for the MS mating system (set *N_e_* = 100, *n* = 100 and *v* = 2, 4, 6 or 8). The solid, dashed, dash-dotted and dotted lines denote the values for diploids, tetraploids, hexaploids and octoploids in turn, and the black and the red lines denote the exact and the approximate values, respectively.

Figure 2 shows that *d*^2^ or *δ*^2^ are high at a low recombination rate, and decrease gradually to a relatively low value as *c* increases. The rate of decrease steepens as the ploidy level increases. However, after *c* reaches ~0.5, *d*^2^ (at *v* = 2) or *δ*^2^ (at all ploidy levels) both begin to increase. The approximate values of *d*^2^ are close to their exact values, whilst the difference between the approximate and the exact values of *δ*^2^ are more obvious, especially when *c* > 0.5.

The exact values for *d*^2^ and *δ*^2^ for the unlinked loci located on the same or different chromosomes are calculated for all five mating systems (set *N_e_* = 100, *n* = 100, *v* = 2, 4, 6, 8 or 10, *c* = 0.5 or 1 — *1/v* and *f* = 1, 2 or 5 for DR /DH). Moreover, the error rates for *d*^2^ or *δ*^2^ under different conditions are also calculated. The results are presented in Table S6. It is clear that the difference between 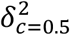 and 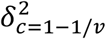 is low under all conditions, but the difference between 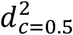 and 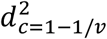 is ~50 to 100 times higher. For example, for tetraploids, the error rate is about 13% for *d*^2^ but it is 0.13% for *δ*^2^.

#### Estimation of effective population size

In this section, we estimate the effective population size *N_e_* from unphased genotypes. Previous methods for this assume that loci are all unlinked, and the recombination rate is 0.5 (e.g. England *et al.* 2006). However, in polysomic inheritances, the recombination rate is 1 — 1/*v* between two loci located on different chromosomes. Because 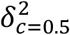 and 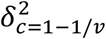 are close, with the error rate at most 1.5% (Table S6), we assume the recombination rate *c* = 0.5 between any two loci.

We preliminarily solve *N_e_* using the approximated *δ*^2^, e.g. Equation (7b), by

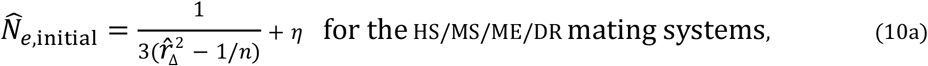

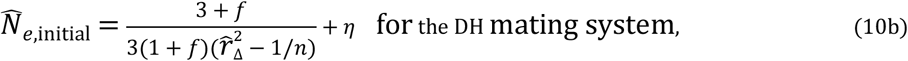

where 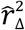 is calculated by Equation (9), and the expression of *η* is either 0 for the HS mating system, 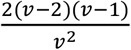 for the MS mating system, or 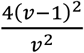 for the ME/DR/DH mating systems.

We further optimize the solution of *N_e_* using the exact *δ*^2^, i.e. Equation(6b). The exact *δ*^2^ is related to the double non-identity coefficients and the effective population size *N_e_*. Therefore, the exact *δ*^2^ can be regarded as a function of *N_e_*, denoted by *δ*^2^(*N_e_*) such that *N_e_* is the root of the equation:

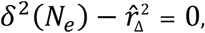

and we solve *N_e_* with Newton’s method using 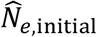 as the initial solution. This approach is denoted as NEWTON’S approach.

Equation (10a) or (10b) demonstrates that *N_e_* is approximately proportional to 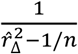. If 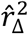 is close to 1/*n*, then 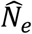 becomes very large, indicating that the estimation of *N_e_* is biased. However, because 1/*N_e_* is approximately proportional to 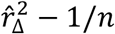, the estimation of 1/*N_e_* is almost unbiased. Therefore, it is more convenient to evaluate the statistical performance of newton’s approach by using the estimator 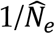.

We use a Monte-Carlo method to simulate the estimation of effective population size *N_e_* from unphased genotypes, and then evaluate the statistical performance of NEWTON’S approach under different ploidy levels, numbers of loci, numbers of alleles and sample sizes. Two types of markers are used during simulation: (i) SNP (diallelic) and (ii) SSR (hexa-allelic). For simulation, first a founder population with 200 individuals all with a ploidy level of either 2, 4, 6 or 8 is created. To avoid the fixation of alleles, each allele in the founder generation is set as being unique. Second, the 200 individuals are genotyped at 100 or 200 diallelic SNPs, or at 20 or 40 hexa-allelic SSRs. These loci are assumed to be isometrically distributed on 10 chromosomes, and the length of each chromosome is 100 cM. Third, the founder population is allowed to reproduce for a fixed number of generations to reach the linkage equilibrium; the number of generations is 44 or 86 for SNP, and 11 or 19 for SSR. Fourth, after the final generation has been attained, to reduce the number of alleles *k*, we repeat collapsing two randomly selected alleles until the value of *k* is less than 2 (for SNP) or 6 (for SSR). Fifth, for the final generation, 400 individuals are created in total, and *n* individuals are randomly sampled from this generation, where *n* = 40,80, —, 400 (interval 40). Finally, using the data of unphased genotypes of the *n* individuals sampled (*n* = 40,80, —,400), 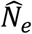 can be estimated by using newton’s approach. This process is performed 2000 times. If we subsquently let 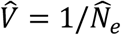, the RMSE and the bias of 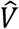 can be calculated, the results being shown in Figures 3 and S4.

**Figure 3.**
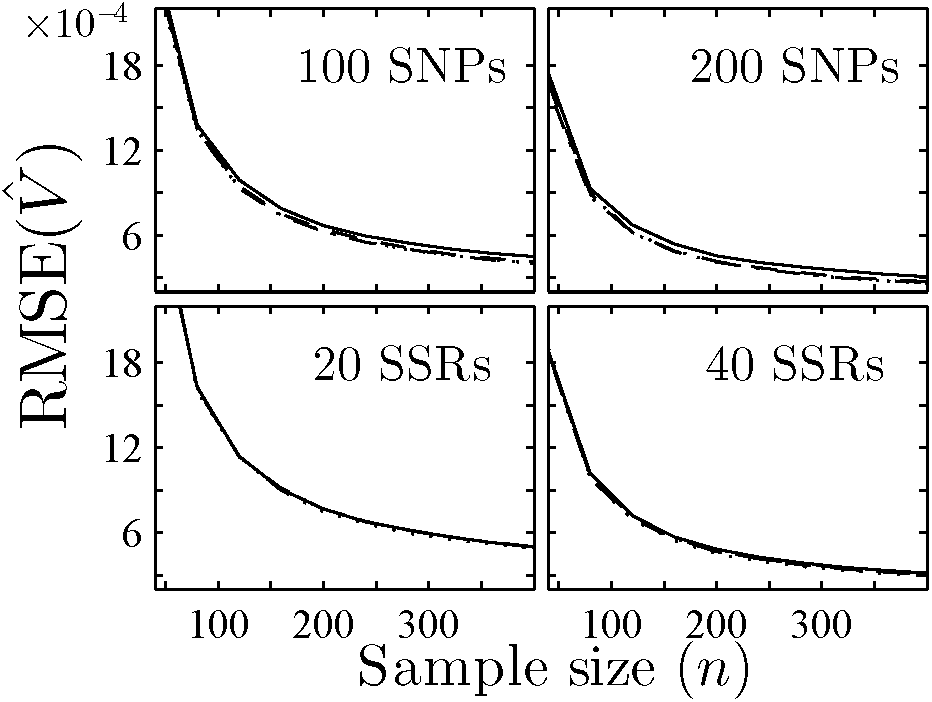
The relationship between the RMSE of 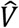 and the sample size *n* (set *N_e_* = 200, *v* = 2, 4, 6 or 8, *L* = 100 or 200 for SNP and *L* = 20 or 40 for SSR). The results are obtained from the unphased genotypes of 40 to 400 individuals (interval 40). The solid, dashed, dash-dotted and dotted lines denote results for disomic, tetrasomic, hexasomic and octosomic inheritances in turn.

Figure S4 shows that the results for SNP are more biased than those for SSR, with 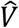 slightly increasing as the number of loci *L* also increases. The bias of 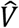 is small, and is generally less than 2 × 10^-3^, especially less than 3 × 10^-4^ for the hexasomic and the octosomic inheritances, thus 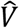 is nearly unbiased, as expected. Therefore, each RMSE of 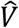 is approximately equal to its standard deviation. Figure 3 shows that the RMSEs of 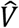 decrease as *n* increases, the values of which among different ploidy levels are similar. Moreover, the RMSEs for polyploids are slightly smaller than that for diploids. in general, the performances of SNPs and SSRs are similar.

## Data Accessibility

polygene is a user-friendly software and can be run on 64bit Windows/Linux/Mac OS X platform. The binary executable files, user manual and source code of polygene V1.3 are available on GitHub (https://github.com/huangkang1987/polygene). polygene was developed using C++ and C# and requires.net framework V4.0 runtime library.

The simulation program, the source code (developed with C#), the readme file and the script to generate figures (developed with matlab) are provided as supplementary materials and can be downloaded via a link on the online version of this manuscript.

## Discussion

### LD test

We here follow the method proposed by Weir and Cockerham (1979) to extend two LD measures, *D* and the Burrow’s *Δ*, to account for different levels of polysomic inheritance. These two measures can be used to perform the LD test. The null hypothesis of a LD test is that a pair of loci is under linkage equilibrium, which is equivalent to all *D_AB_* (or all Δ_*AB*_) values being equal to zero.

For a sample with *n* individuals, there are *nv* haplotypes. The observed and the expected occurrences of a haplotype *AB* are, respectively, 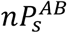 and *np_A_q_B_*. Because 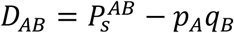, the *χ*^2^ statistic for the LD measure *D* can be established as follows:

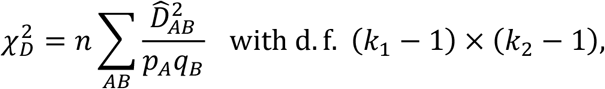

where d.f. is the number of degrees of freedom, *k_i_* is the number of alleles among the allele copies in those haplotypes at the *i*^th^ locus (*i* = 1,2), *A* is taken from all *k*_1_ alleles at the first locus, and *B* is taken from all *k*_2_ alleles at the second locus.

Next, for a sample with *n* individuals, there are *nv*^2^ allele pairs, the observed and the expected occurrences of an allele pair *AB* are respectively 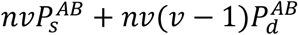 and 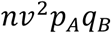. Because 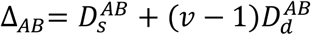 and 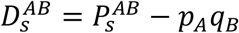, the *χ*^2^ statistic for Burrow’s *Δ* statistic can be established as follows:

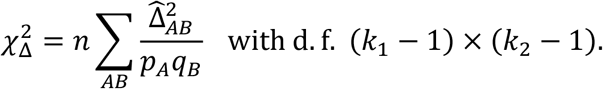

### *d*^2^ and *δ*^2^

In this study, various moments of LD measures are derived by extending Weir & Hill’s (1980) double non-identity coefficients, and thus the exact *d*^2^ can be obtained by using the moments 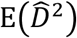 and 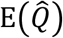 under various mating systems. The exact *δ*^2^ can also be obtained by using the moments 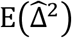 and 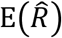. Hence the value of 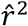 (or 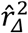) can be approximately replaced by that of *d*^2^ (or *δ*^2^) under each mating system at the equilibrium state. Moreover, the approximate expressions of *d*^2^ and *δ*^2^ under various mating systems are derived by using the transitional matrix, and several relations are discussed, such as the relation between 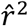 (or 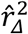) and the number of generations during reproduction, the relation between *d*^2^ (or *δ*^2^) and the recombination rate *c*, and so on.

Figure 1 shows that after the population has been allowed to reproduce for about 40 generations, 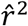 (or 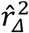) reaches a relatively steady state remains close to the exact *d*^2^ (or *δ*^2^) for several generations. After that, 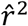 (or 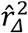) begins to decrease again, due to both the fixation of alleles and the positive correlation between 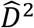 and 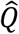 (or between 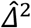 and 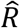). As the number *L* of diallelic locus pairs decreases, the number of terms in the numerator or the denominator in Equation (9) is reduced, due to the weighted scheme in Equation (9) being unable to effectively eliminate the correlation. The number of generations at which 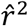 (or 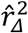) begins to decrease again depends on *v, N_e_, L* and the initial heterozygosity.

Figure S2 shows that regardless of 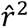 or 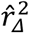, the smaller the recombination rate, the slower the rate of convergence. Generally, 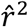 and 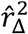 decrease to a relatively steady state after about —4.21/ln1 — *c* generations. Moreover, under the same recombination rate, the convergent rates of 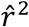 (or 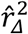) are similar for all levels of ploidy but differ markedly under different recombination rates.

Figure 2 (and S3) show that the relationship between *d*^2^ (or *δ*^2^) and the recombination rate *c* has two main features: (i) if *c* is small (e.g. < 0.25), both *d*^2^ and *δ*^2^ for polysomic inheritance decreases more rapidly than those for disomic inheritance and (ii), the difference between 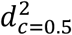 and 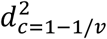 under polysomic inheritance is considerable (the error rate ranges from 10% to 23%), whereas the difference between 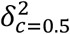 and 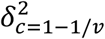 is negligible (the error rate is less than 1.5% for non-HS mating systems).

For (i), this infers that a higher density genetic map is required to detect any linkage in polyploids. A rough estimate would be the locus density in tetraploids (hexaploids or octoploids) to be 1.58 (2.16 or 2.67) times of that for diploids (estimated by the threshold *δ*^2^ = 0.2, see Figure 2). However, if the locus density is sufficient, the gene mapping in polyploids may be more accurate than that in diploids due to the steep slope of the curve at a low *c*.

For (ii) this indicates that it is unnecessary to distinguish whether two loci are located on the same chromosome or not if the effective population size *N_e_* is estimated by 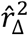. From this reason, we can simply let the recombination rate between any two loci be equal to 0.5, as is assumed in other methods (e.g. England *et al.* 2006). However, it is necessary to model the case that two loci are located on different chromosomes if *N_e_* is estimated by 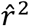 using phased genotypes.

### Effective population size

Among the parameters 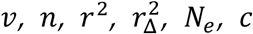 and *f,* the first two *v* and *n* are known, the next two *r*^2^ and 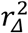 can be estimated from the genotype data, and the mating system and the ratio *f* can be obtained from either *a priori* information, field observations or experiments. The remaining two parameters *N_e_* and *c* are the parameters we usually need to estimate, and one can be estimated if the other is known.

After simulation, we evaluate the RMSE and the bias of 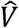 (i.e. 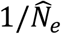). However, in practice, we are interested in *N_e_*. To ensure that the estimation of *N_e_* is applicable, the estimator 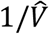 (i.e. 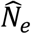) should be within ± 20% of the true *N_e_*. Therefore, the RMSE of 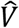 should be smaller than 1/10N_e_. For example, if *N_e_* = 1000, the RMSE of 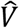 should be below 0.0001. It is hard to achieve such a small value if *N_e_* is large. This is because the influence of sampling error 1 /*n* on *δ*^2^ is relatively high, which covers the effect of *N_e_* on *δ*^2^ (e.g. see Equation (6b)), and thus more samples are required to reduce the sampling variance.

From Figure 3 it is clear that the curves of RMSE among different ploidy levels are similar, indicating that the estimations of the effective population sizes *N_e_* in polyploids is almost the same as for diploids. Such estimation therefore requires similar numbers of samples and loci regardless of ploidy level. Figure S4 also shows that the results for polyploids may be better than for diploids due to smaller biases. The performance of 100/200 diallelic SNPs is as good as that of 20/40 hexa-allelic SSRs, indicating that the RMSE is mainly determined by the number 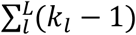 of independent alleles.

From Figure S4, the estimation of 1/*N_e_* is biased. Some possible sources of this bias are enumerated as follows. (i) According to Equation (10a) or (10b), 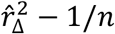 is actually proportional to 1/(*N_e_* — *η*), not 1/*N_e_*, indicating that the estimation of 1/(*N_e_* — *η*) may be unbiased, but the estimation of *1/N_e_* is biased. (ii) The recombination rate between two loci located on the same chromosome is less than 0.5, but it is assumed to be 0.5. (iii) The recombination rate between two loci located on different chromosomes is 1 — 1/*v*, but it is also assumed to be 0.5.

We suggest that (ii) is the main source of this bias. This is because the bias is largely influenced by both the number *L* of loci used and the ploidy level *v* (Figure S4). Because the length of each chromosome is a constant in each simulation, the loci become denser at higher levels of *L*. The value of *δ*^2^ between two close loci (implying smaller *c*) therefore increases in the deviation from 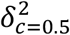 (Figure 2). In addition, the simulation results for polyploids are less biased. This is because the curve of *δ*^2^ at a higher ploidy level is flat for the majority of situations (e.g. *c* > 0.2). To validate our prediction, we simulated the scenario that the loci on the same chromosome are all unlinked (i.e. *c* = 0.5 for all locus pairs) and found that bias is reduced to ~10_^4^ in diploids and in tetraploids (data not shown).

The bias sources (ii) and (iii) can be reduced if the *a priori* information about the length of chromosomes is considered. For this case, it is possible to calculate *δ*^2^ by

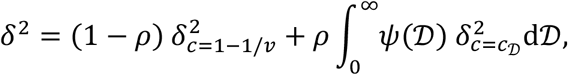

where *ρ* is the probability that two loci are on the same chromosome, 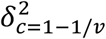 and 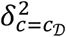 are the exact *δ*^2^ at the recombination rates 1 — 1/*v* and 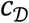, respectively, 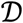 is the genetic distance between two loci in the same chromosome, and 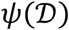 is the probability density function of 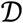, which is the following weighted average:

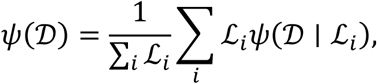

in which 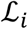 is the length of *i*^th^ chromosome, and 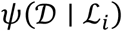 is the probability density function of 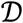 given 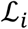, whose expression is

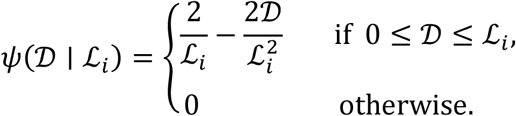

## Acknowledgment

KH would like to thank Prof. Kermit Ritland for providing a visiting professor position at UBC.

## Funding

This study is funded by the Strategic Priority Research Program of the Chinese Academy of Sciences (XDB31020302), the National Natural Science Foundation of China (31770411 and 31730104), and the National Key Programme of Research and Development, Ministry of Science and Technology (2016YFC0503200). DWD is supported by a Shaanxi Province Talents 100 Fellowship and KH is supported by a scholarship from China Scholarship Council.

## Conflicts of Interest

The authors declare no conflict of interest.

## Author Contributions

KH and BGL designed the project, KH constructed the model and wrote the draft, WKL and DW performed simulations and drawn the figures, and DWD checked the model and helped to write the manuscript.

## Supplementary Materials

### Appendix A. Expansion of Δ_*AB*_ in triploids

In triploids, the value of Δ_*AB*_ is given by 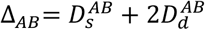, where 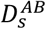 and 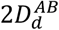 can be respectively expanded as

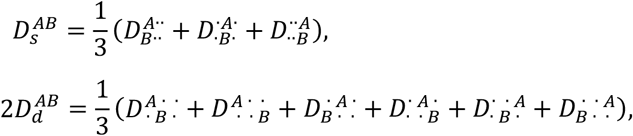

in which the superscripts (or the subscripts) of *D* in the right side of equal sign denote the phased genotype at the first (or the second) locus, and the dot · denotes any allele.

Each term in the right side can be further expanded as follows (the terms with the same two-locus unphased genotypes in the expansion are put together):

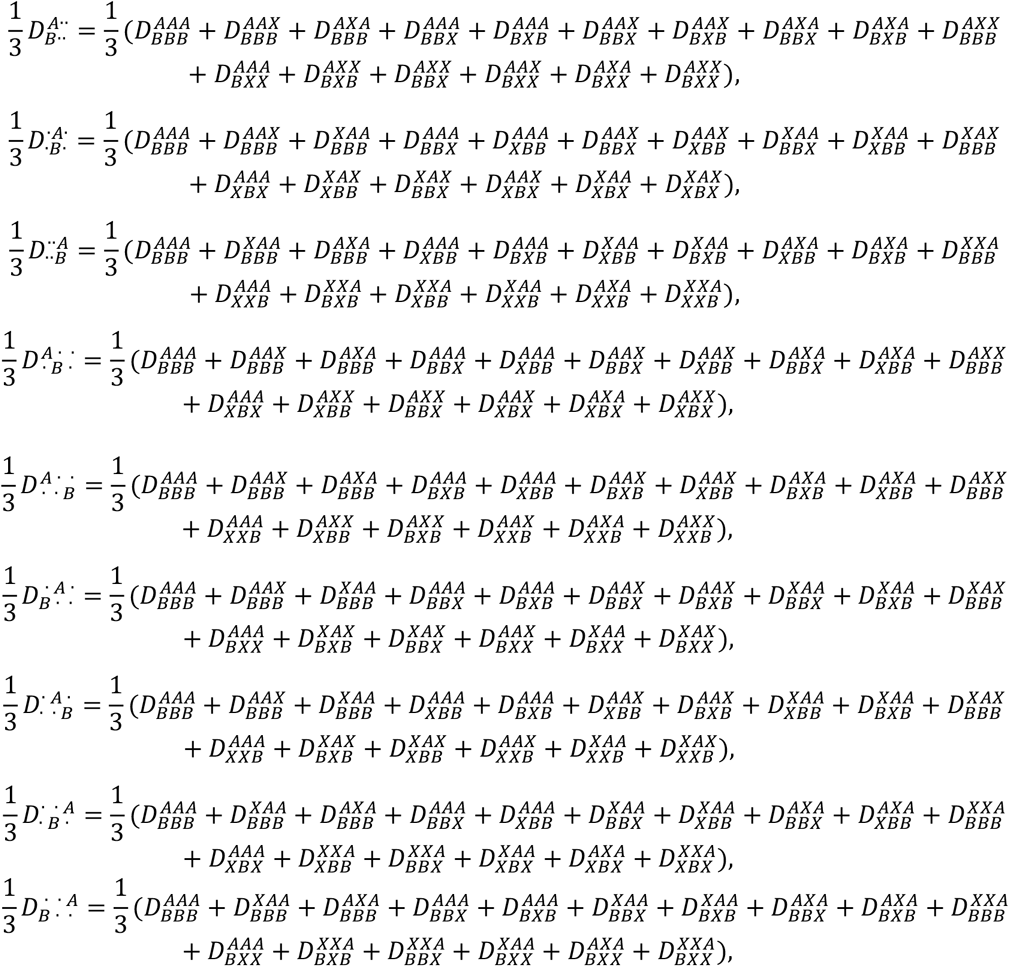

By summing all terms of the same two-locus unphased genotypes in the right sides of equal signs, we obtain the following equalities:

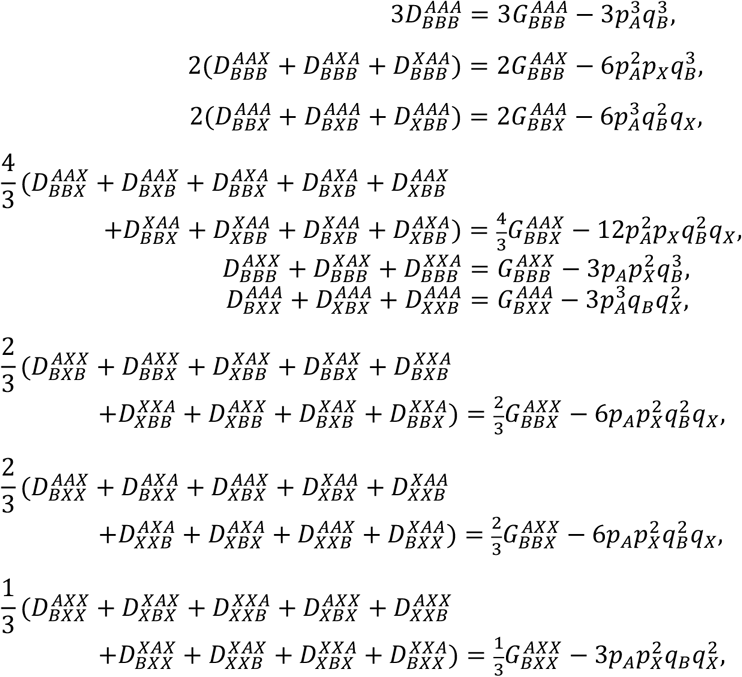

where each 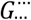 denotes a two-locus unphased genotypic frequency, whose superscript and subscript represent two unphased genotypes. It can be seen that each expression on the right sides of above equalities is one of the following:

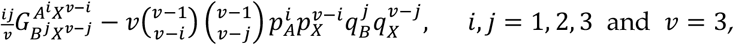

in which *A^i^X^v-i^* denotes an unphased genotype containing exactly *i* copies of *A*, and the meaning of *B^j^X^v-j^* is similar. Because Δ_*AB*_ is the sum of these expressions, it follows

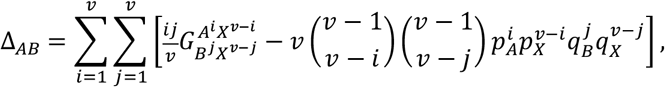

where *v* = 3. This formula can be extended to a higher ploidy level *v* by similar discussion. Note that

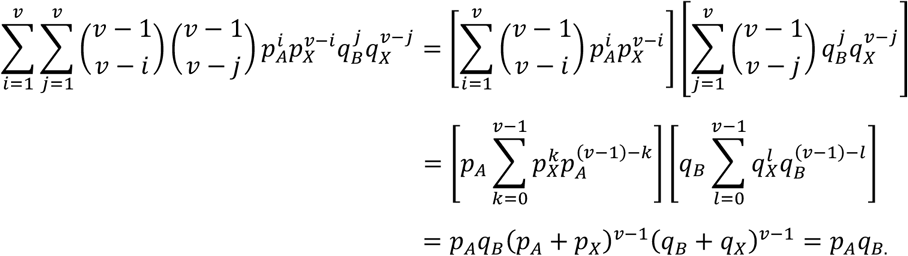

The next formula is valid for any ploidy level *v*:

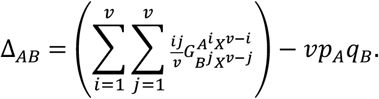

### Appendix B. Derivation of moments of LD measurements

In the process of deriving the moments of LD measurements, we need to use the estimates of frequencies *P*_1_, *P*_2_, –, *P*_22_ listed in Table 1, and so we first discuss 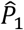 to 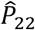, and then derive various moments.

#### 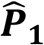 to 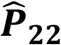

Denote *x_ij_* for the state indicator of allele *A* related to the *j*^th^ haplotype within the *i*^th^ individual at the first locus, and *y_ij_* for that of *B* at the second locus, where *x_ij_* = 1 if the allele copy at the first locus is *A*, otherwise *x_ij_* = 0; the meaning of *y_ij_* is similar. Moreover, we let the number of the sampled individuals be *n* (the sample size), and let the number of haplotypes within each individual be *v* (the ploidy level). Then 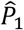 to 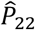 can be expressed as follows.

**Digenic:**

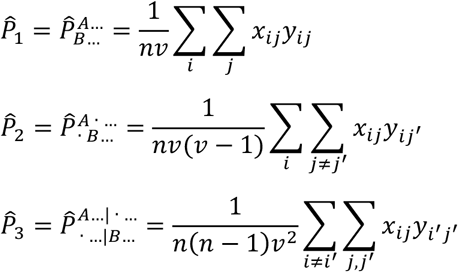

**Trigenic:**

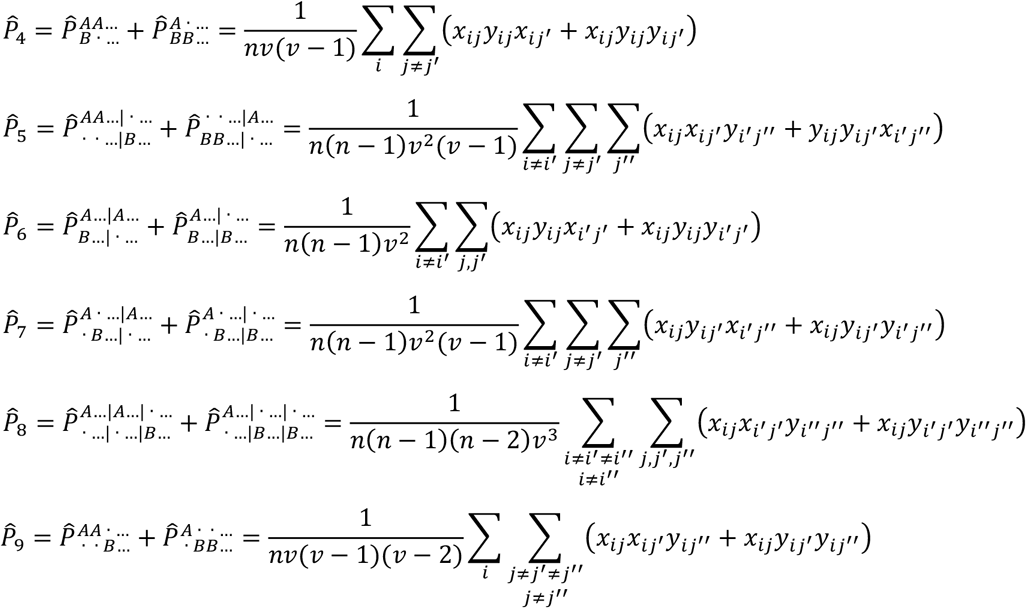

**Quadrigenic:**

Dihaplotypic:

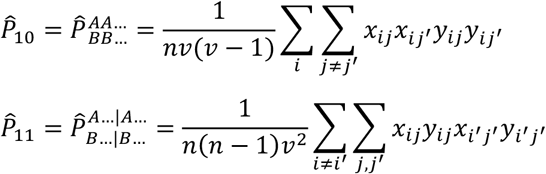

Trihaplotypic:

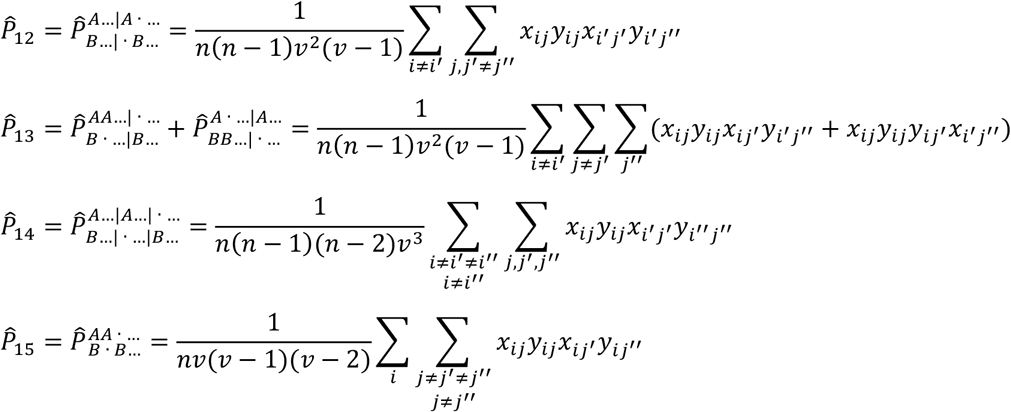

Dihaplotypic:

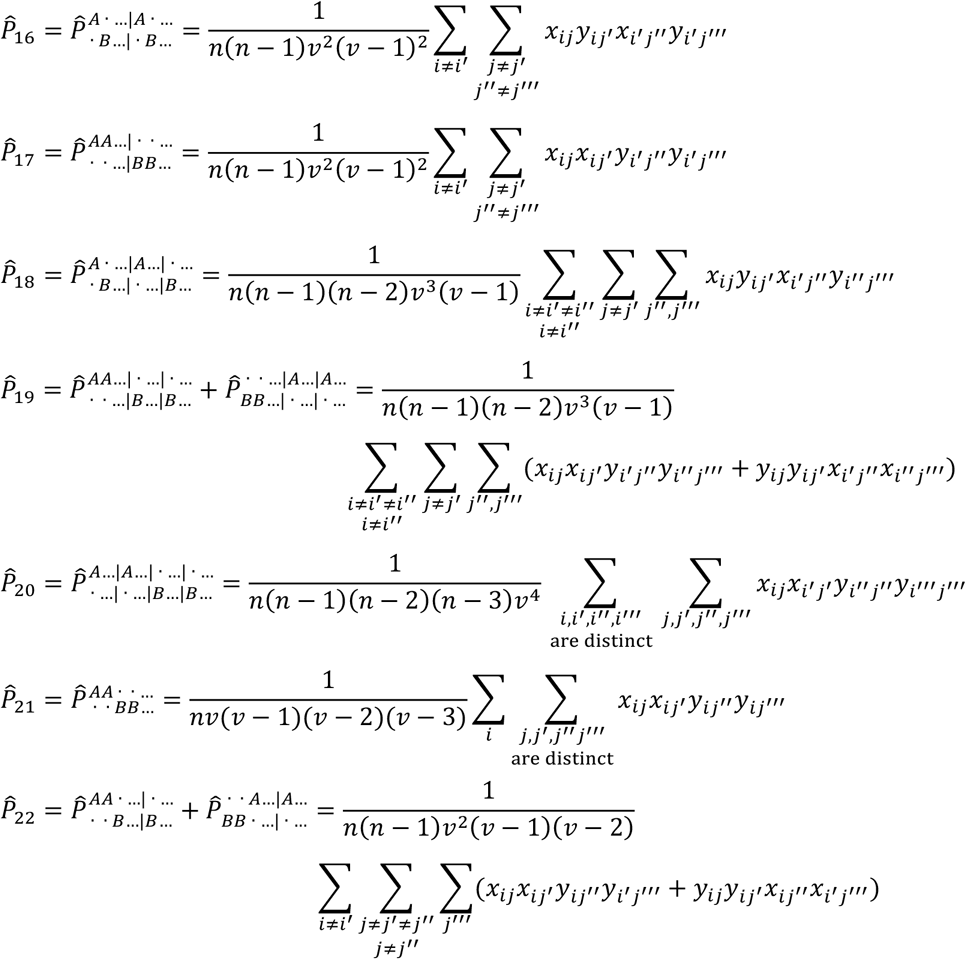

#### 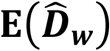 and 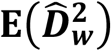

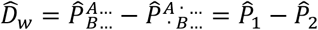 by the definition of *D_w_*, in which

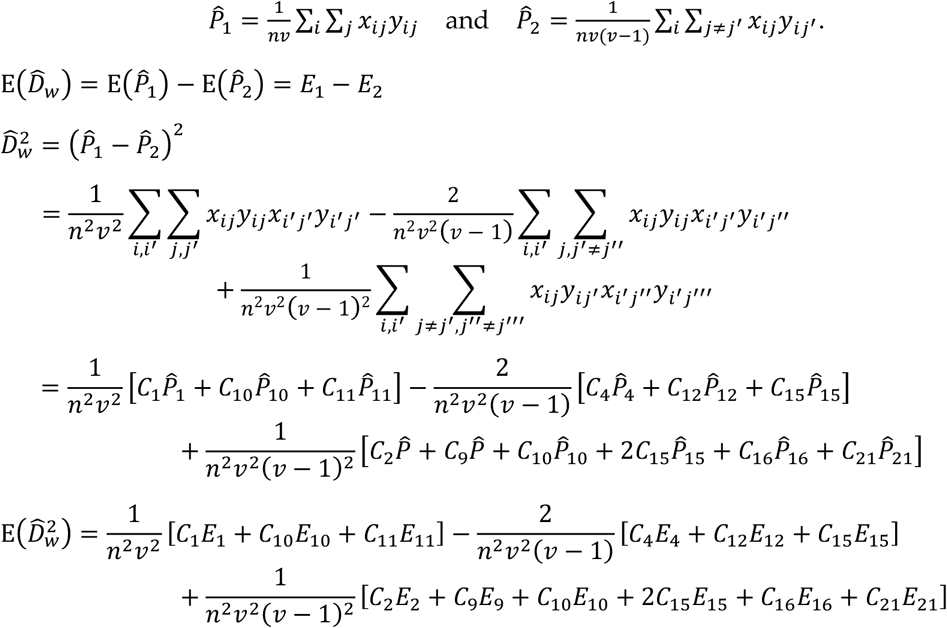

where the coefficient *c_i_* is the reciprocal of coefficient before the summation sign in the expression of 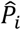, e.g. the final coefficient *C*_21_ is *nv*(*y* — 1)(*v* — 2)(*v* – 3).

#### 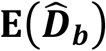 and 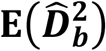

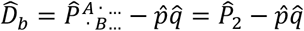 by the definition of *D_b_* in which

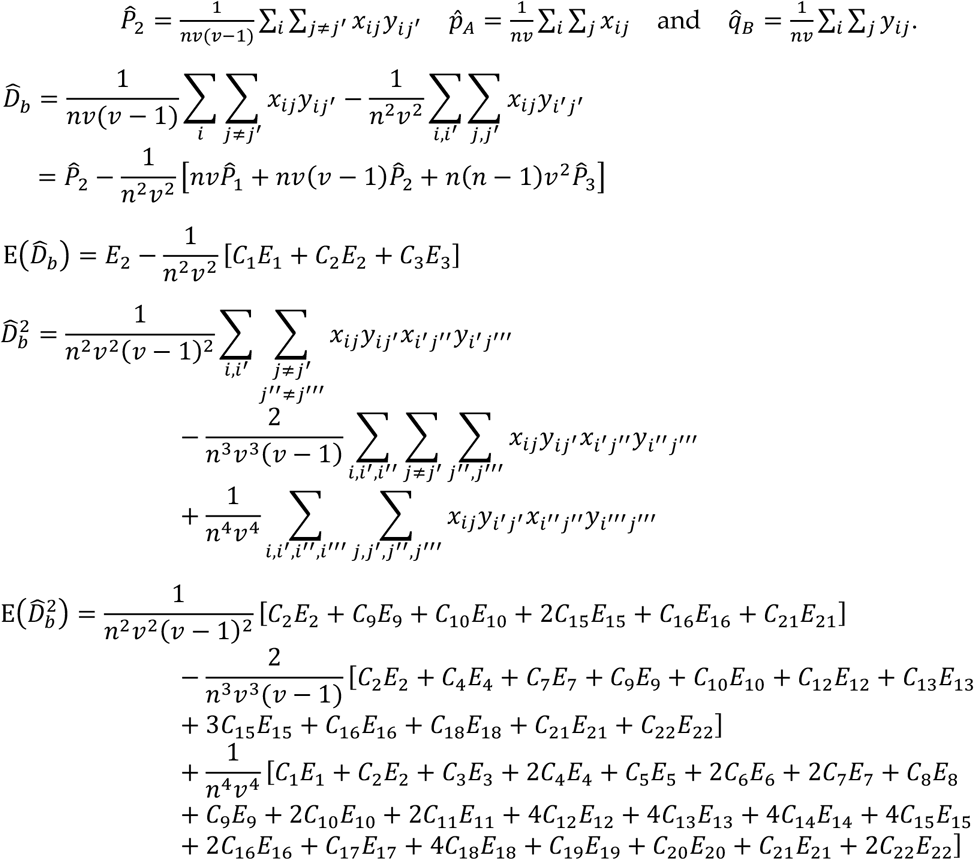

#### 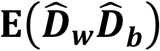

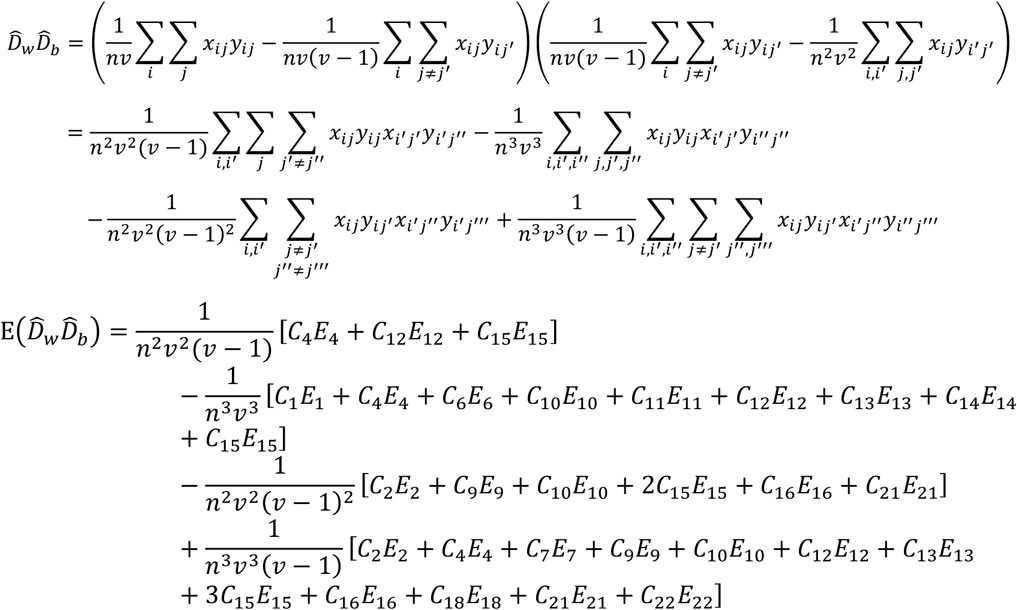

#### 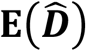 and 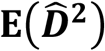

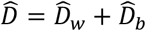 by the definition of *D*, then

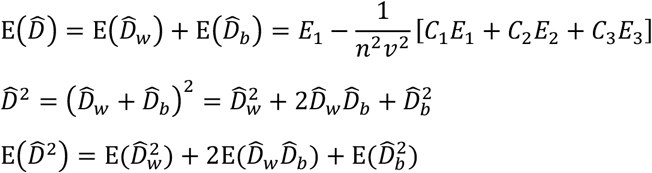

#### 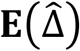 and 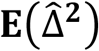

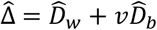 by the definition of *Δ*, then

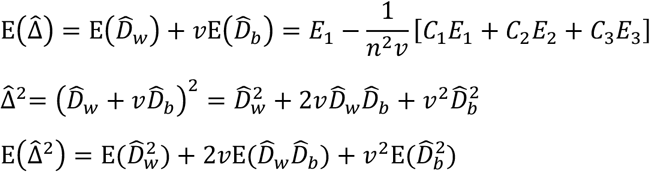

#### 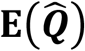

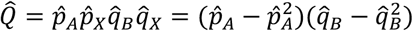 by Q = *p_A_p_X_q_B_q_X_* in which

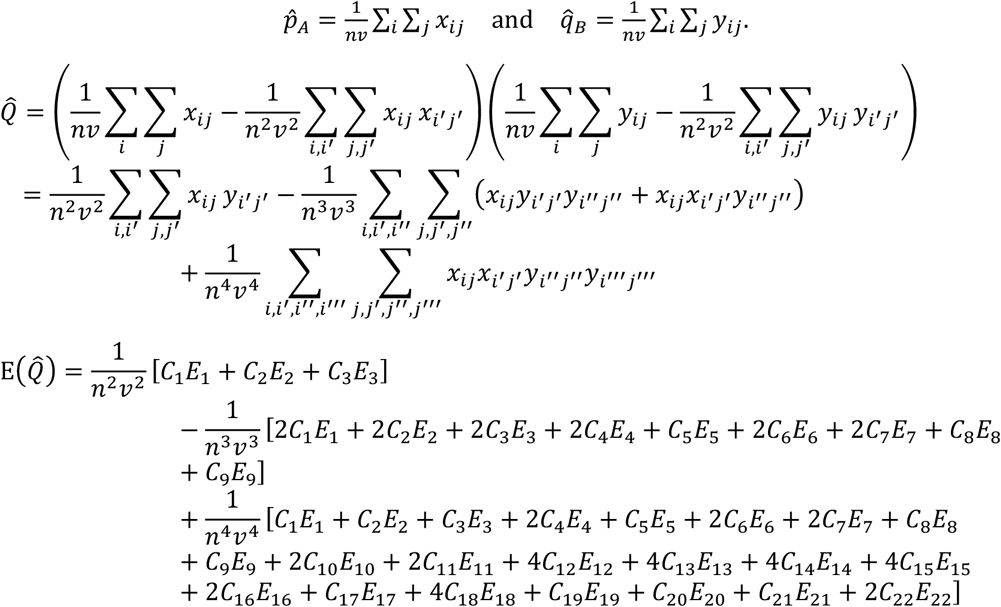

#### 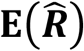

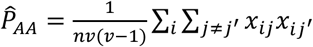 and 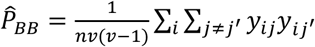 by the definition of *P_AA_*. 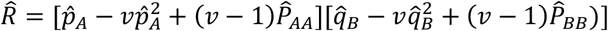 by Equation (2).

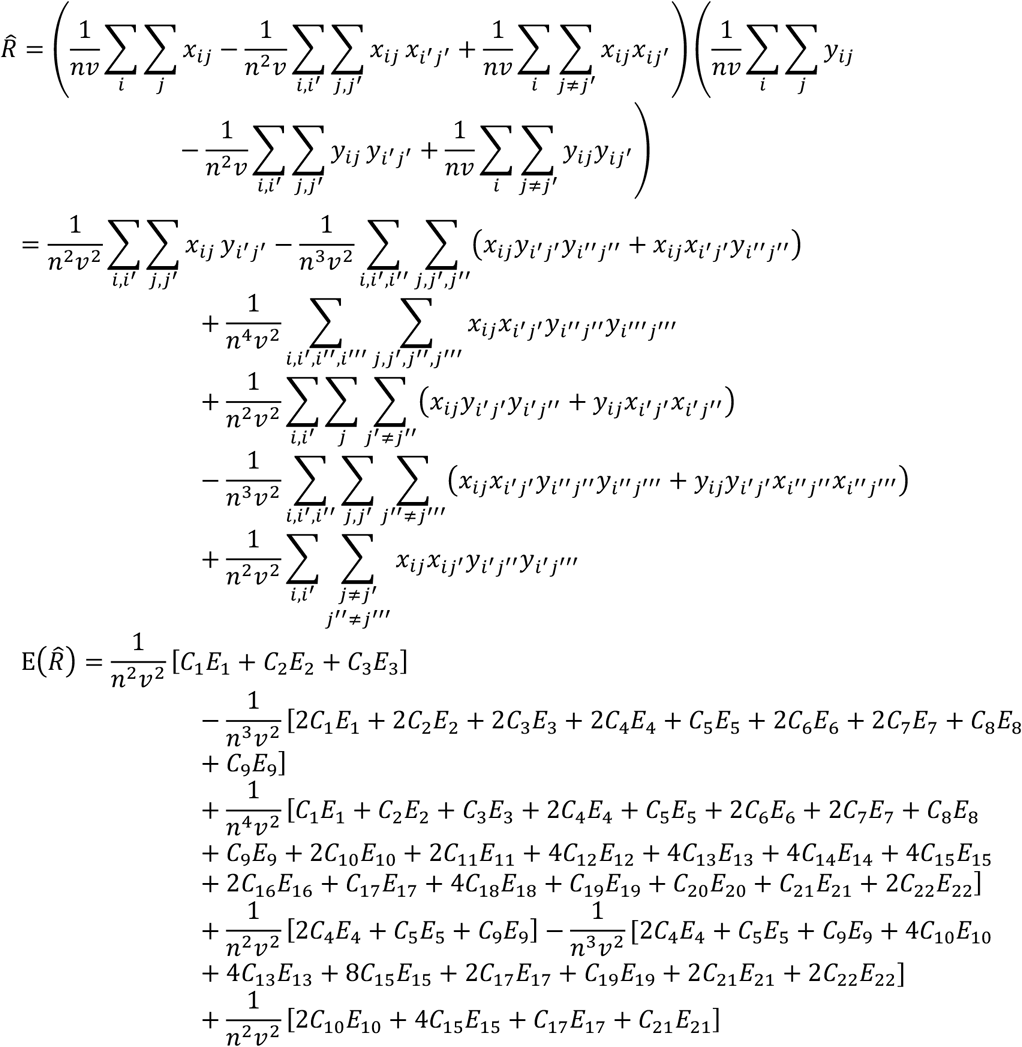

### Appendix C. HS mating system

The double non-identity coefficients are closely related to the haplotypes that are used to detect the specific alleles. In this appendix, we consider such relationships in the haplotype sampling (HS) mating system. The effective population size *N_e_* in this system is assumed to be the same as the population size *N*. We will adopt *N_e_* instead of *N* in our discussion in order to accommodate other mating systems, and we will also write the double non-identity coefficient as the dni-coefficient for brevity.

For the case of dni-coefficients Θ_1_ and Θ_2_, it can be seen from Table 1 that only two haplotypes (say *H*_1_ and *H*_2_) are sampled, and both alleles in each haplotype need to be detected. These two haplotypes can be copied from either the same haplotype (written as *H*_1_ ≡ *H*_2_), or different haplotypes in the same individual (written as *H*_1_ = *H*_2_), or different haplotypes in different individuals (written as *H*_1_ ~ *H*_2_). For this case, we will divide into three situations (named HSΘ1, HSΘ2 and HSΘ3) to carry out our discussion.

HSΘ1 *H*_1_ ≡ *H*_2_, weight 1, dni-coefficient 0;

HSΘ2 *H*_1_ ≍ *H*_2_, weight *v* — 1,

a. none recombined, probability (1 – *c*)^2^, dni-coefficient Θ_1_;
b. one recombined, probability 2*c*(1 — *c*), dni-coefficient 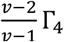;
c. both recombined, prob. *c*^2^, dni-coefficient 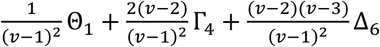;

HSΘ3 *H*_1_ ~ *H*_2_, weight (*N_e_* – 1)*v*,

a. none recombined, probability (1 – *c*)^2^, dni-coefficient Θ_2_;
b. one recombined, probability 2*c*(1 — *c*), dni-coefficient Γ_1_;
c. both recombined, probability *c*^2^, dni-coefficient Δ_1_.

Now, the expression of dni-coefficients 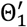 or 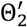 in the next generation can be written out. In fact, let **W**_*θ*_ = [*w*_1*θ*_, *w*_2*θ*_, *w*_3*θ*_] be the vector consisting of those weights, i.e. **W**_*θ*_ = [1, *v* — 1, (*N_e_* — 1)*v*], and let **Θ** = [*θ*_1_, *θ*_2_, *θ*_3_], where each *θ_i_* is the weighted sum of dni-coefficients in HSΘ*i*, with the corresponding recombination probabilities as their weights (if the recombination probability does not occur, *θ_i_* is set as the dni-coefficient in HSΘ*i*), *i* = 1,2,3, that is

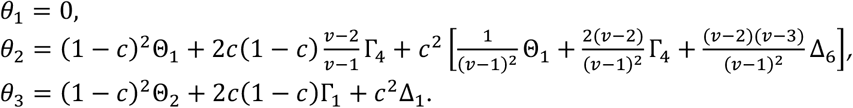

Then 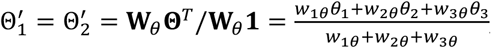, where **1** is the column vector [1,1,1]^*T*^. This is a linear combination of dni-coefficients in the current generation, and the products of combination coefficients times *N_e_v*(*v* – 1) are listed in the second column of Table S3.

For the case of Γ_1_ to Γ_4_, it can be seen from Table 1 that three haplotypes are sampled, in which one is the haplotype that both alleles need to be detected, denoted by *H*_1_, another is that only the allele at the first locus needs to be detected, denoted by *H*_2_, and the third is that only the allele at the second locus needs to be detected, denoted by *H*_3_. Because *H*_2_ and *H*_3_ are only detected the allele at a single locus, it is unnecessary to model their recombination. For this case, the combinations among three relations ≡, ≍ and ~ can be divided into nine situations (named HSΓ1 to HSΓ9).

HSΓ1 *H*_1_ ≍ *H*_2_ ≡ *H*_3_, weight 1, dni-coefficient 0;

HSΓ2 *H*_1_ ≡ *H*_2_ ≍ *H*_3_ or *H*_1_ ≡ *H*_3_ ≍ *H*_2_, weight 2(*v* — 1),

a. not recombined, probability 1 — *c*, dni-coefficient 0;
b. recombined, probability *c*, dni-coefficient 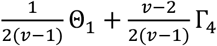;

HSΓ3 *H*_1_≍*H*_2_ ≡ *H*_3_, weight *v* – 1,

a. not recombined, probability 1 — *c*, dni-coefficient Θ_1_;
b. recombined, probability *c*, dni-coefficient 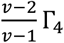;

HSΓ4 *H*_1_≍*H*_2_ ≡ *H*_3_, weight (*v* – 1)(*v* – 2),

a. not recombined, probability 1 — *c*, dni-coefficient Γ_4_;
b. recombined, probability *c*, dni-coefficient 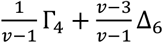;

HSΓ5 *H*_1_≡*H*_2_ ~ *H*_3_ or *H*_1_≡*H*_3_ ~ *H*_2_, weight 2(*N_e_* – 1)*v*,

a. not recombined, probability 1 — *c*, dni-coefficient 0;
b. recombined, probability c, dni-coefficient Γ_2_/2;

HSΓ6 *H*_2_≡*H*_3_ ~*H*_1_, weight (*N_e_* – 1)*v*,

a. not recombined, probability 1 – *c*, dni-coefficient Θ_2_;
b. recombined, probability *c*, dni-coefficient Γ_1_;

HSΓ7 *H*_1_≍*H*_2_ ~ *H*_3_ or *H*_1_≍*H*_3_ ~ *H*_2_, weight 2(*N_e_* – 1)*v*(*v* – 1),

a. not recombined, probability 1 – *c*, dni-coefficient Γ_2_;
b. recombined, probability *c*, dni-coefficient 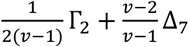;

HSΓ8 *H*_1_~*H*_2_≍ *H*_3_, weight (*N_e_* — 1)*v*(*v* — 1),

a. not recombined, probability 1 — *c*, dni-coefficient Γ_1_;
b. recombined, probability c, dni-coefficient Δ_1_;

HSΓ9 *H*_1_~*H*_2_~ *H*_3_, weight (*N_e_* — 1)(*NN_e_* — 2)*v*^2^,

a. not recombined, probability 1 — *c*, dni-coefficient Γ_3_;
b. recombined, probability *c*, dni-coefficient Δ_3_.

Now, let **W**_*γ*_ = [*w*_1*γ*_, *w*_2*γ*_, —, *w*_9*γ*_] and Γ = [*γ*_1_, *γ*_2_, —, *γ*_9_], where the definitions of **W**_*γ*_ and Γ are similar to those of **W**_*θ*_ and **Θ**. Then

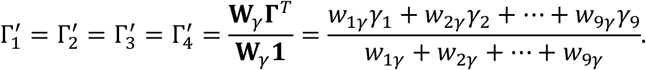

This is also a linear combination, and the products of combination coefficients times 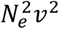 are listed in the third column of Table S3.

For the case of Δ_1_ to Δ_7_, we see from Table 1 that four haplotypes are sampled, in which two are the haplotypes that the allele at the first locus is detected, denoted by *H*_1_ and *H*_2_, and the other two are that the allele at the second locus is detected, denoted by *H*_3_ and *H*_4_. Because there is only the allele at a single locus to be detected, the recombination of these four haplotypes need not be modelled. For this case, the combinations of three relations can be divided into 22 situations (named HSΔ1 to HSΔ22).

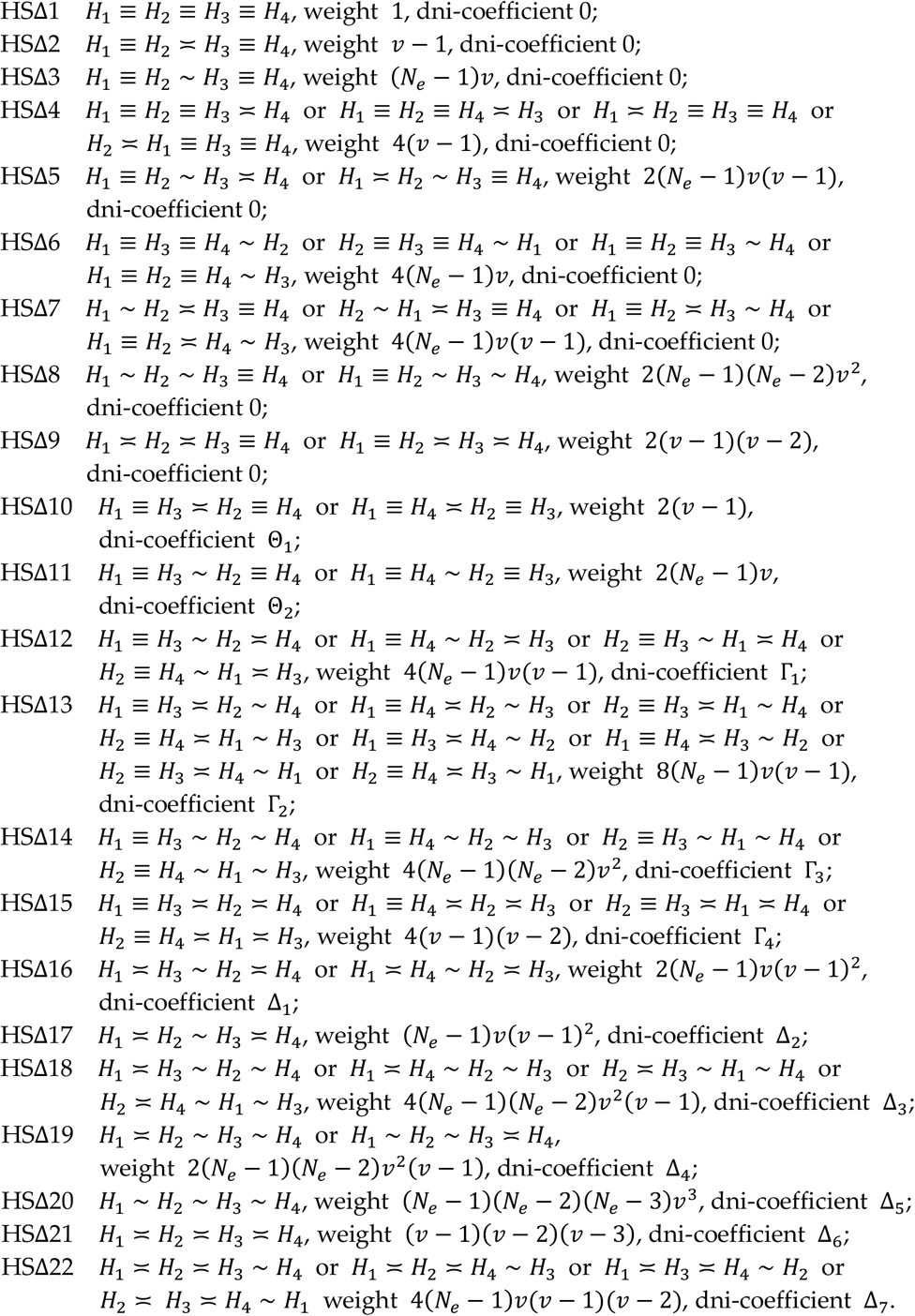

Now, let **W**_*δ*_ and **Δ** be the row vectors consisting of 22 weights and 22 dnicoefficients in HSΔ1 to HSΔ22, respectively. Then

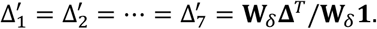

This is still a linear combination, and the products of combination coefficients times 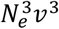 are listed in the final column of Table S3.

The expressions in Table S3 are the essential factors to form **Ω**^*T*^ of the transition matrix **Ω** for the HS mating system. Moreover, the matrices **T** and **S** in the principal part of **Ω** are listed in Appendix H.

### Appendix D. Corrections for finite sample size

It is easy to see from Equations (3) and (4) that the next formula is valid:

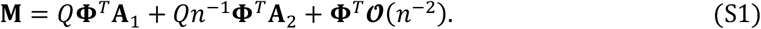

When the sample size *n* is large enough, the principal part of **M** is *Q***Φ**^*T*^**A**_1_, and the remainder can be neglected, then **M** ≈ *Q***Φ**^*T*^**A**_1_, indicating that the matrix **A**_1_ can be used to approximate the moments of LD measurements. For example, using the fourth and the sixth column elements in **A**_1_ (see Table S1), we have 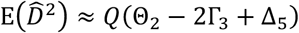 and 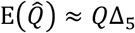. Thus

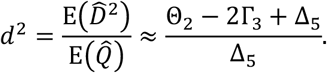

Similarly, *δ*^2^ can be approximately expressed as

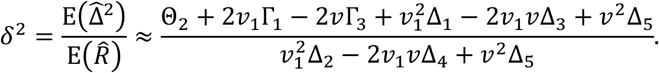

In practice, sampling error will influence the estimation of 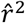 and 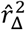. For example, if the two loci are unlinked, the moments 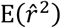 and 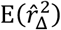 are greater than zero. This effect is related to the sample size *n*, then the term *Qn*^-1^**Φ**^*T*^**A**_2_ in Equation (S1) containing the coefficient *n*^-1^ cannot be neglected. To accommodate this effect, we consider the next approximate expression:

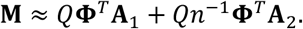

Denote the fourth expression in the vector *Qn*^-1^**Φ**^*T*^**A**_2_ by M_D/*n*_, and the sixth by M_Q/*n*_. Using the fourth and the sixth column elements in **A**_2_ (see Table S2), we obtain

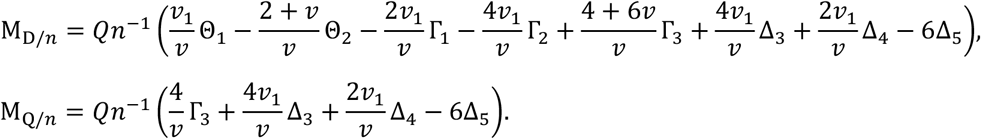

Then 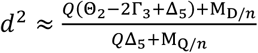. Because *n* is large, we omit M_Q/*n*_ in the denominator for the convenience of derivation. Therefore

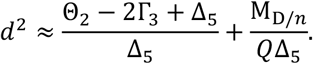

The net effect for *d*^2^ caused by 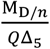 is 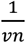 if **Φ** is an eigenvector belonging to the maximal eigenvalue of **Ω**. Here, we take the HS mating system as an example to show this. In fact, for this system, according to Equation (8) and Table 4, we have

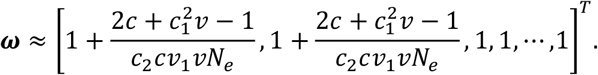

Now, replacing 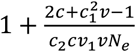 for each of Θ_1_ and Θ_2_ in 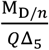 and 1 for each of the other double non-identity coefficients, it follows 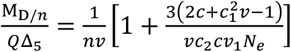. Hence 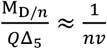 by *N_e_* ≥ *n* ≫ 1. Similarly, it can be shown that the net effect for *δ*^2^ is 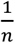. Thus, the approximate expressions of *d*^2^ and *δ*^2^ considered the effect of sample size *n* on sampling are

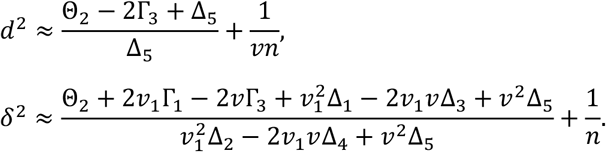

### Appendix E. Monoecious mating systems

The method to derive the expression of each element in **Ω** for the monoecious mating systems is the same as that for the HS mating system. It is noteworthy that unlike the HS mating system, two haplotypes sampled within the same individual need to be detected whether they are from the same gamete: (i) if they are from the same gamete, the probability is 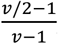,(ii) otherwise, the probability is 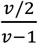. We will denote (*H, H*’, …) for which those haplotypes within brackets are from the same gamete.

For (i), it is assumed that the chromosomes form bivalents during meiosis, and the double-reduction will never happen, then the paired chromosomes will segregate into different oocytes, which means that two haplotypes *H* and *H*’ within the same gamete are copied from different haplotypes. However, this is not strictly equivalent to *H ≍ H*’. This is because the paired chromosomes will segregate into different oocytes. In order to avoid repeatition, we first discuss nine situations similar to HSΓ1 to HSΓ9 in Appendix C, named MO*r*1 to MO*r*9 in turn.

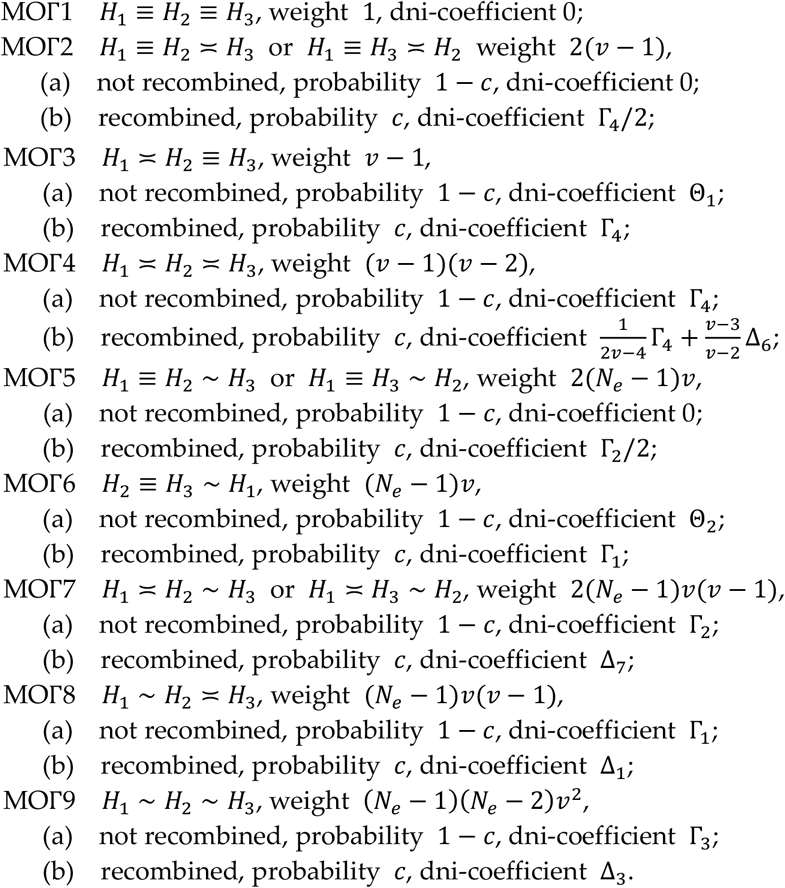

For (ii), if the mating system is MS, because selfing is allowed, two haplotypes *H* and *H*’ from different gametes can be copied from either the same haplotype (*H* ≡ *H*’), or different haplotypes in the same individual (*H ≍ H*’), or different haplotypes in different individuals (*H ~ H*’). These relationships are obviously equivalent to those in the HS mating system. If the mating system is ME, because selfing is excluded, two haplotypes from different gametes must be from different individuals (*H ~ H*’).

There may be more than one situation of HSΘ (HSΓ, HSΔ or MOΓ) appearing in an item, and we will use some symbols to denote this phenomenon. For example, the symbol (MOΓ2/2,3,4,7/2) in the item (1) of 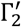 below denotes that there are four situations (i.e. MOΓ2/2, MOΓ3, MOΓ4 and MOΓ7/2) appearing in this item, in which MOΓ2/2 represents that the number of expressions describing the relations among the haplotypes is half of that for MOΓ2. This is because *H*_1_ and *H*_2_ are in the same gamete in the item (1) of 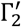, thus only the second expression in the two expressions in MOΓ2 (*H*_1_ ≡ *H*_2_ ≍ *H*_3_ and *H*_1_ ≡ *H*_3_ ≍ *H*_2_) can hold. The weight for MOΓ2/2 is also half of the weight for MOΓ2, and the meaning of MOΓ7/2 is analogous. It is noteworthy that MOΓ2 and MOΓ2/2 are the same except their number of expressions and weights, as are MOΓ7 and MOΓ7/2.

We next discuss the dni-coefficients in the next generation one by one.

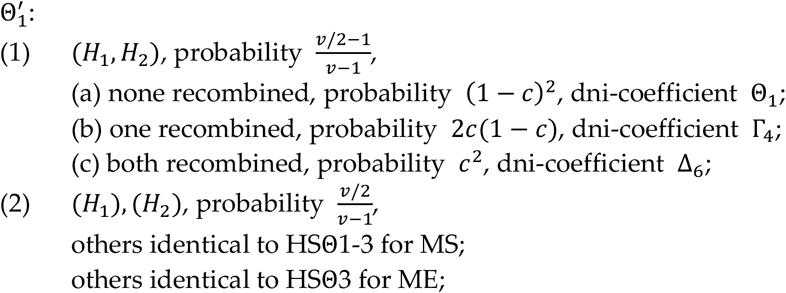

then 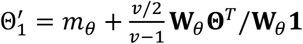 for MS, or 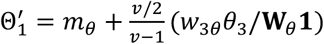 for ME, where

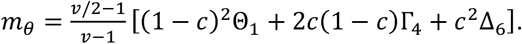

The expression of 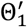 is a linear combination of dni-coefficients in the current generation, whose combination coefficients are the first row of **Ω** for the MS or the ME mating system.

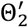: identical to HSΘ1-3, and thus 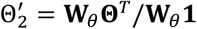 for MS or ME.

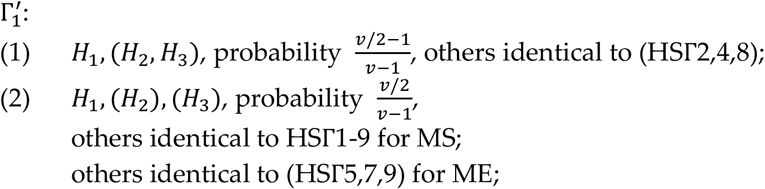

then 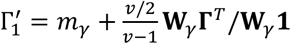 for MS or 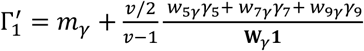 for ME, where

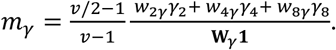

In this way, the expressions of other double non-identity coefficients can be written down, so can the elements in the corresponding rows of **Ω**. We will omit these lengthy explanations from the following discussion.

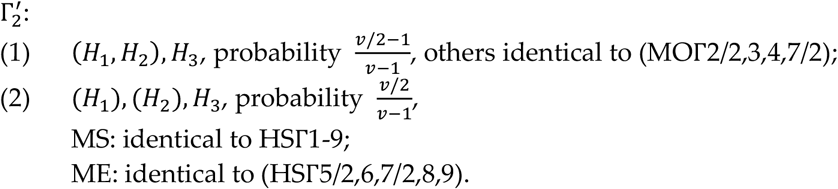

Here *‘others’* is omitted for brevity, the same below.

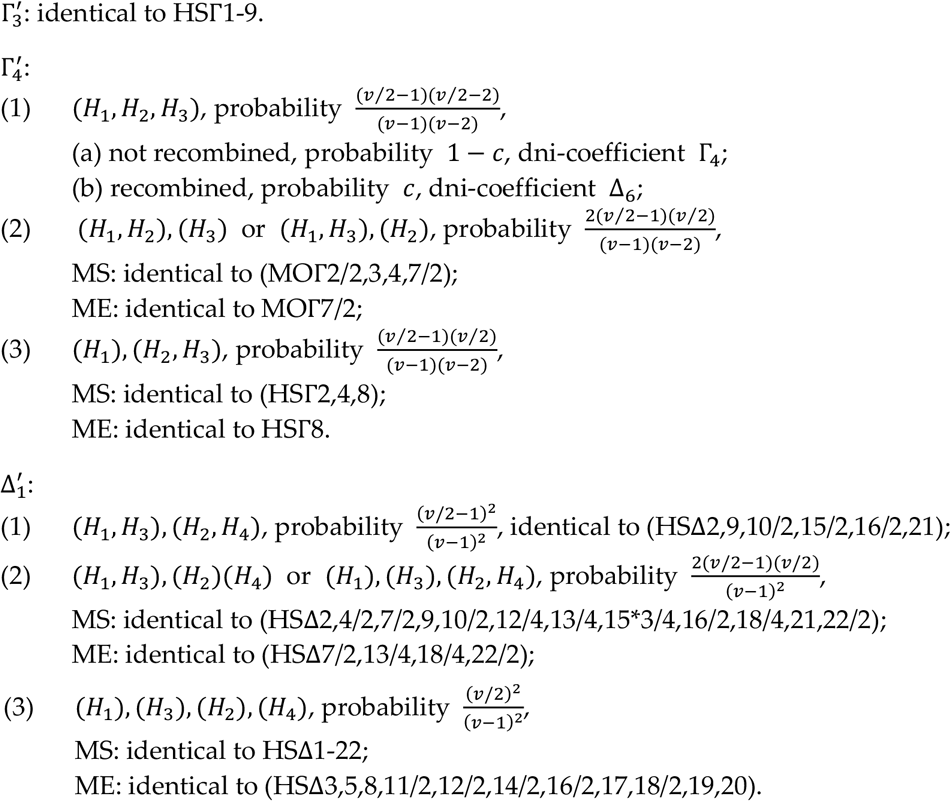

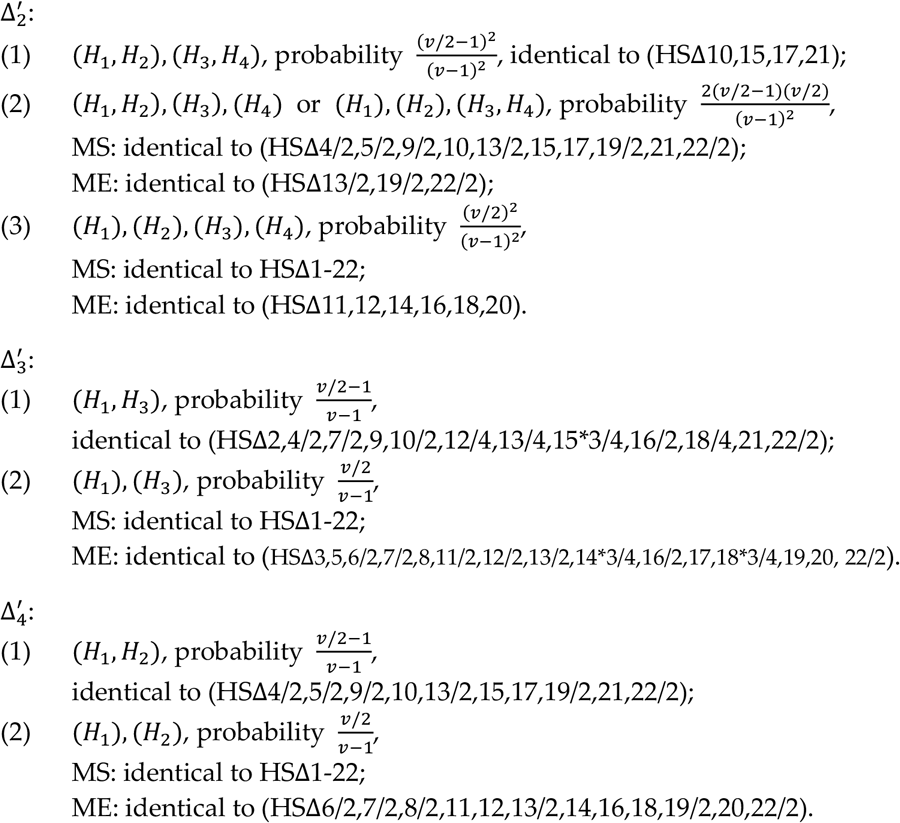

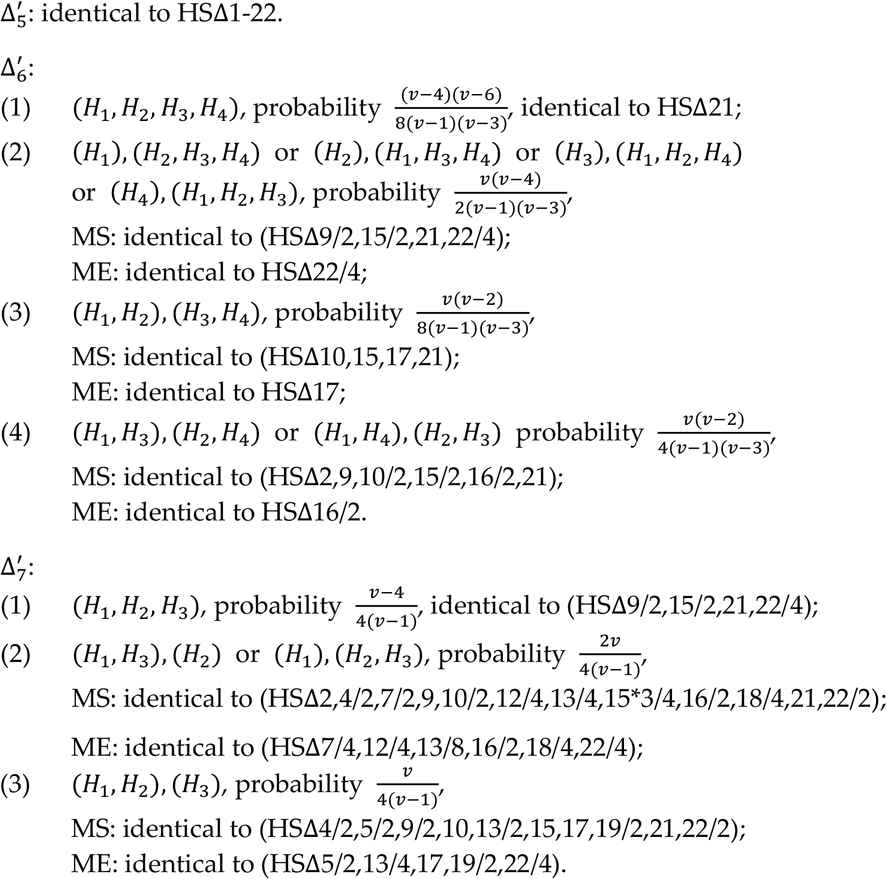

The transition matrix **Ω** for the MS or the ME mating system is not shown, but the matrices **T** and **S** in the principal part of **Ω** are listed in Appendix H.

### Appendix F. DR mating system

In the dioecious mating systems, no matter whether it is DR or DH, each individual is formed by a sperm and an egg that are independently sampled from the sperm and the egg pools, respectively. We will discuss the double non-identity coefficients one by one for the DR mating system in this appendix.

For simplicity, we will use the symbol { * } to be the identifier of an item, e.g. the item {HSΘ2} means that its contents are the same as those in HSΘ2 except for the weight, and we also use the symbol 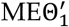 to represent 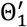 in ME, and so on.

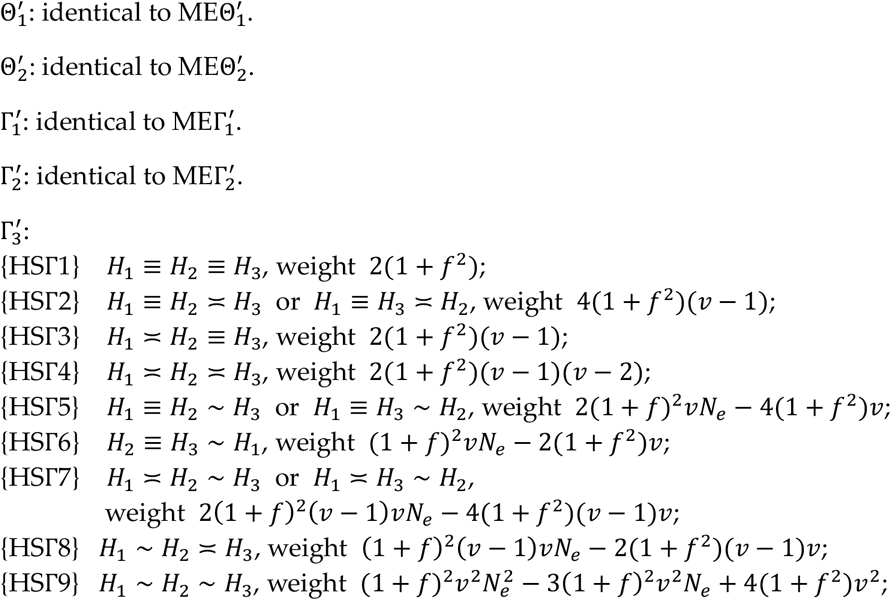

then 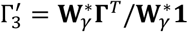 where 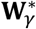 is the row vector consisting of the above nine weights. 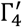: identical to 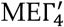.

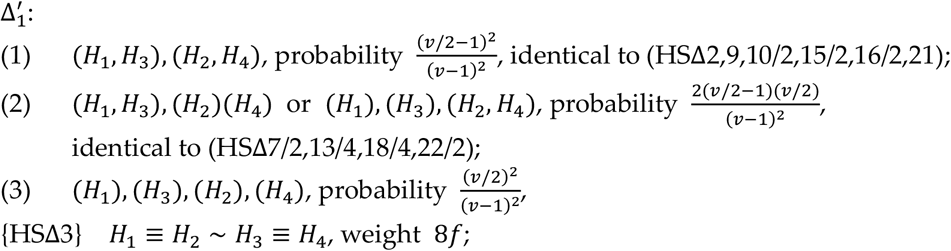

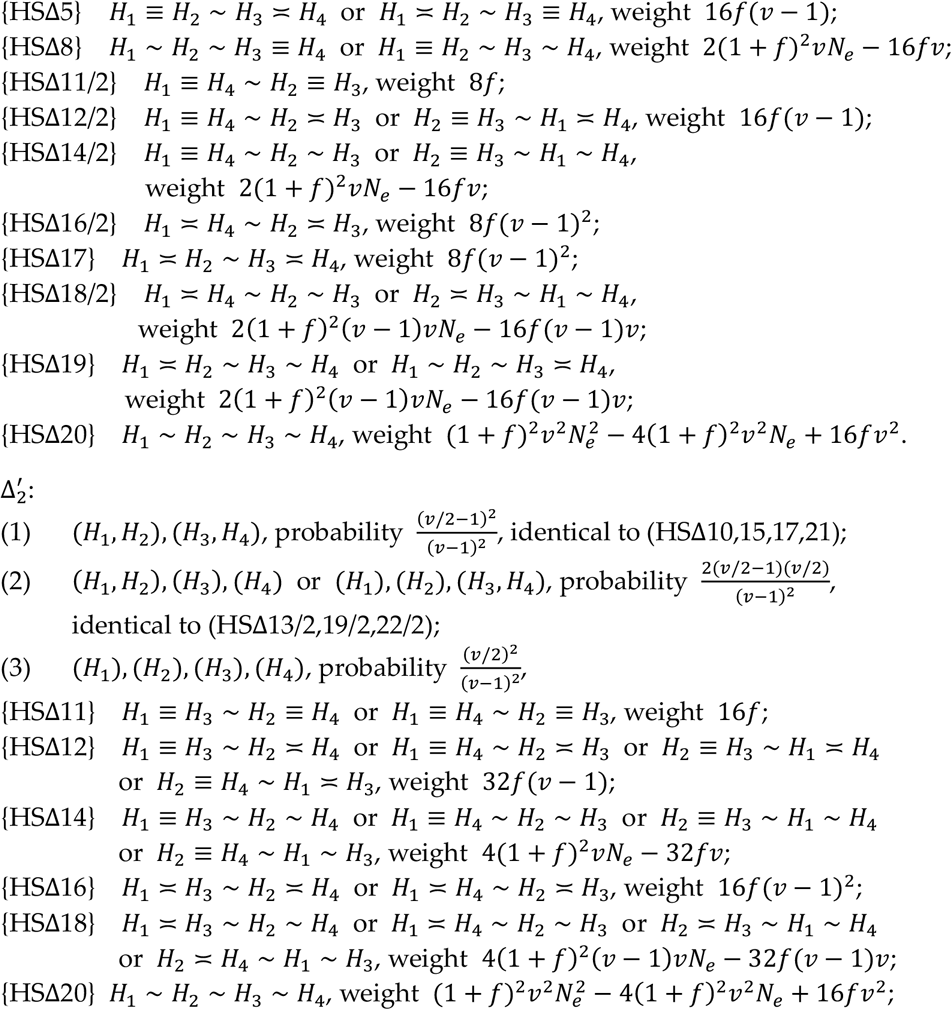

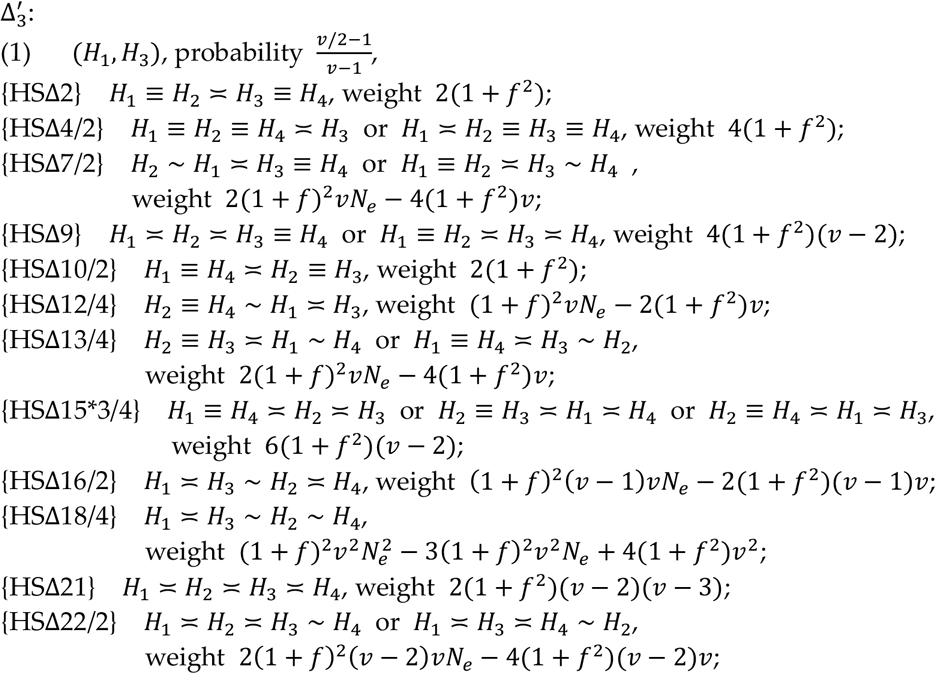

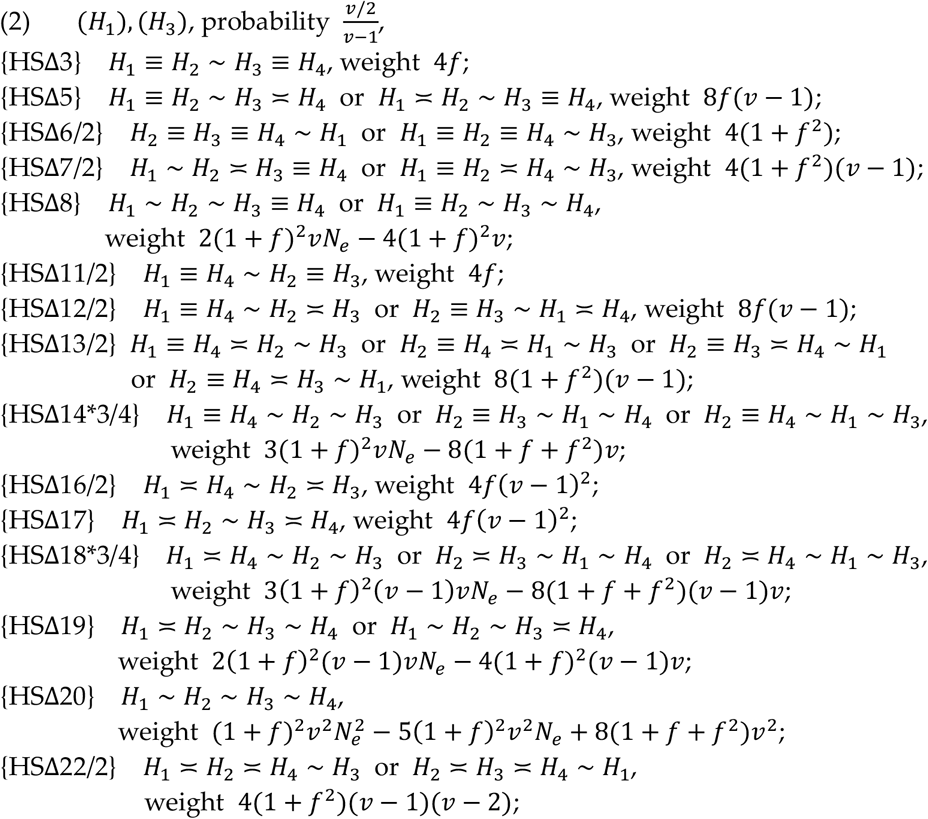

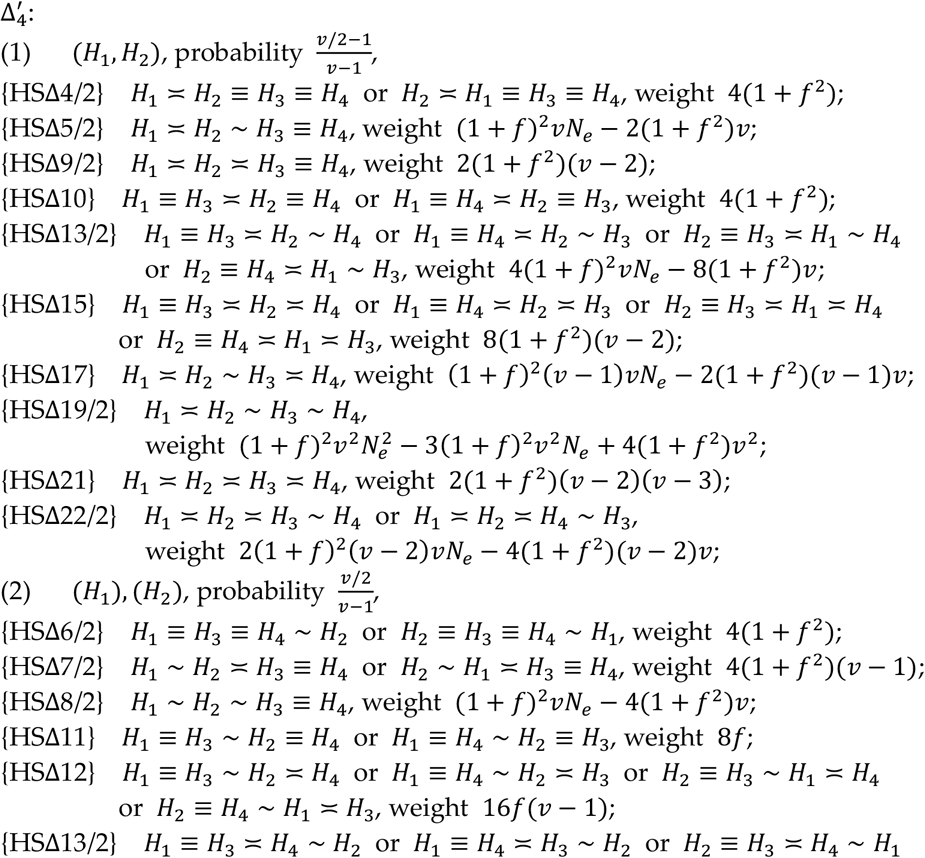

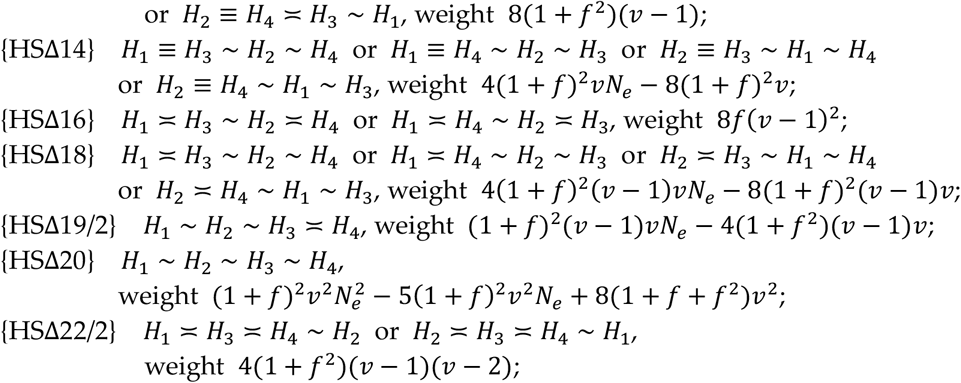

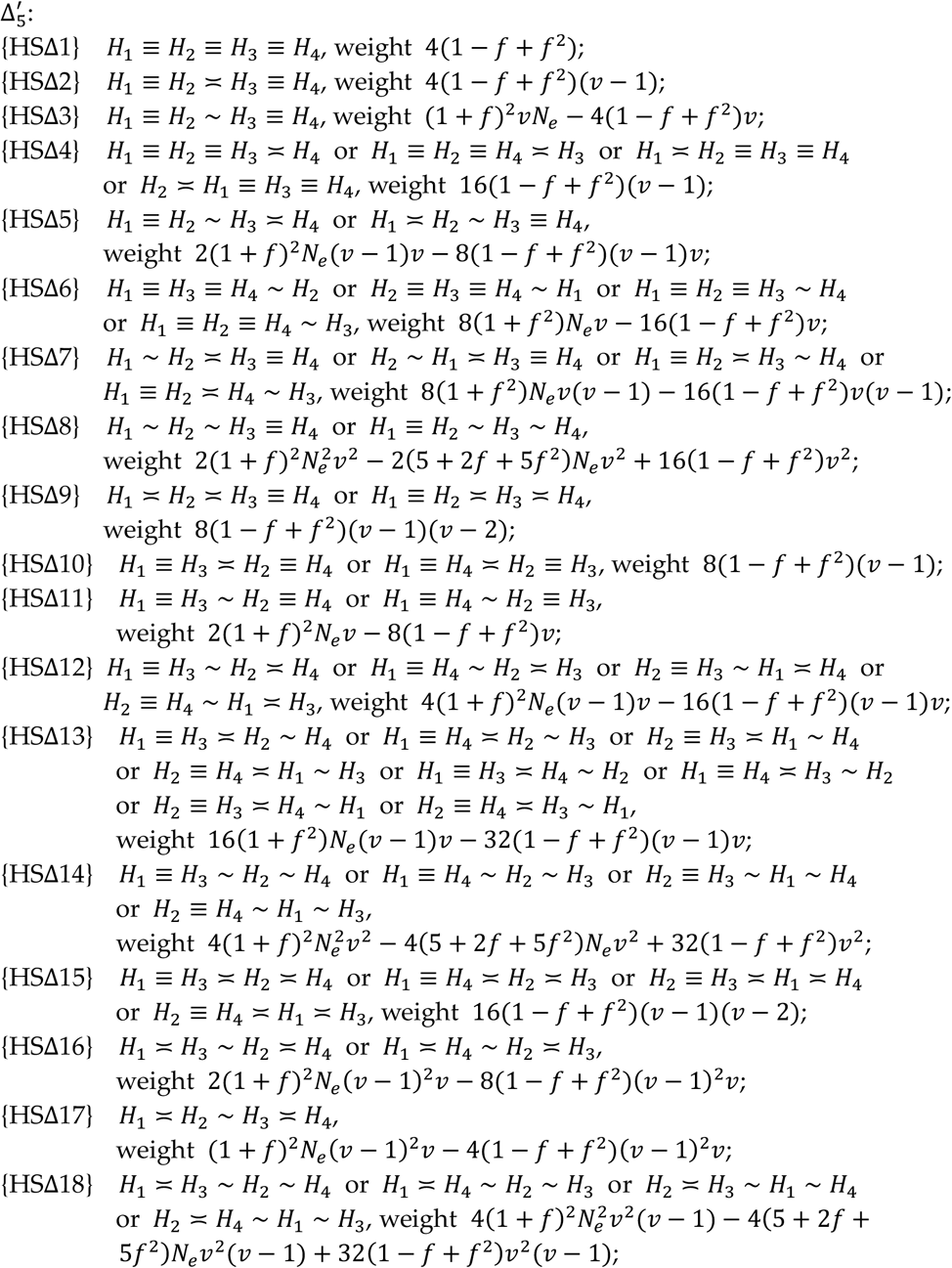

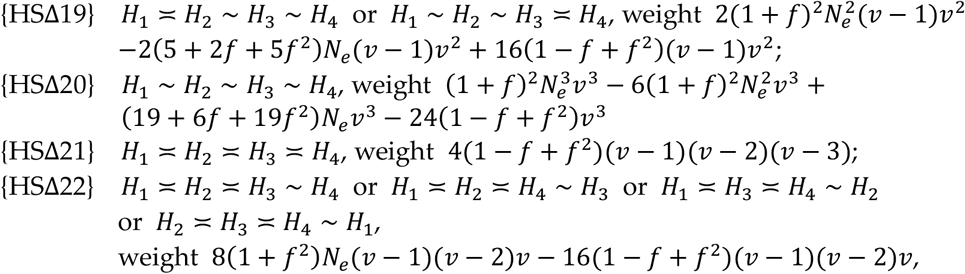

then 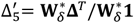 where 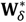 is the row vector consisting of the above 22 weights.

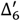: identical to 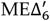.
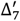: identical to 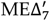.

The transition matrix **Ω** for the DR mating system is not shown, but the matrices **T** and **S** in the principal part of **Ω** are listed in Appendix H.

### Appendix G. DH mating system

For the DH mating system, because each individual remains in a reproductive unit for its entire lifetime, the offspring produced within each reproductive unit are either full-or half-sibs. We will denote [*H, H*’, …] for which those haplotypes within square brackets are from the same reproductive unit.

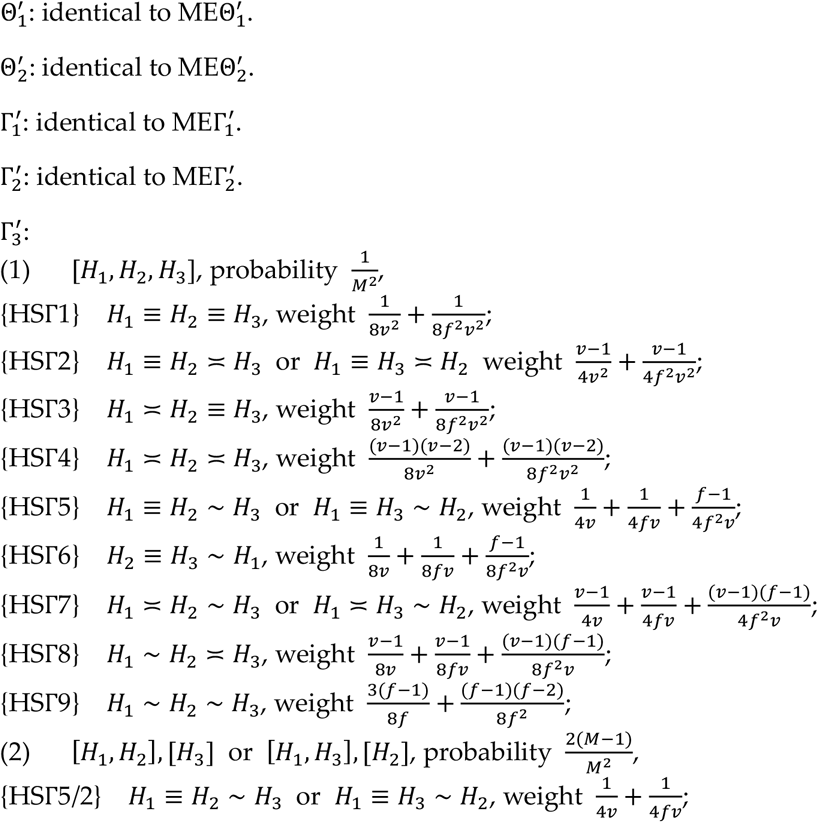

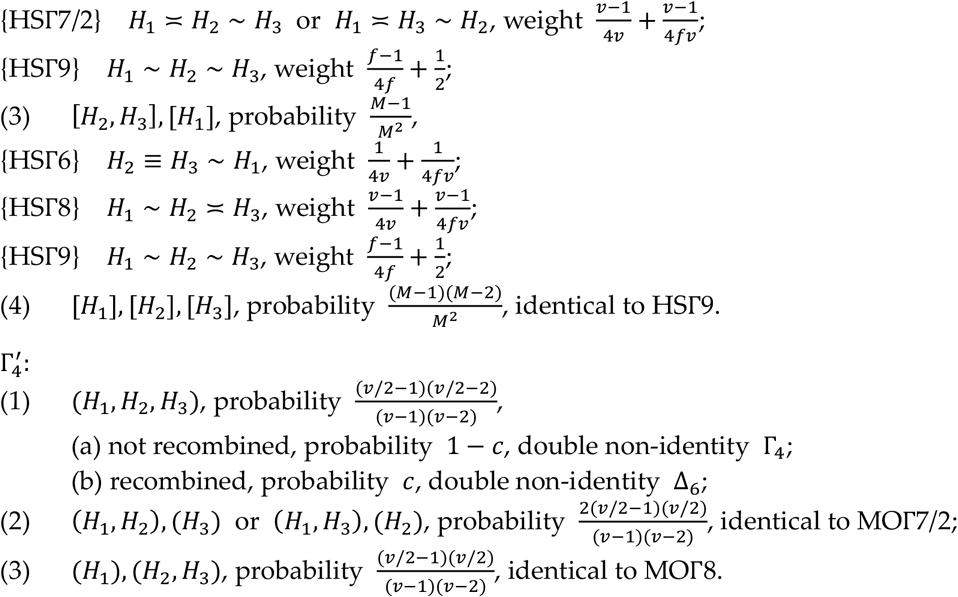

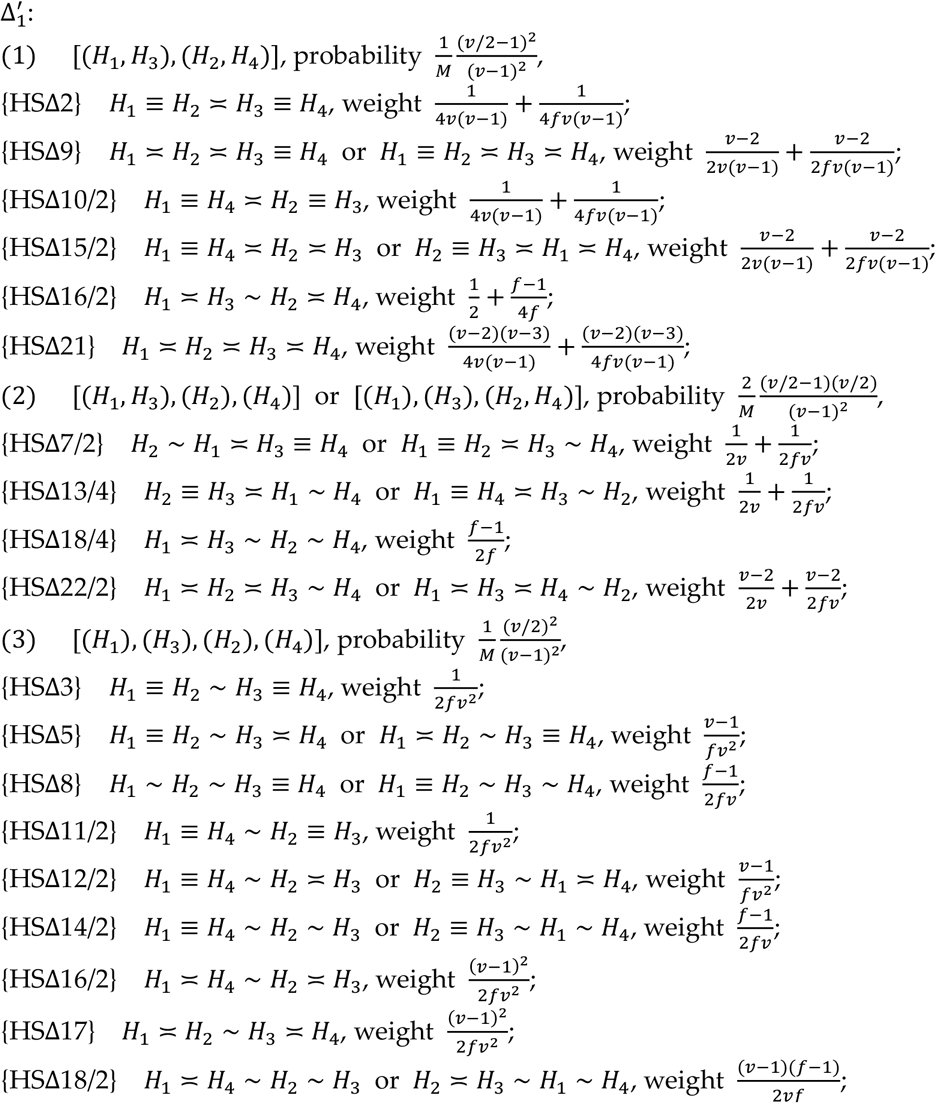

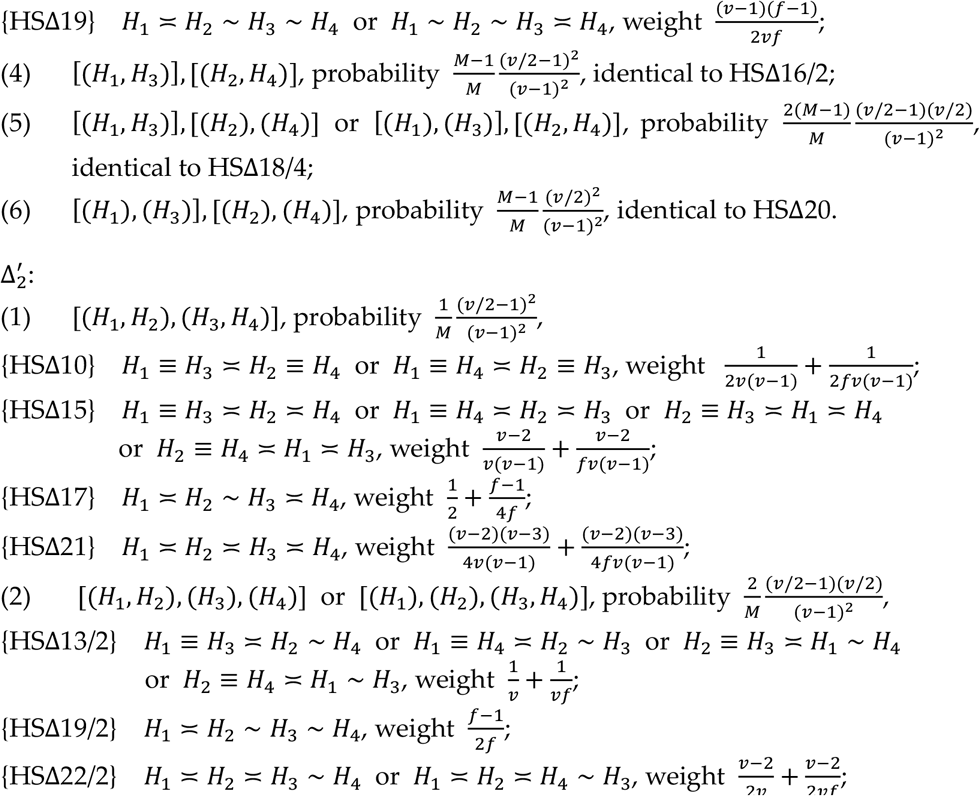

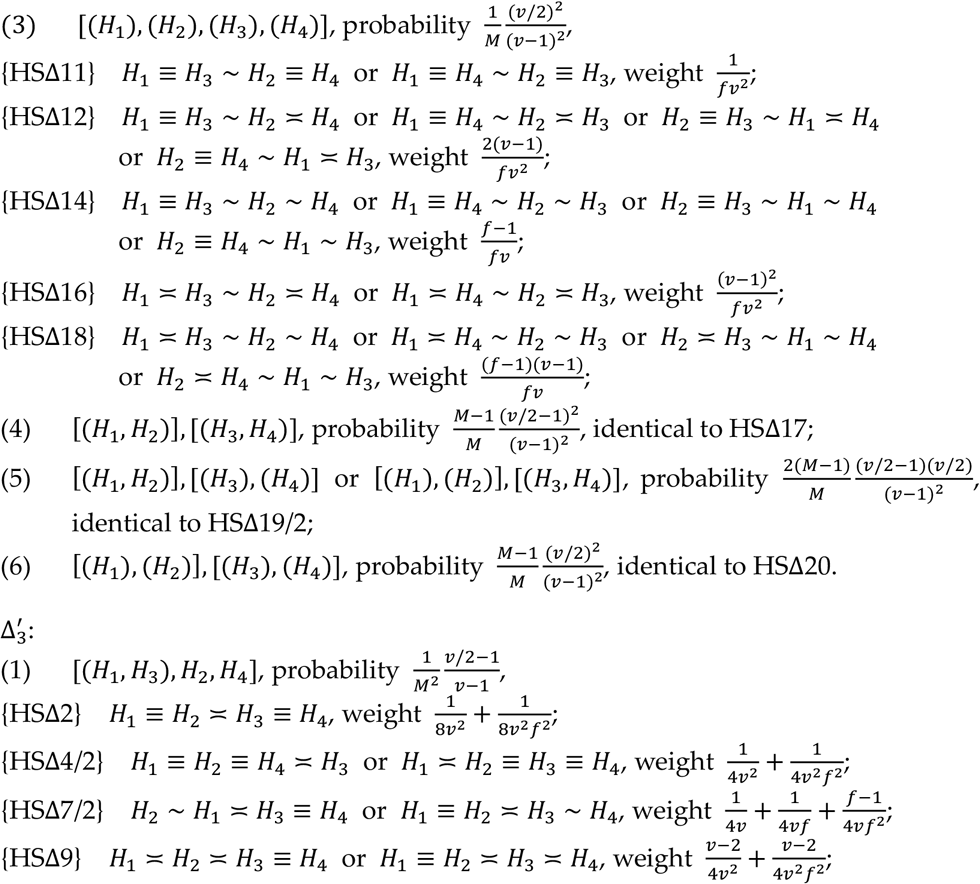

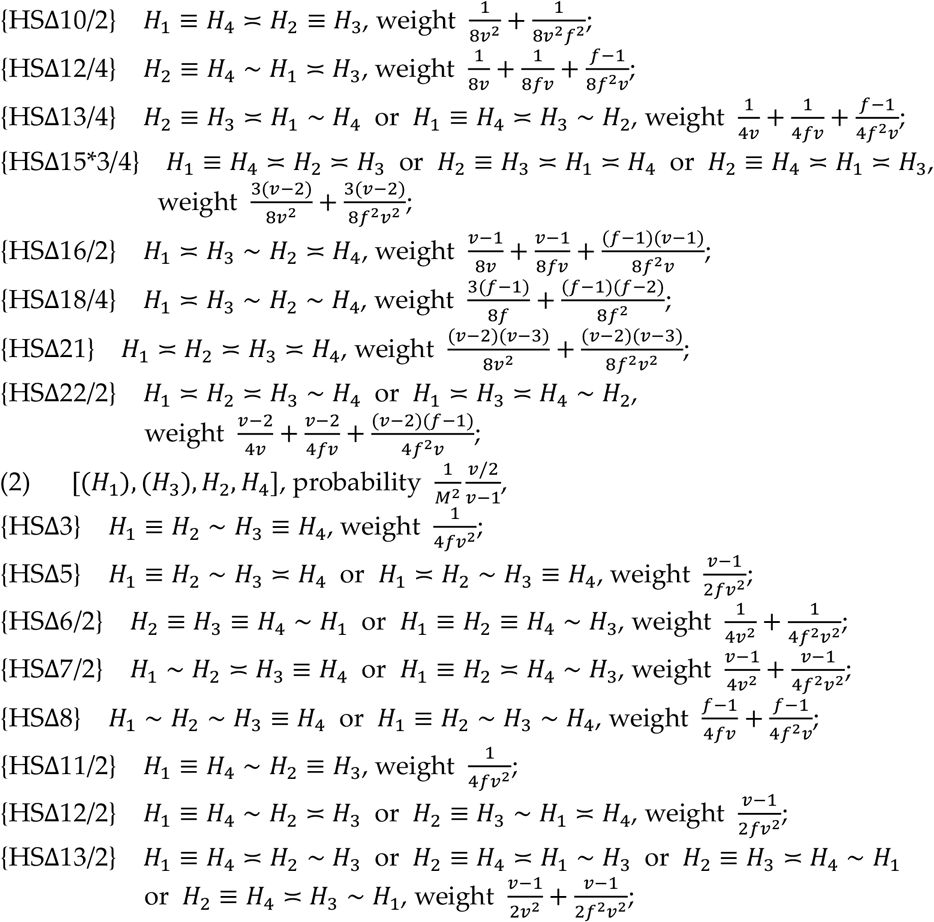

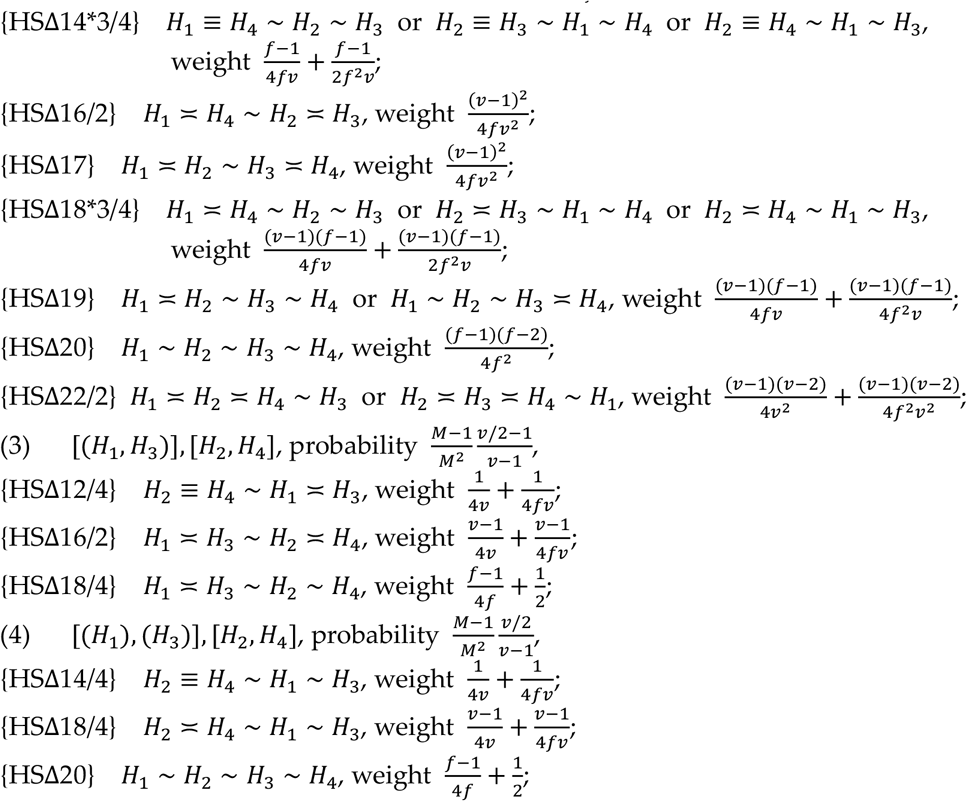

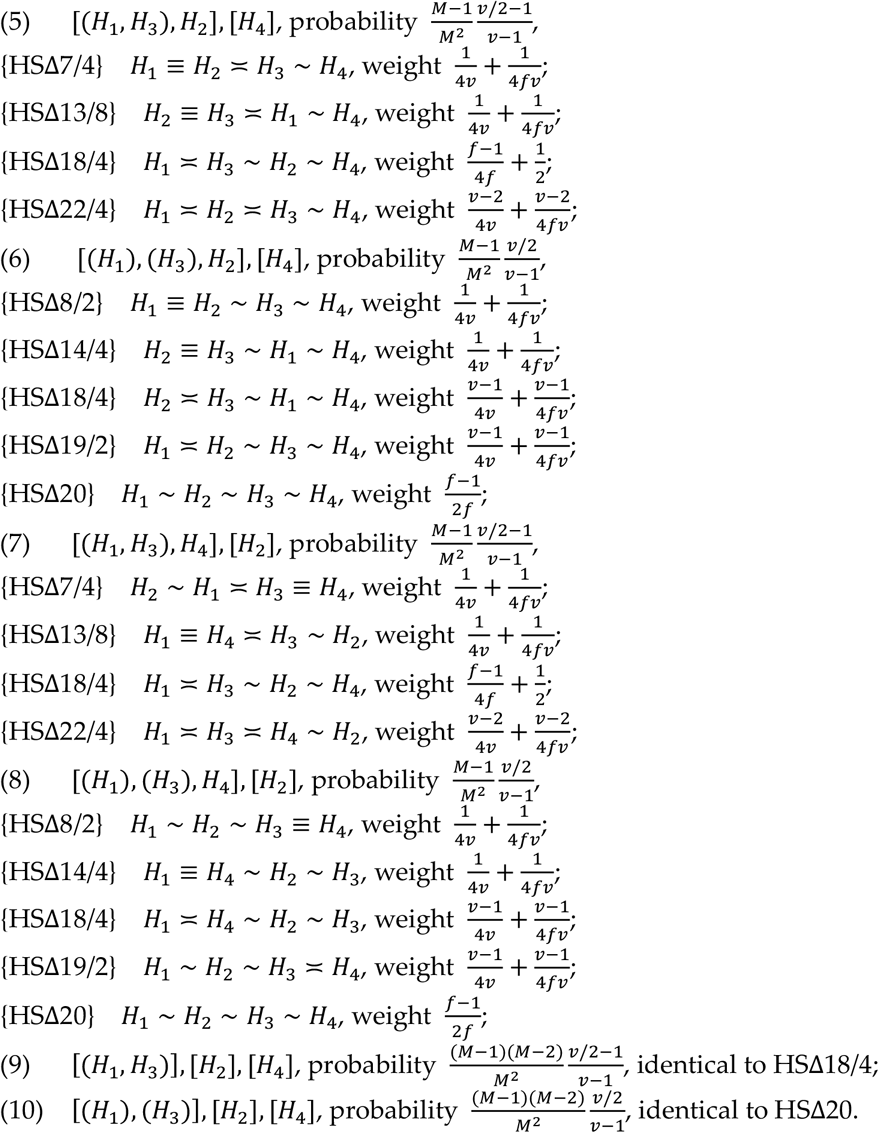

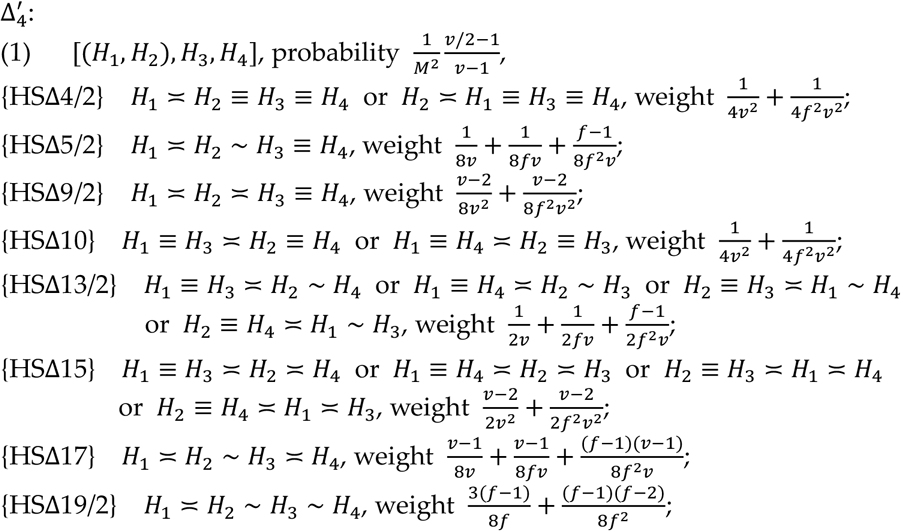

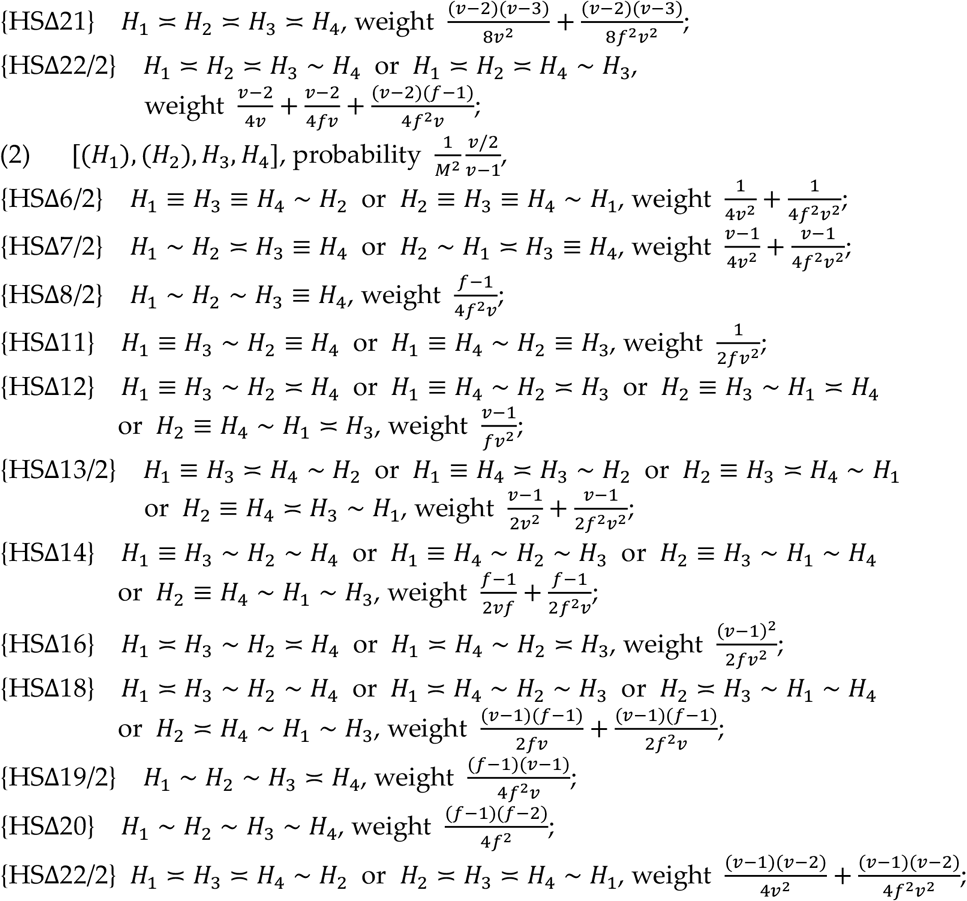

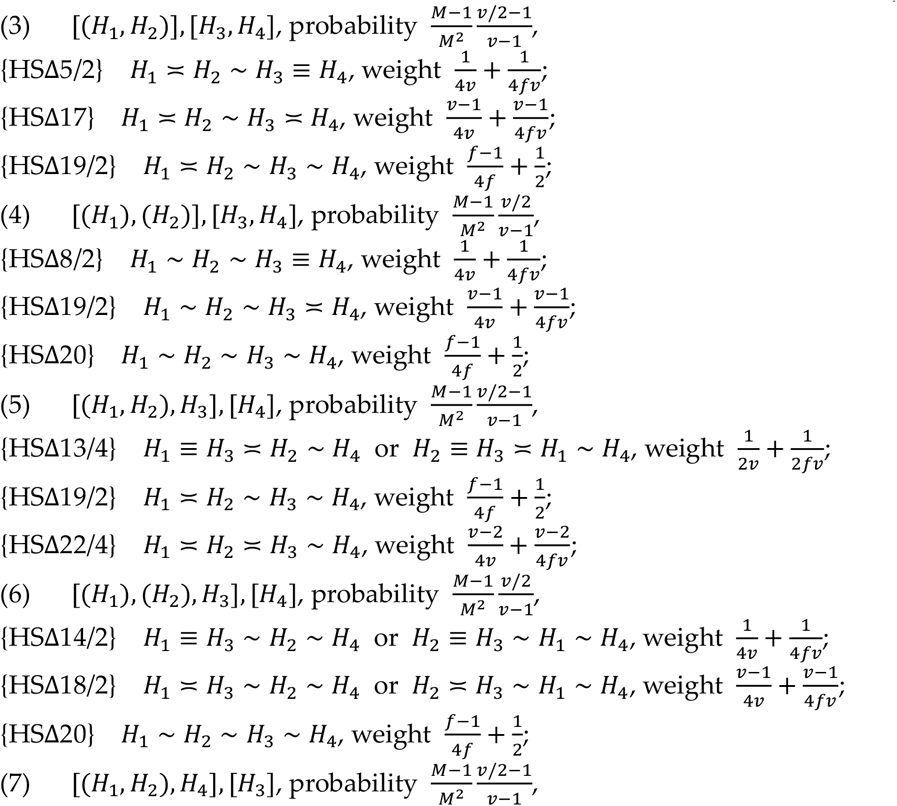

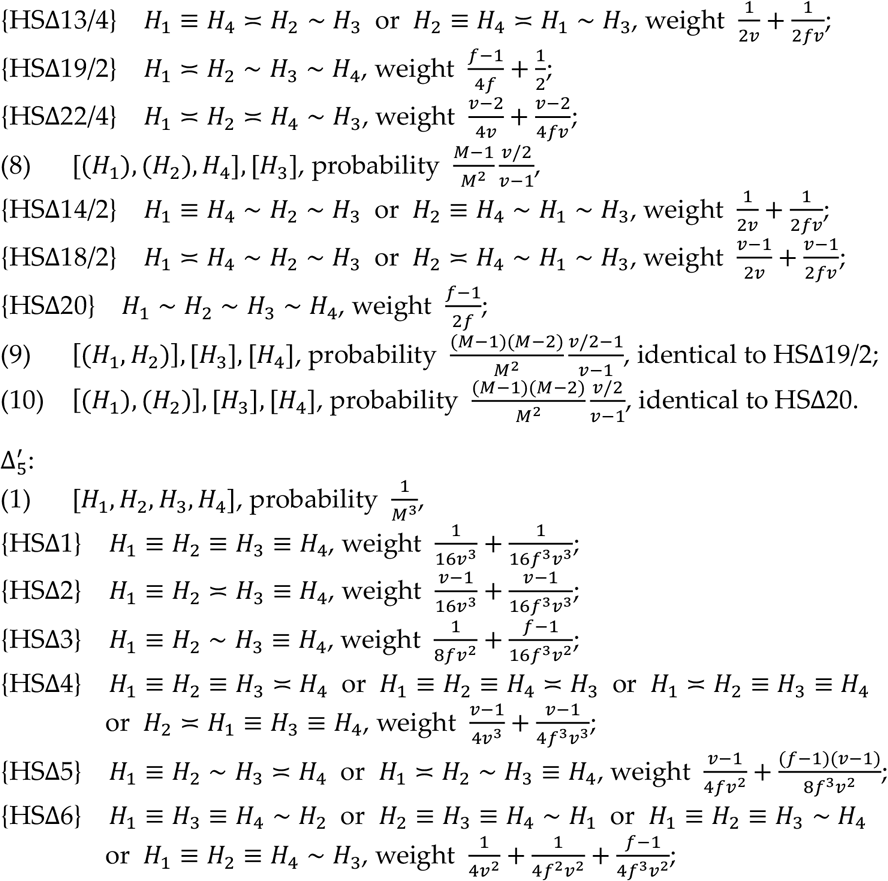

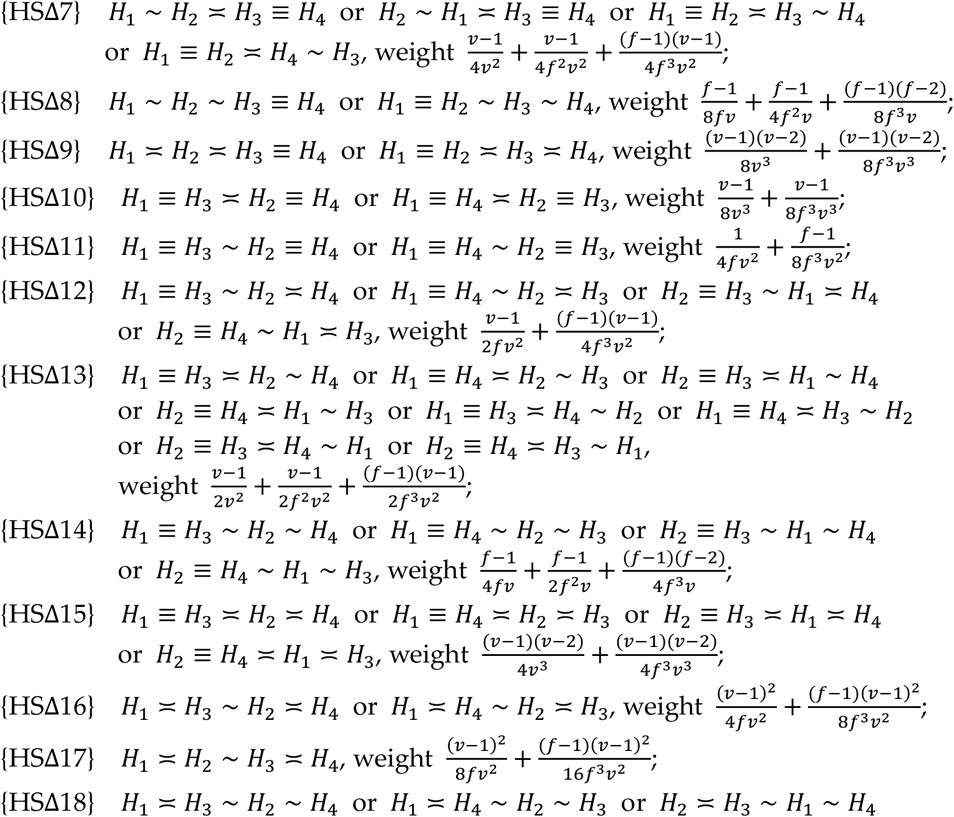

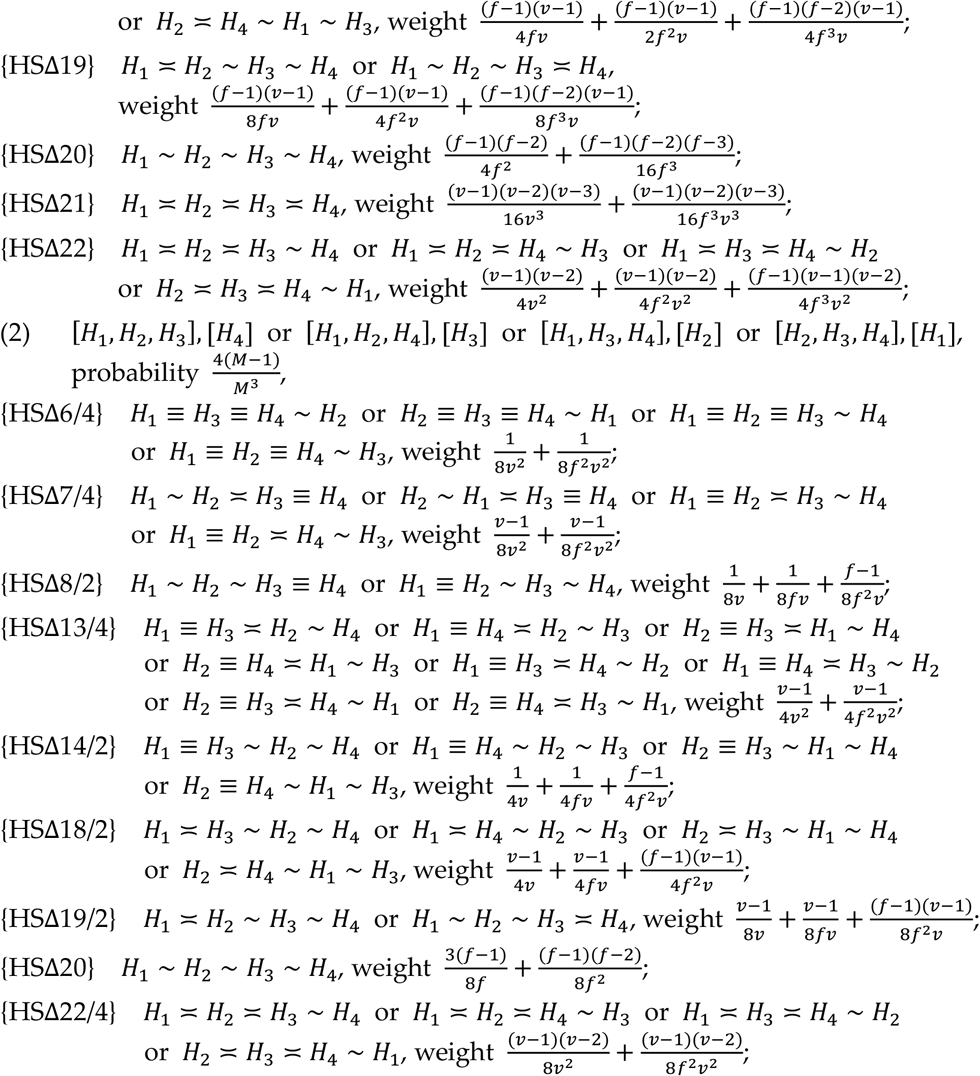

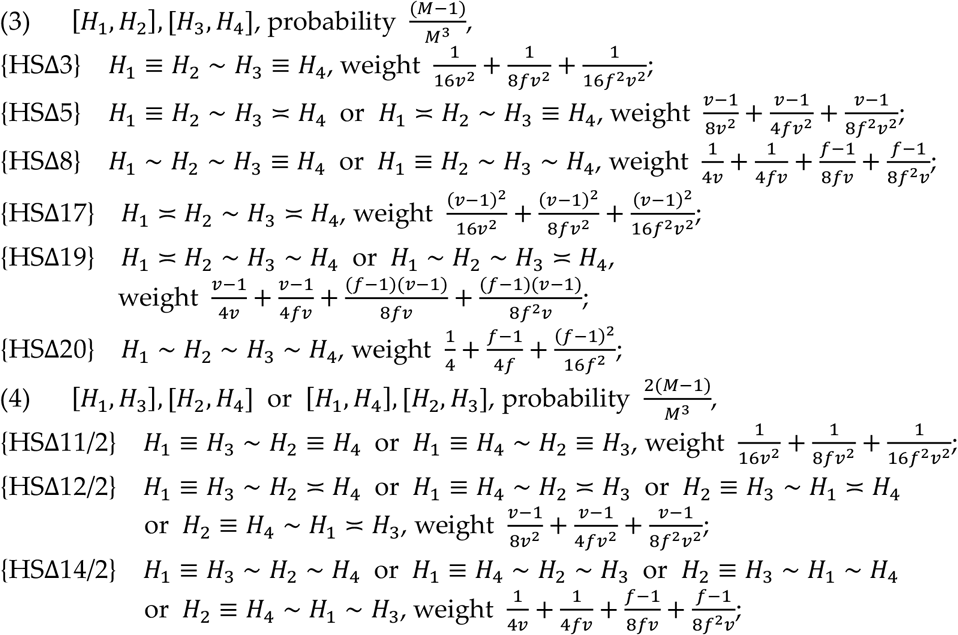

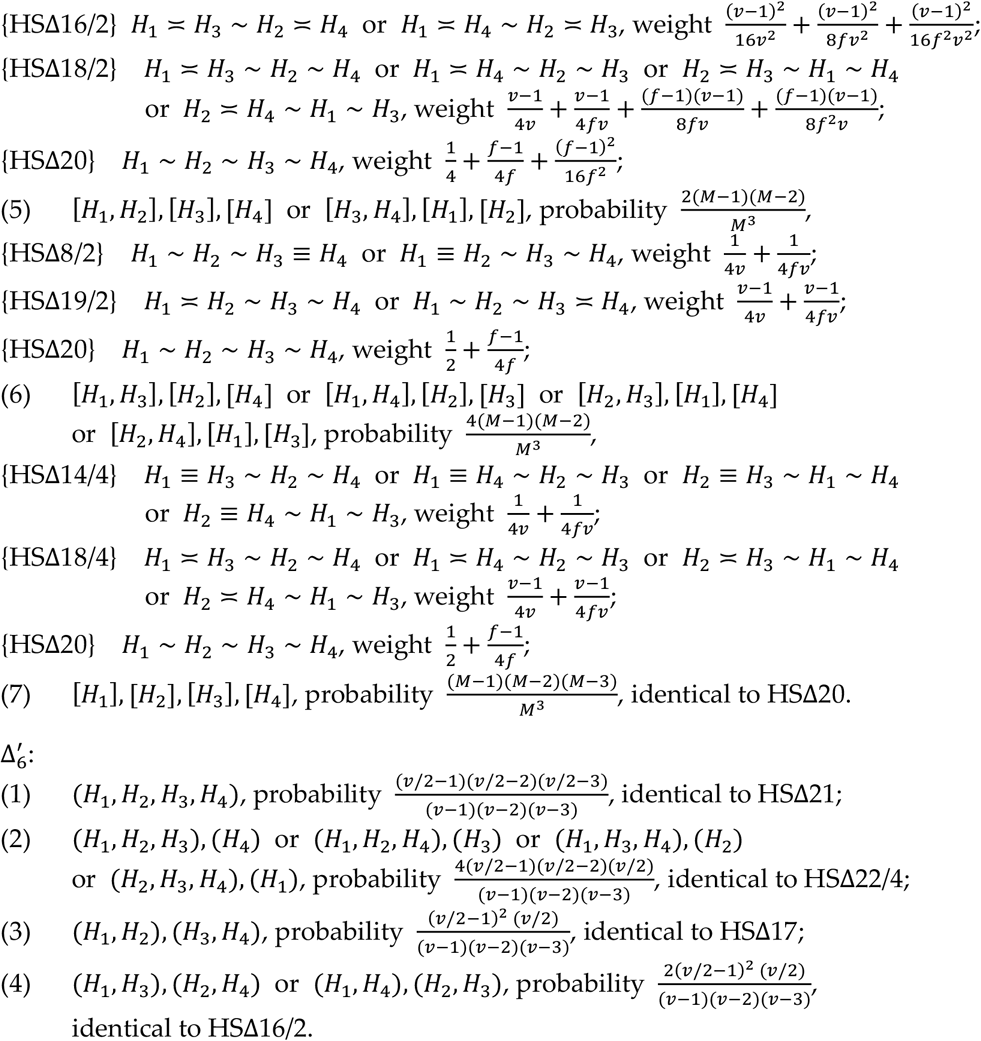

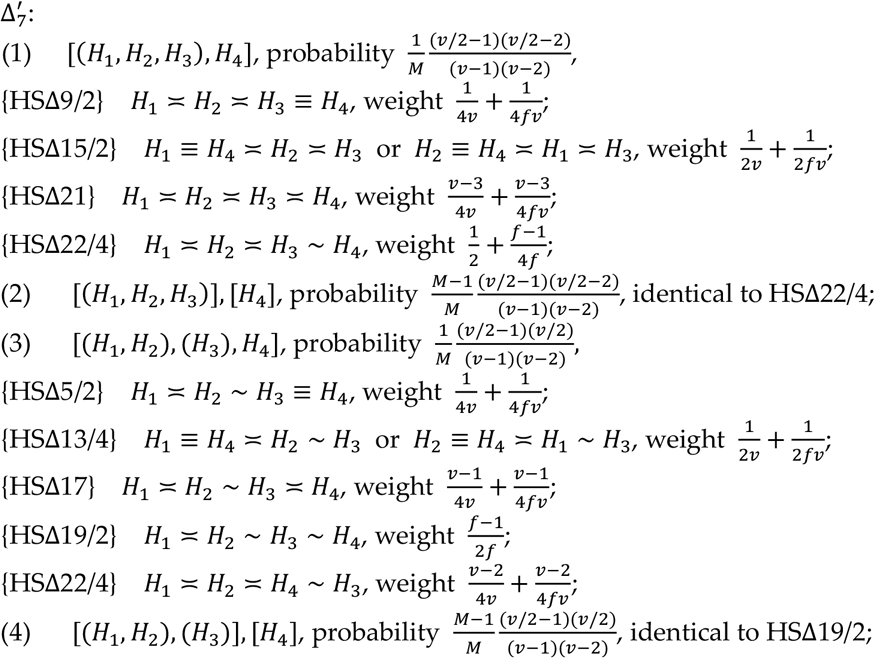

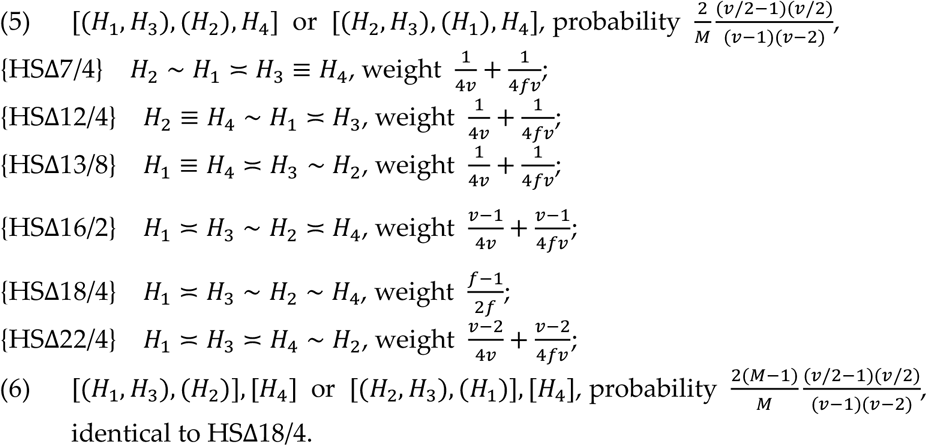

The transition matrix **Ω** for the DH mating system is not shown, but the matrices **T** and **S** in the principal part of **Ω** are listed in Appendix H.

### Appendix H. Matrices T and S for various mating systems

**HS mating system**

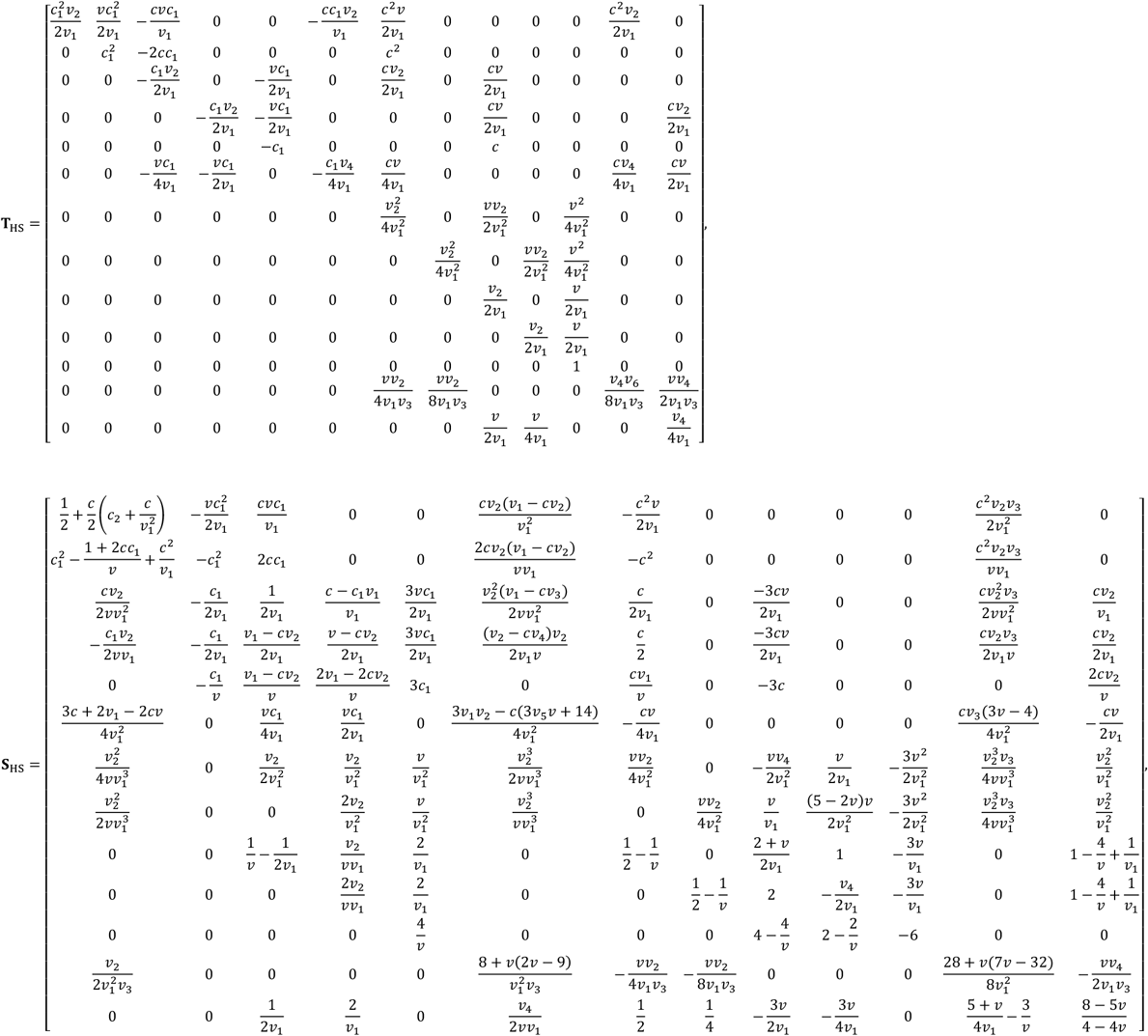

**MS mating system**

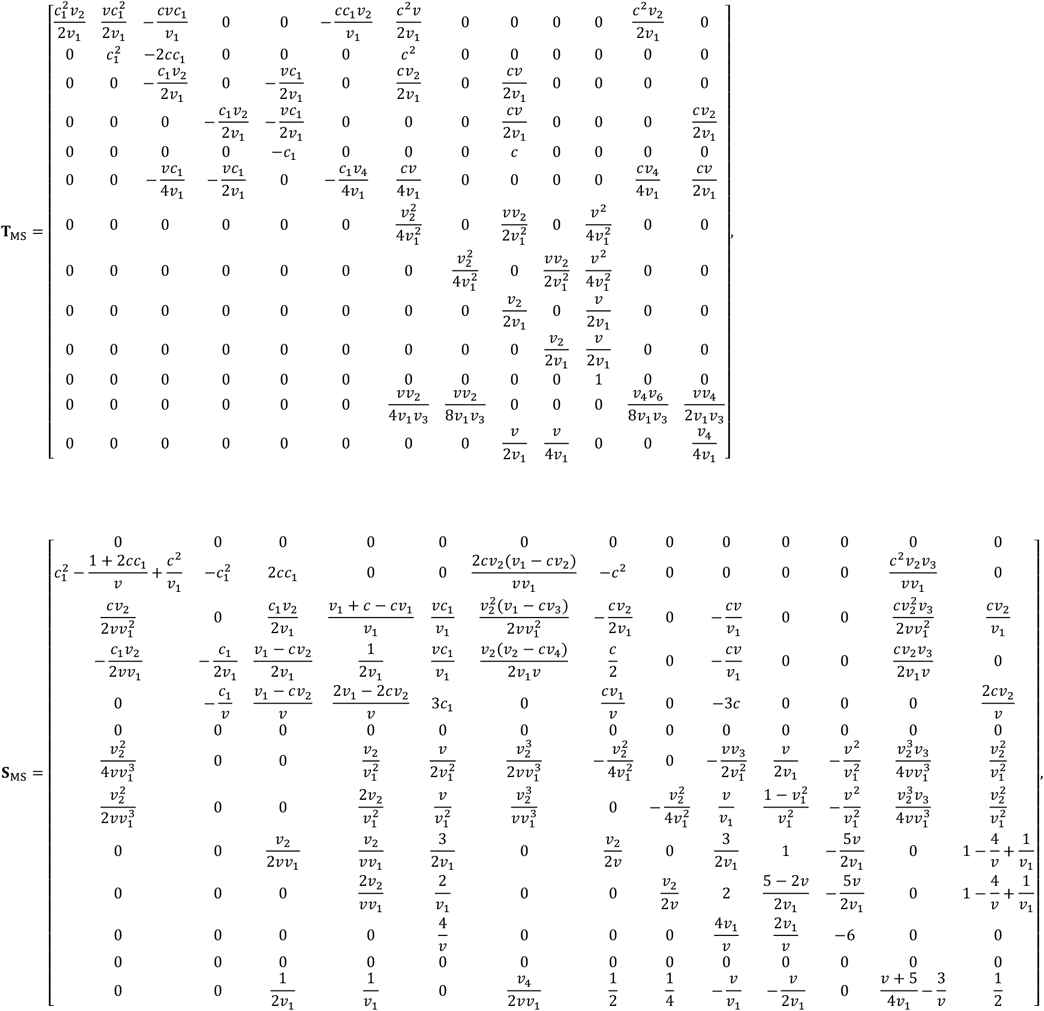

**ME and DR mating systems**

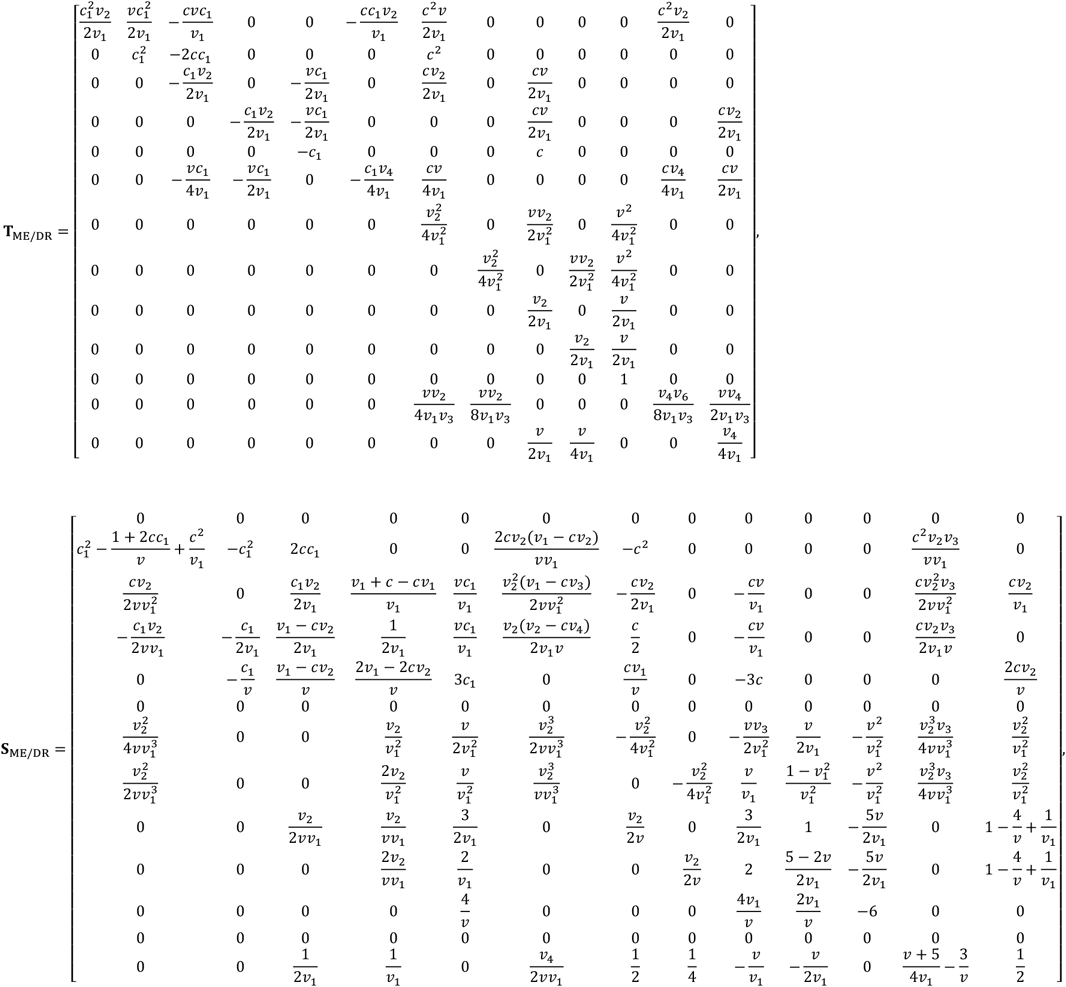

**DH mating system**

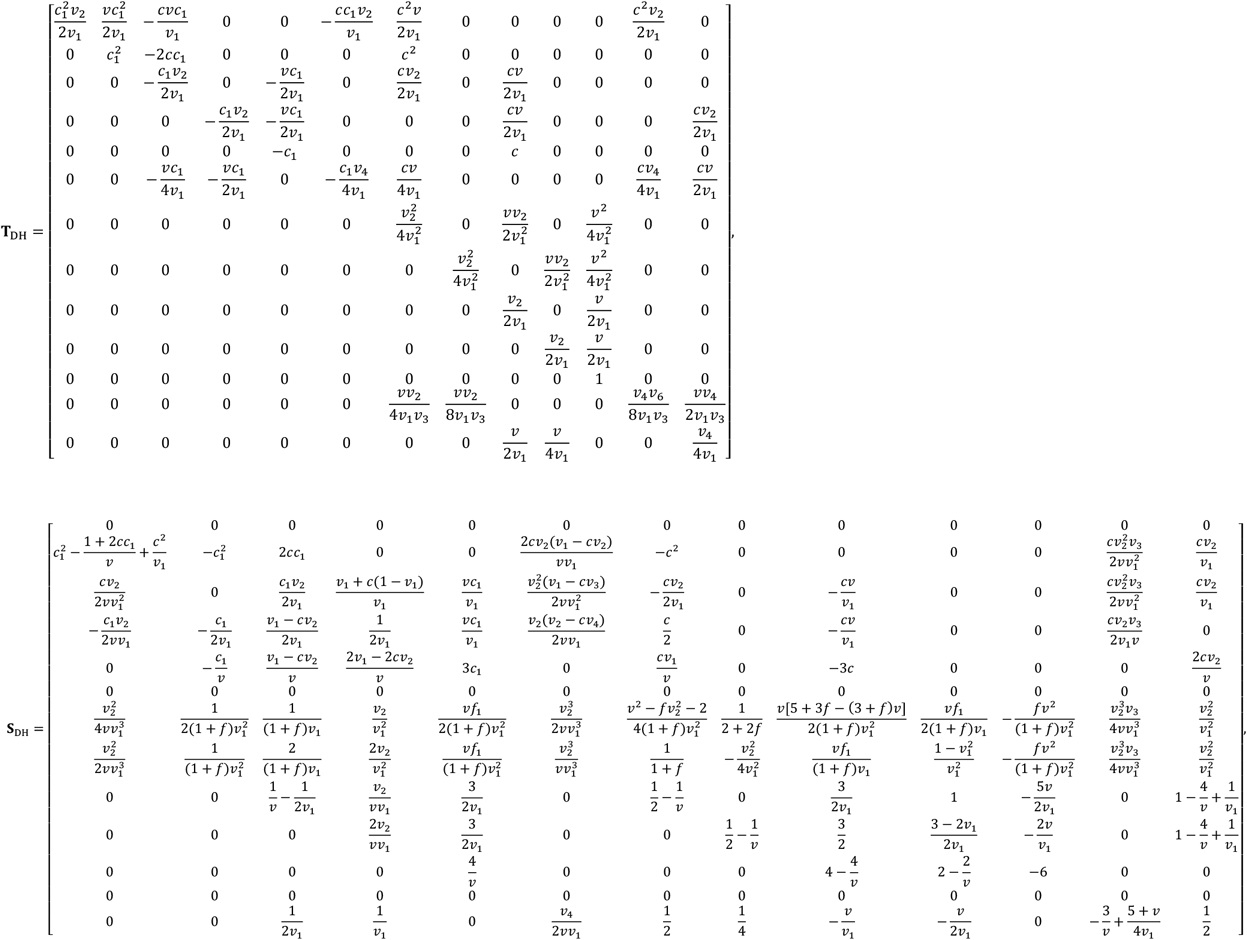

### Appendix I. Bias correction for *D* and Δ

If the allele frequency is estimated from the samples, the estimates 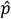 and 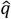 are unbiased, but their product 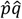 is biased. Because *pq* appears in the expressions of *D* and Δ, we should consider the correction of biases during estimating *D* and Δ. In this appendix, we will follow the method for the unbiased estimation of Weir (1979) to give the unbiased estimate formulas for *D* and Δ, where *D*, Δ, *p* and *q* are short for *D_AB_*, *Δ*_*AB*_, *q_A_* and *q_B_* in turn.

We begin our discussion with the expectation 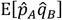. According to the facts that 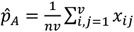 and 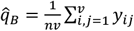, we have

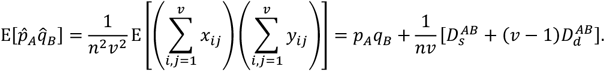

For the case of phased genotypes, since 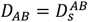, the following holds:

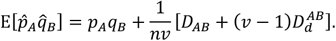

In addition, 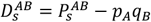 and 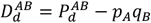, we obtain

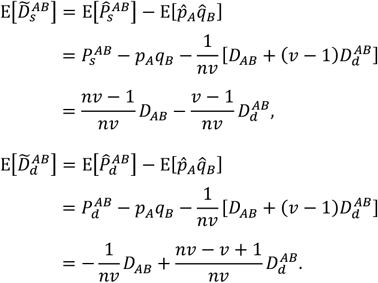

Replacing 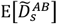 by 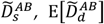 by 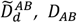 by 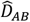 and 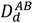 by 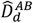, it follows

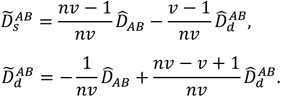

This is a linear equation set with 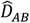 and 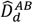 as the unknowns, the solution of which for 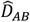 is the unbiased estimate for *D_AB_*, and its expression is as follows:

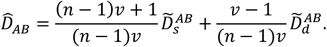

For the case of unphased genotypes, by 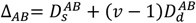, the following holds:

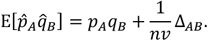

In addition, 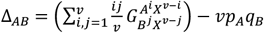 by Equation (1). Hence

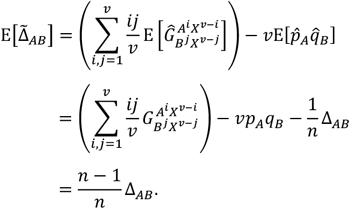

Now, replacing 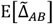 by 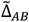 and Δ_*AB*_ by 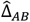, it follows the unbiased estimate formula for Δ_*AB*_ as follows:

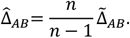

### Supplementary Tables

**Table S1.** Elements in combination matrix *A*_1_

| | $E(\widehat{D}_w^2)$ | $E(\widehat{D}_b^2)$ | $E(\widehat{D}_w\widehat{D}_b)$ | $E(\widehat{D}^2)$ | $E(\widehat{\Delta}^2)$ | $E(\widehat{Q})$ | $E(\widehat{R})$ |
| --- | --- | --- | --- | --- | --- | --- | --- |
| $\Theta_1$ | 0 | 0 | 0 | 0 | 0 | 0 | 0 |
| $\Theta_2$ | 1 | 0 | 0 | 1 | 1 | 0 | 0 |
| $\Gamma_1$ | -2 | 0 | 1 | 0 | $2v_1$ | 0 | 0 |
| $\Gamma_2$ | 0 | 0 | 0 | 0 | 0 | 0 | 0 |
| $\Gamma_3$ | 0 | 0 | -1 | -2 | $-2v$ | 0 | 0 |
| $\Gamma_4$ | 0 | 0 | 0 | 0 | 0 | 0 | 0 |
| $\Delta_1$ | 1 | 1 | -1 | 0 | $v_1^2$ | 0 | 0 |
| $\Delta_2$ | 0 | 0 | 0 | 0 | 0 | 0 | $v_1^2$ |
| $\Delta_3$ | 0 | -2 | 1 | 0 | $-2v_1v$ | 0 | 0 |
| $\Delta_4$ | 0 | 0 | 0 | 0 | 0 | 0 | $-2v_1v$ |
| $\Delta_5$ | 0 | 1 | 0 | 1 | $v^2$ | 1 | $v^2$ |
| $\Delta_6$ | 0 | 0 | 0 | 0 | 0 | 0 | 0 |
| $\Delta_7$ | 0 | 0 | 0 | 0 | 0 | 0 | 0 |

**Table S2.** Elements in combination matrix *A*_2_

| | $E(\widehat{D}_w^2)$ | $E(\widehat{D}_b^2)$ | $E(\widehat{D}_w\widehat{D}_b)$ | $E(\widehat{D}^2)$ | $E(\widehat{\Delta}^2)$ | $E(\widehat{Q})$ | $E(\widehat{R})$ |
| --- | --- | --- | --- | --- | --- | --- | --- |
| $\Theta_1$ | $\frac{v_1^2 + 1}{vv_1}$ | $\frac{1}{vv_1}$ | $-\frac{1}{vv_1}$ | $\frac{v_1}{v}$ | $\frac{2v_1}{v}$ | 0 | $\frac{2v_1}{v}$ |
| $\Theta_2$ | -1 | 0 | $-\frac{1}{v}$ | $-\frac{2+v}{v}$ | -3 | 0 | 0 |
| $\Gamma_1$ | 2 | $-\frac{2}{v}$ | $-\frac{2v_1}{v}$ | $-\frac{2v_1}{v}$ | $-6v_1$ | 0 | 0 |
| $\Gamma_2$ | 0 | $-\frac{4}{v}$ | $-\frac{2v_2}{v}$ | $-\frac{4v_1}{v}$ | $-8v_1$ | 0 | $-8v_1$ |
| $\Gamma_3$ | 0 | $\frac{4}{v}$ | 3 | $\frac{4+6v}{v}$ | $10v$ | $\frac{4}{v}$ | $4v$ |
| $\Gamma_4$ | $-\frac{2v_2^2}{vv_1}$ | $\frac{2v_2}{vv_1}$ | $\frac{v_2v_3}{vv_1}$ | 0 | $\frac{4v_1v_2}{v}$ | 0 | $\frac{4v_1v_2}{v}$ |
| $\Delta_1$ | -1 | $\frac{2-3v}{v}$ | $\frac{2v-1}{v}$ | 0 | $-3v_1^2$ | 0 | 0 |
| $\Delta_2$ | 0 | 0 | 0 | 0 | 0 | 0 | $-3v_1^2$ |
| $\Delta_3$ | 0 | $\frac{10v-4}{v}$ | -3 | $\frac{4v_1}{v}$ | $10vv_1$ | $\frac{4v_1}{v}$ | $4vv_1$ |
| $\Delta_4$ | 0 | $\frac{2v_1}{v}$ | 0 | $\frac{2v_1}{v}$ | $2vv_1$ | $\frac{2v_1}{v}$ | $8vv_1$ |
| $\Delta_5$ | 0 | -6 | 0 | -6 | $-6v^2$ | -6 | $-6v^2$ |
| $\Delta_6$ | $\frac{v_2v_3}{vv_1}$ | $\frac{v_2v_3}{vv_1}$ | $-\frac{v_2v_3}{vv_1}$ | 0 | $\frac{v_1v_2v_3}{v}$ | 0 | $\frac{v_1v_2v_3}{v}$ |
| $\Delta_7$ | 0 | $-\frac{4v_2}{v}$ | $\frac{2v_2}{v}$ | 0 | $-4v_1v_2$ | 0 | $-4v_1v_2$ |

**Table S3.** Essential factors to form Ω^*T*^ for HS mating system

| | $\Theta'_1$ to $\Theta'_2$ | $\Gamma'_1$ to $\Gamma'_4$ | $\Delta'_1$ to $\Delta'_7$ |
| --- | --- | --- | --- |
| $\Theta_1$ | $c^2 + c_1^2 v_1^2$ | $v_1 - cv_2$ | $2v_1$ |
| $\Theta_2$ | $vc_1^2 N_1 v_1$ | $-c_1 N_1 v$ | $2vN_1$ |
| $\Gamma_1$ | $-2cvc_1 N_1 v_1$ | $vN_1(v_1 - cv_2)$ | $4vN_1 v_1$ |
| $\Gamma_2$ | 0 | $2vN_1(v_1 - cv_2)$ | $8vN_1 v_1$ |
| $\Gamma_3$ | 0 | $-v^2 c_1 N_1 N_2$ | $4v^2 N_1 N_2$ |
| $\Gamma_4$ | $-2cv_2(cv_2 - v_1)$ | $(v_1 - cv_4)v_2$ | $4v_1 v_2$ |
| $\Delta_1$ | $c^2 v N_1 v_1$ | $cvN_1 v_1$ | $2vN_1 v_1^2$ |
| $\Delta_2$ | 0 | 0 | $vN_1 v_1^2$ |
| $\Delta_3$ | 0 | $cv^2 N_1 N_2$ | $4v^2 N_1 N_2 v_1$ |
| $\Delta_4$ | 0 | 0 | $2v^2 N_1 N_2 v_1$ |
| $\Delta_5$ | 0 | 0 | $v^3 N_1 N_2 N_3$ |
| $\Delta_6$ | $c^2 v_2 v_3$ | $cv_2 v_3$ | $v_1 v_2 v_3$ |
| $\Delta_7$ | 0 | $2cvN_1 v_2$ | $4vN_1 v_1 v_2$ |
| Divisor | $Nvv_1$ | $N^2 v^2$ | $N^3 v^3$ |

There are 13 columns for $\Omega^T$ , the first two columns are the same, each of which is $Nvv_1$ times of the combination coefficients of $\Theta'_1$ or $\Theta'_2$ . Similarly, the next four columns are the same, so are the last seven columns. Moreover, $c - 1$ is denoted by $c_1$ , $N - i$ by $N_i$ and $v - i$ by $v_i$ , $i = 1, 2, 3$ .

**Table S4.** Approximations of *d*^2^

| $v$ | HS | MS/ME/DR | DH |
| --- | --- | --- | --- |
| 2 | $\frac{1 - 2c + 2c^2}{4cN_e - 2c^2N_e}$ | $\frac{1 - 2c + 2c^2}{4cN_e - 2c^2N_e}$ | $-\frac{1 + f - 2c(1 + f) + c^2(3 + 2f)}{2(-2 + c)c(1 + f)N_e}$ |
| 4 | $\frac{3 - 6c + 4c^2}{24cN_e - 12c^2N_e}$ | $\frac{16 - 24c + 12c^2 + 5c^3}{128cN_e - 32c^3N_e}$ | $-\frac{16(1 + f) - 24c(1 + f) + 4c^2(5 + 3f) + c^3(3 + 5f)}{32c(-4 + c^2)(1 + f)N_e}$ |
| 6 | $\frac{5 - 10c + 6c^2}{60cN_e - 30c^2N_e}$ | $\frac{63 - 84c + 14c^2 + 32c^3}{126c(6 + c - 2c^2)N_e}$ | $-\frac{63(1 + f) - 84c(1 + f) + 7c^2(5 + 2f) + c^3(26 + 32f)}{126c(-6 - c + 2c^2)(1 + f)N_e}$ |
| 8 | $\frac{7 - 14c + 8c^2}{112cN_e - 56c^2N_e}$ | $\frac{160 - 200c - 10c^2 + 99c^3}{2560cN_e + 640c^2N_e - 960c^3N_e}$ | $-\frac{-10c^2(-3 + f) + 160(1 + f) - 200c(1 + f) + c^3(87 + 99f)}{320c(-8 - 2c + 3c^2)(1 + f)N_e}$ |
| 10 | $\frac{9 - 18c + 10c^2}{180cN_e - 90c^2N_e}$ | $\frac{-325 + 390c + 78c^2 - 224c^3}{650c(-10 - 3c + 4c^2)N_e}$ | $-\frac{-325(1 + f) + 390c(1 + f) + 13c^2(1 + 6f) - 4c^3(51 + 56f)}{650c(-10 - 3c + 4c^2)(1 + f)N_e}$ |

Each expression should be added $\mathbf{1}/(vn)$ in order to consider the sampling variance.

**Table S5.** Approximations of δ^2^

| $v$ | HS | MS/ME/DR | DH |
| --- | --- | --- | --- |
| 2 | $\frac{1 - 2c + 2c^2}{4cN_e - 2c^2N_e}$ | $\frac{-1 + 2c - 2c^2}{2(-2 + c)c(N_e - v^2\eta)}$ | $-\frac{1 + f - 2cf + 2c^2(1 + f)}{2(-2 + c)c(1 + f)(N_e - v^2\eta)}$ |
| 4 | $\frac{3 - 6c + 4c^2}{24cN_e - 12c^2N_e}$ | $\frac{-4 + 3c - 12c^2 + 4c^3}{2c(-4 + c^2)(4N_e - v^2\eta)}$ | $\frac{3c(-5 + f) - 4(1 + f) + 4c^3(3 + f) - 4c^2(5 + 3f)}{2c(-4 + c^2)(1 + f)(4N_e - v^2\eta)}$ |
| 6 | $\frac{5 - 10c + 6c^2}{60cN_e - 30c^2N_e}$ | $\frac{9(-7 - 4c - 26c^2 + 12c^3)}{14c(-6 - c + 2c^2)(9N_e - v^2\eta)}$ | $\frac{9[-7(1 + f) + 12c^3(3 + f) - 2c(27 + 2f) - 2c^2(25 + 13f)]}{14c(-6 - c + 2c^2)(1 + f)(9N_e - v^2\eta)}$ |
| 8 | $\frac{7 - 14c + 8c^2}{112cN_e - 56c^2N_e}$ | $\frac{4(-10 - 19c - 44c^2 + 24c^3)}{5c(-8 - 2c + 3c^2)(16N_e - v^2\eta)}$ | $\frac{4[-10(1 + f) + 24c^3(3 + f) - 4c^2(23 + 11f) - c(117 + 19f)]}{5c(-8 - 2c + 3c^2)(1 + f)(16N_e - v^2\eta)}$ |
| 10 | $\frac{9 - 18c + 10c^2}{180cN_e - 90c^2N_e}$ | $\frac{25(-13 - 42c - 66c^2 + 40c^3)}{26c(-10 - 3c + 4c^2)(25N_e - v^2\eta)}$ | $\frac{25[-13(1 + f) + 40c^3(3 + f) - 6c(34 + 7f) - 2c^2(73 + 33f)]}{26c(-10 - 3c + 4c^2)(1 + f)(25N_e - v^2\eta)}$ |

Where $\eta = \frac{2(v-2)(v-1)}{v^2}$ for the MS mating system, or $\eta = \frac{4(v-1)^2}{v^2}$ for the ME/DR/DH mating systems. Each expression
should be added $1/n$ in order to consider the sampling variance.

**Table S6.**
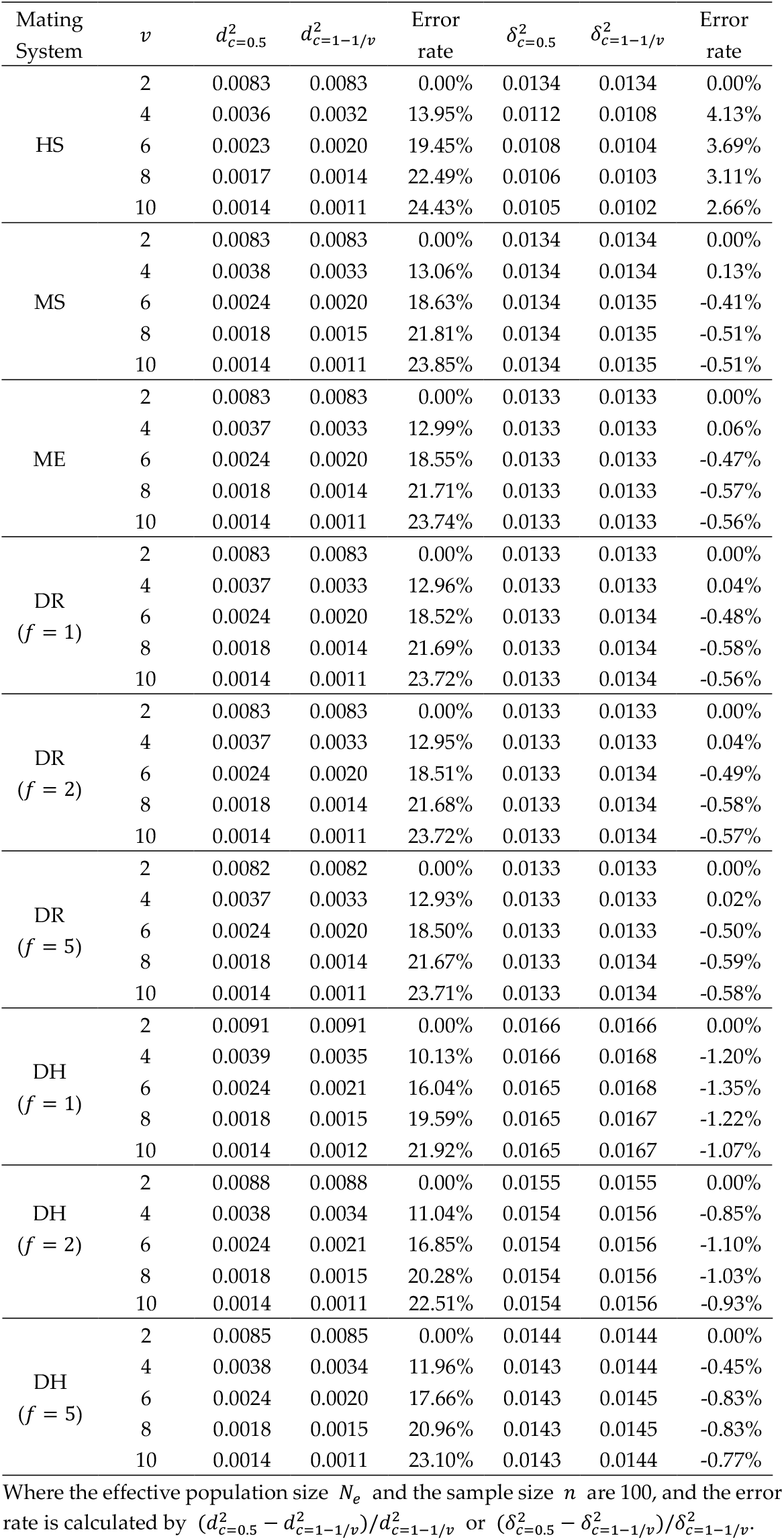
Exact *d*^2^ and *δ*^2^

### Supplementary Figures

**Figure S1.**
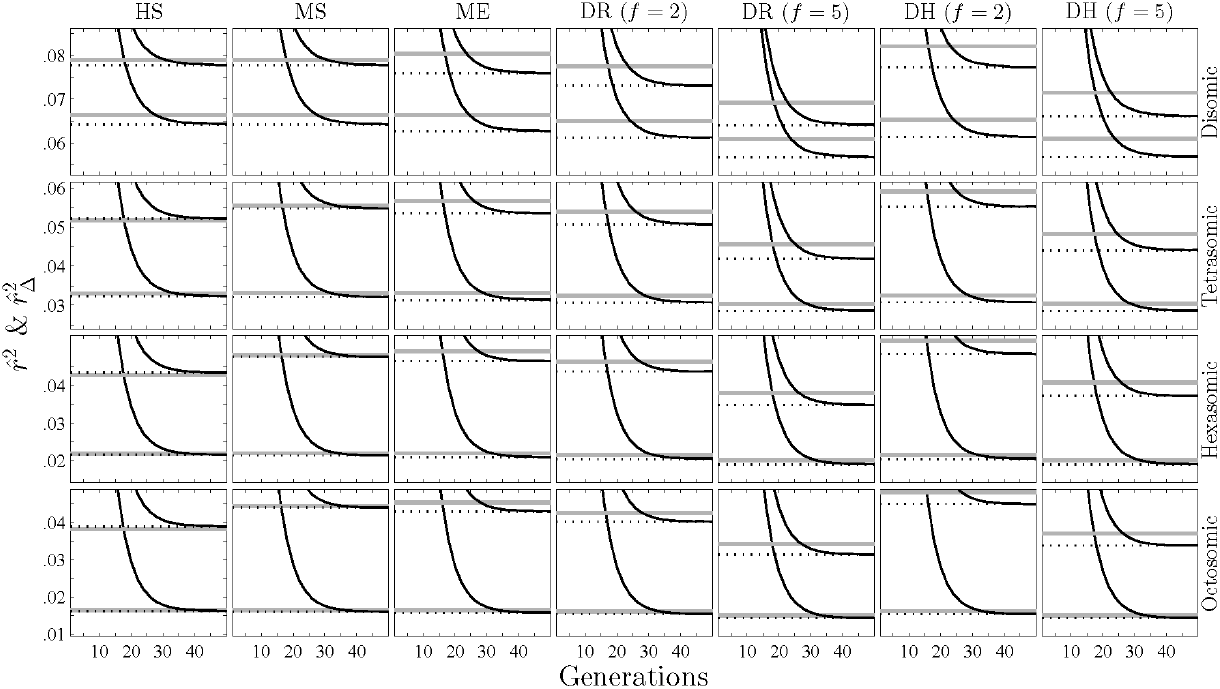
The behaviors of 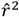 and 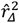 for various mating systems (set *N_e_* = 40, *v* = 2, 4, 6 or 8, *L* = 200 and *c* = 0.1; for DR and DH, also set *f* = 2 or 5). Each column shows the results under a different mating system. Each row shows the results under a different ploidy level. Solid gray lines denote the approximate *d*^2^ or *δ*^2^, dotted gray lines denote of the exact *d*^2^ or *δ*^2^, and solid lines denote 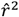 and 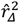, where the lines denoting *δ*^2^ (or 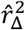) are above those denoting *d*^2^ (or 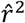) for each situation.

**Figure S2.**
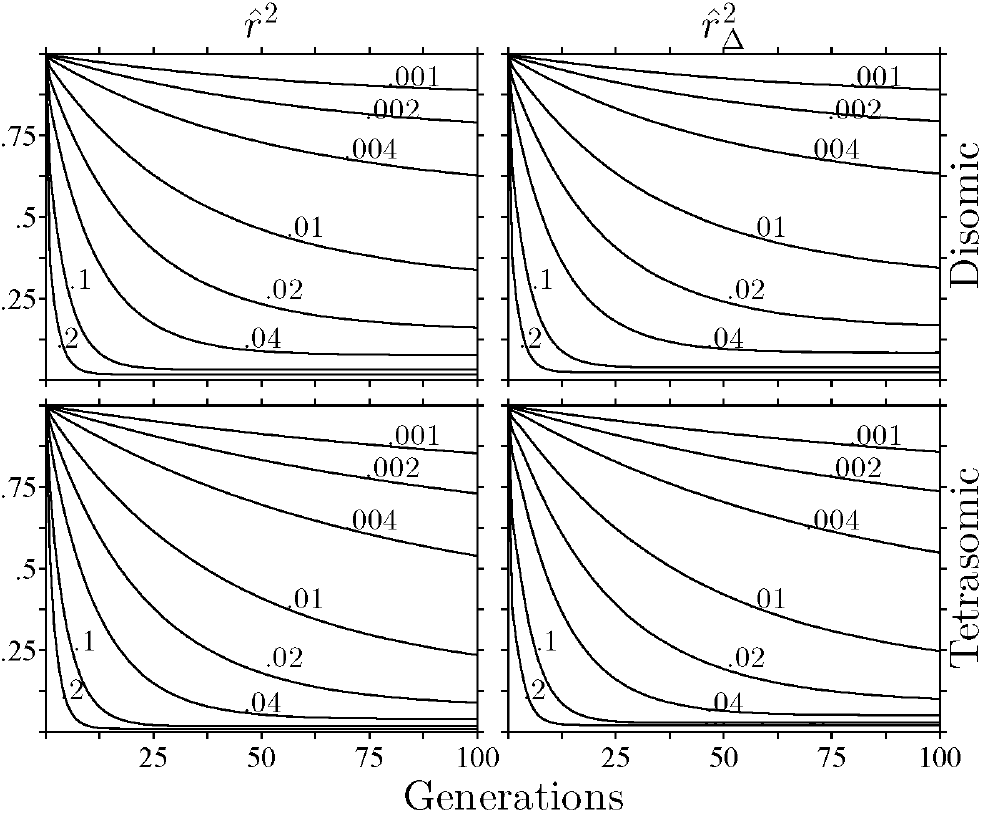
The behaviors of 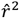 and 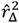 for the MS mating system under different recombination rates (set *N_e_* =80 and *L* = 200). The number above a line represents a recombination rate.

**Figure S3.**
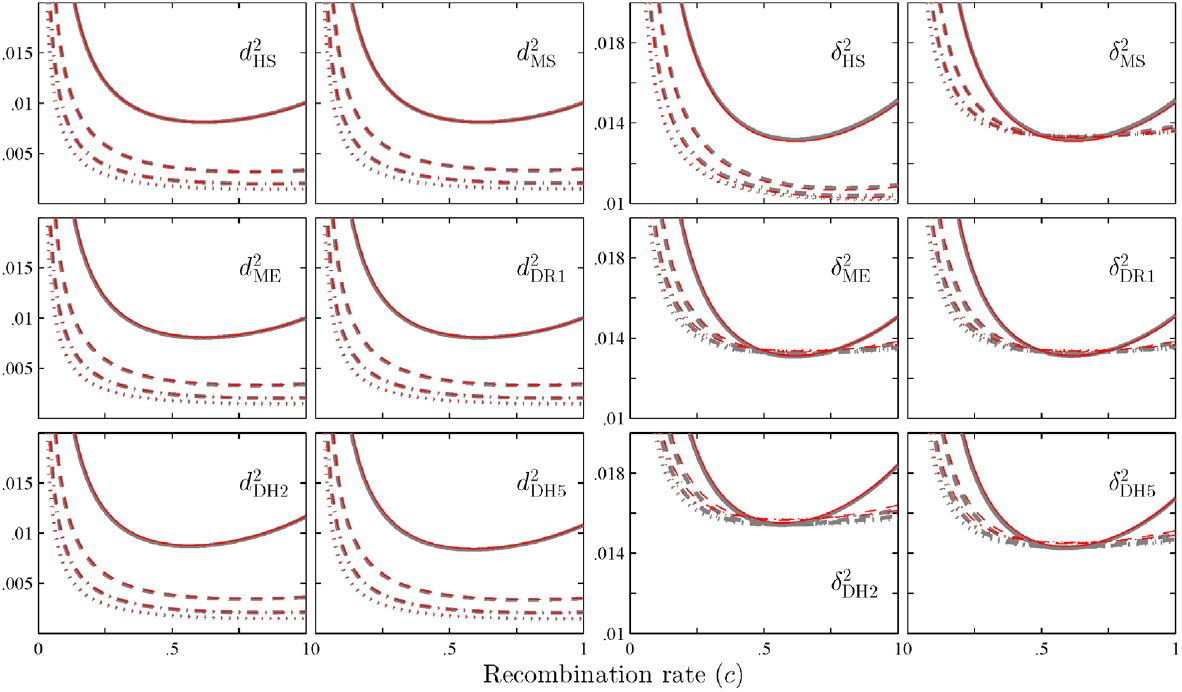
The relationship between *d*^2^ (or *δ*^2^) and the recombination rate *c* for various mating systems (set *N_e_* = 100 and *n* = 100). The line styles are the same as those in Figure 2. Each subscript of *d*^2^ or *δ*^2^ denotes a mating system, e.g. the subscript DR1 is the DR mating system with *f* = 1.

**Figure S4.**
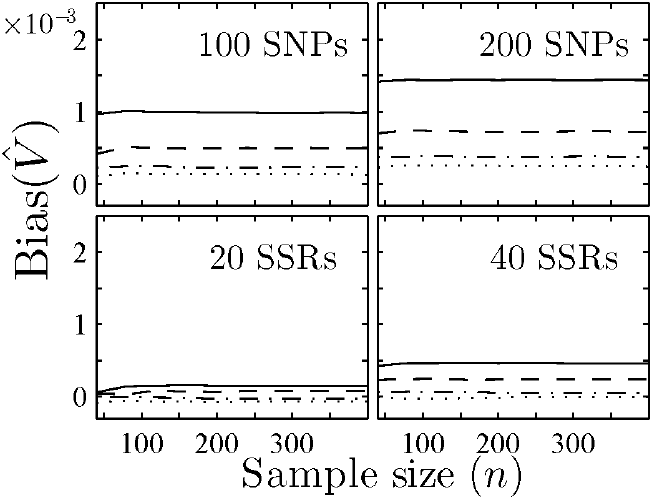
The relationship between the bias of 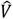 and the sample size *n*. The figure layout and the line style are the same as those in Figure 3.

## Literature Cited

Brown, A. H. D., M. W. Feldman and E. Nevo, 1980 Multilocus structure of natural populations of *Hordeum spontaneum*. Genetics 96: 523–536.

Burow, M. D., C. E. Simpson, J. L. Starr and A. H. Paterson, 2001 Transmission genetics of chromatin from a synthetic amphidiploid to cultivated peanut (*Arachis hypogaea* L.): broadening the gene pool of a monophyletic polyploid species. Genetics 159: 823.

Butruille, D. V., and L. S. Boiteux, 2000 Selection–mutation balance in polysomic tetraploids: Impact of double reduction and gametophytic selection on the frequency and subchromosomal localization of deleterious mutations. Proceedings of the National Academy of Sciences 97: 6608–6613.

Cockerham, C. C., and B. S. Weir, 1977 Digenic descent measures for finite populations. Genetics Research 30: 121–147.

Devlin, B., and N. Risch, 1995 A comparison of linkage disequilibrium measures for fine-scale mapping. Genomics 29: 311–322.

England, P. R., J.-M. Cornuet, P. Berthier, D. A. Tallmon and G. Luikart, 2006 Estimating effective population size from linkage disequilibrium: severe bias in small samples. Conservation Genetics 7: 303.

Hästbacka, J., A. de la Chapelle, I. Kaitila, P. Sistonen, A. Weaver et al., 1992 Linkage disequilibrium mapping in isolated founder populations: diastrophic dysplasia in Finland. 2: 204.

Hayes, B. J., P. M. Visscher, H. C. McPartlan and M. E. Goddard, 2003 Novel multilocus measure of linkage disequilibrium to estimate past effective population size. Genome Research 13: 635–643.

Hill, W. G., 1974 Disequilibrium among several linked neutral genes in finite population I. Mean changes in disequilibrium. Theoretical Population Biology 5: 366–392.

Hill, W. G., 1975 Linkage disequilibrium among multiple neutral alleles produced by mutation in finite population. Theoretical Population Biology 8: 117–126.

Hill, W. G., 1981 Estimation of effective population size from data on linkage disequilibrium. Genetics Research 38: 209–216.

Hill, W. G., and A. Robertson, 1968 Linkage disequilibrium in finite populations. Theoretical Applied Genetics 38: 226–231.

Hill, W. G., and B. S. Weir, 1994 Maximum-likelihood estimation of gene location by linkage disequilibrium. American Journal of Human Genetics 54: 705.

Hosking, L. K., P. R. Boyd, C. F. Xu, M. Nissum, K. Cantone et al., 2002 Linkage disequilibrium mapping identifies a 390 kb region associated with CYP2D6 poor drug metabolising activity. The Pharmacogenomics Journal 2: 165.

Huang, K., D. W. Dunn, K. Ritland and B. G. Li, 2020 polygene: Population genetics analyses for autopolyploids based on allelic phenotypes. Methods in Ecology and Evolution 11: 448–456.

Jorde, L. B., 1995 Linkage disequilibrium as a gene-mapping tool. American Journal of Human Genetics 56: 11.

Lewontin, R. C., 1964 The interaction of selection and linkage. I. General considerations; heterotic models. Genetics 49: 49.

Maruyama, T., 1982 Stochastic integrals and their application to population genetics, pp. 151–166 in Molecular Evolution, Protein Polymorphism and the Neutral Theory, edited by M. Kimura. Japan Scientific Societies press, Tokyo.

Nei, M., 1987 Molecular evolutionary genetics. Columbia university press.

Ohta, T., 1980 Linkage disequilibrium between amino acid sites in immunoglobulin genes and other multigene families. Genetics Research 36: 181–197.

Ohta, T., and M. Kimura, 1969 Linkage disequilibrium at steady state determined by random genetic drift and recurrent mutation. Genetics 63: 229.

Otto, S. P., 2007 The evolutionary consequences of polyploidy. Cell 131: 452–462.

Rieger, R., A. Michaelis and M. M. Green, 1968 A glossary of genetics and cytogenetics: Classical and molecular. Springer-Verlag, New York.

Sattler, M. C., C. R. Carvalho and W. R. Clarindo, 2016 The polyploidy and its key role in plant breeding. Planta 243: 281–296.

Slatkin, M., 2008 Linkage disequilibrium-understanding the evolutionary past and mapping the medical future. Nature Reviews Genetics 9: 477.

Sved, J. A., E. C. Cameron and A. S. Gilchrist, 2013 Estimating effective population size from linkage disequilibrium between unlinked loci: theory and application to fruit fly outbreak populations. PLoS oNE 8: e69078.

Sved, J. A., and M. W. Feldman, 1973 Correlation and probability methods for one and two loci. Theoretical Population Biology 4: 129–132.

Udall, J. A., and J. F. Wendel, 2006 Polyploidy and crop improvement. Crop Science 46: S-3–S-14.

Waples, R. S., T. Antao and G. Luikart, 2014 Effects of overlapping generations on linkage disequilibrium estimates of effective population size. Genetics 197: 769–780.

Weir, B. S., 1979 Inferences about linkage disequilibrium. Biometrics: 235–254.

Weir, B. S., and C. C. Cockerham, 1979 Estimation of linkage disequilibrium in randomly mating populations. Heredity 42: 105.

Weir, B. S., and W. G. Hill, 1980 Effect of mating structure on variation in linkage disequilibrium. Genetics 95: 477–488.

